## Supplementary material for "Patterns and drivers of diatom diversity and abundance in the global ocean": Suppl Tables and Figs

Pierella Karlusich, et al.

**Table S1**: Estimated ASV richness for V4 and V9 datasets using multiple biodiversity estimators.

| Marker | Observed ASV count | Chao1 estimator | Chao1 standard error | First-order Jackknife estimator | First-order Jackknife standard error | Second-order Jackknife estimator | Bootstrap estimator | Bootstrap standard error | Total number of observations |
|---|---|---|---|---|---|---|---|---|---|
| V9 | 34243424 | 3424.899 | 1.040739 | 3441.981 | 4.238084 | 3280.522 | 3472.61 | 25.21701 | 931 |
| V4 | 3429 | 3429.878 | 1.043834 | 3443.985 | 4.791743 | 3331.35 | 3473.272 | 16.97236 | 969 |

**Table S2:** Partial least square analyses to explore the correlations between diatom relative abundance and Shannon index with the physicochemical context

| | *Diatom relative abundance* | | *Diatom Shannon* | |
|---|---|---|---|---|
| | *VIP score* | *Standard coefficient* | *VIP score* | *Standard coefficient* |
| *Temperature* | **1.6266765279** | -0.18715260 | **1.5764379** | 0.14211408 |
| *NH4+/DIN* | 0.7449139294 | -0.06770999 | 0.7251692 | 0.04639623 |
| *NH4+* | 0.0004108426 | -0.04073561 | 0.3672286 | 0.04230855 |
| *NO2-* | 0.0186188635 | -0.05158923 | 0.4473117 | 0.05296705 |
| *NO3-* | **1.1938164317** | 0.10643676 | **1.1647857** | -0.07219932 |
| *Fe* | 0.0761344935 | 0.04829419 | 0.3701818 | -0.04770001 |
| *Si* | **1.0040927560** | 0.09568034 | 0.9730387 | -0.06711979 |
| *PO43-* | 0.8829044972 | 0.05801240 | 0.9078752 | -0.03189882 |
| *Chlorophyll a* | **1.1668409466** | 0.12469494 | **1.1261470** | -0.09202232 |
| *Absolute latitude* | **1.4894446409** | 0.18268462 | **1.4563720** | -0.14187917 |

**Table S3:** Partial least square analyses to explore the correlations between diatom relative abundance and biotic factors

| | *Diatom relative abundance* | |
|---|---|---|
| | *VIP score* | *Standard coefficient* |
| *Prochlorococcus* | **2.0589894** | -0.236453116 |
| *Synechococcus* | 0.7559680 | -0.087359284 |
| *Copepoda* | **1.4889173** | 0.151032888 |
| *Centroheliozoa* | **1.5517588** | -0.059139034 |
| *Choanoflagellida* | **1.7055238** | 0.195750218 |
| *Chrysophyceae* | **1.4418286** | -0.007121123 |
| *Dictyochophyceae* | **2.3466947** | -0.082131732 |
| *Nassellaria* | **1.4064072** | -0.162014670 |
| *Phaeodarea* | 0.4485323 | -0.051816101 |
| *Spumellaria* | 0.9581452 | -0.094281817 |
| *Rhizaria* | 0.5690121 | -0.055823984 |

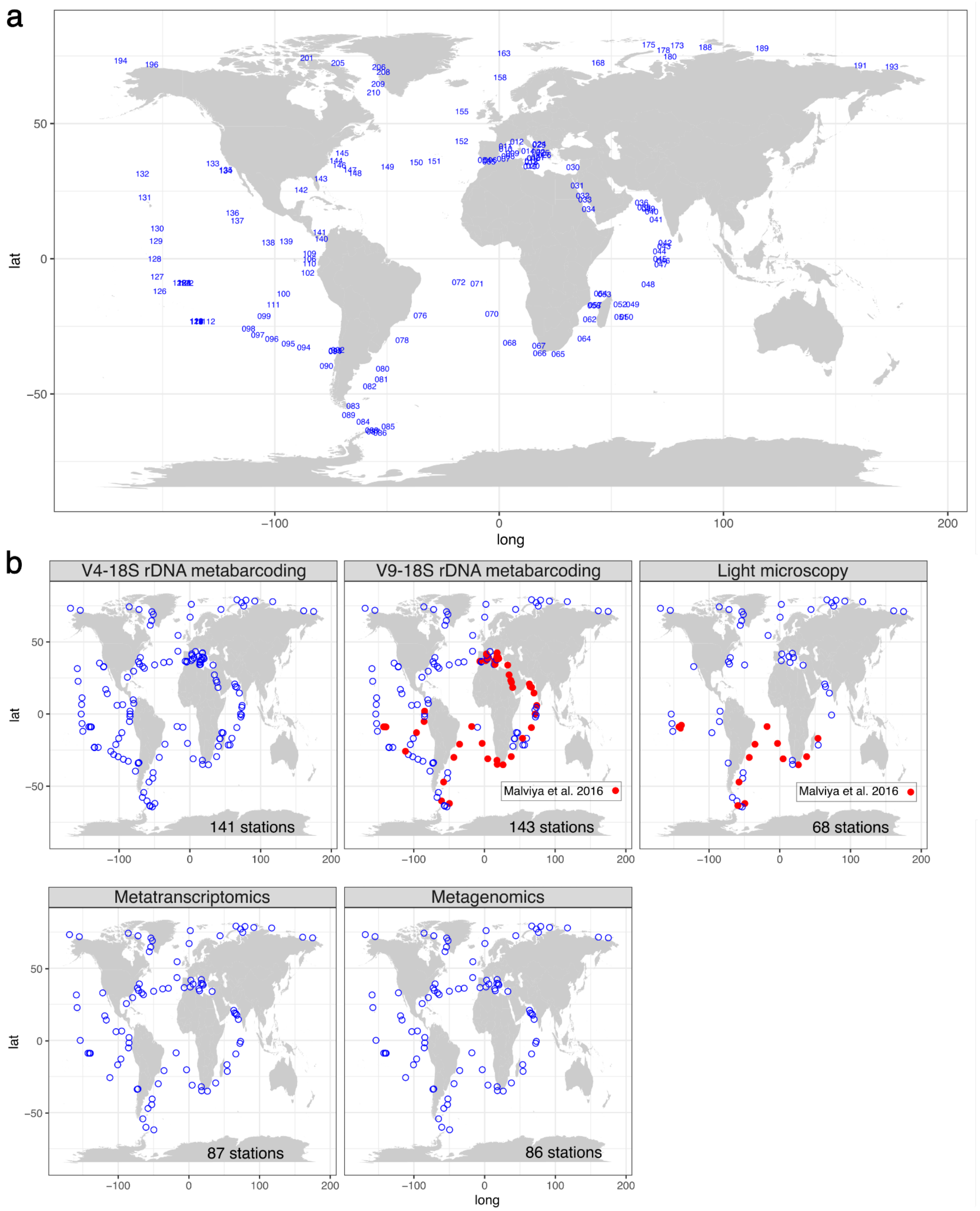


**Figure S1**: Biogeographical coverage of the *Tara* Oceans datasets used in the current work. a) Sampling station labels. b) Biogeography of the datasets. The total number of stations are indicated. The sampling stations reported by Malviya et al. 2016 *PNAS* 113 (11):E1516-E1525 are marked in red: 43 sampling stations for the V9-based survey and 15 stations for the microscopy survey.

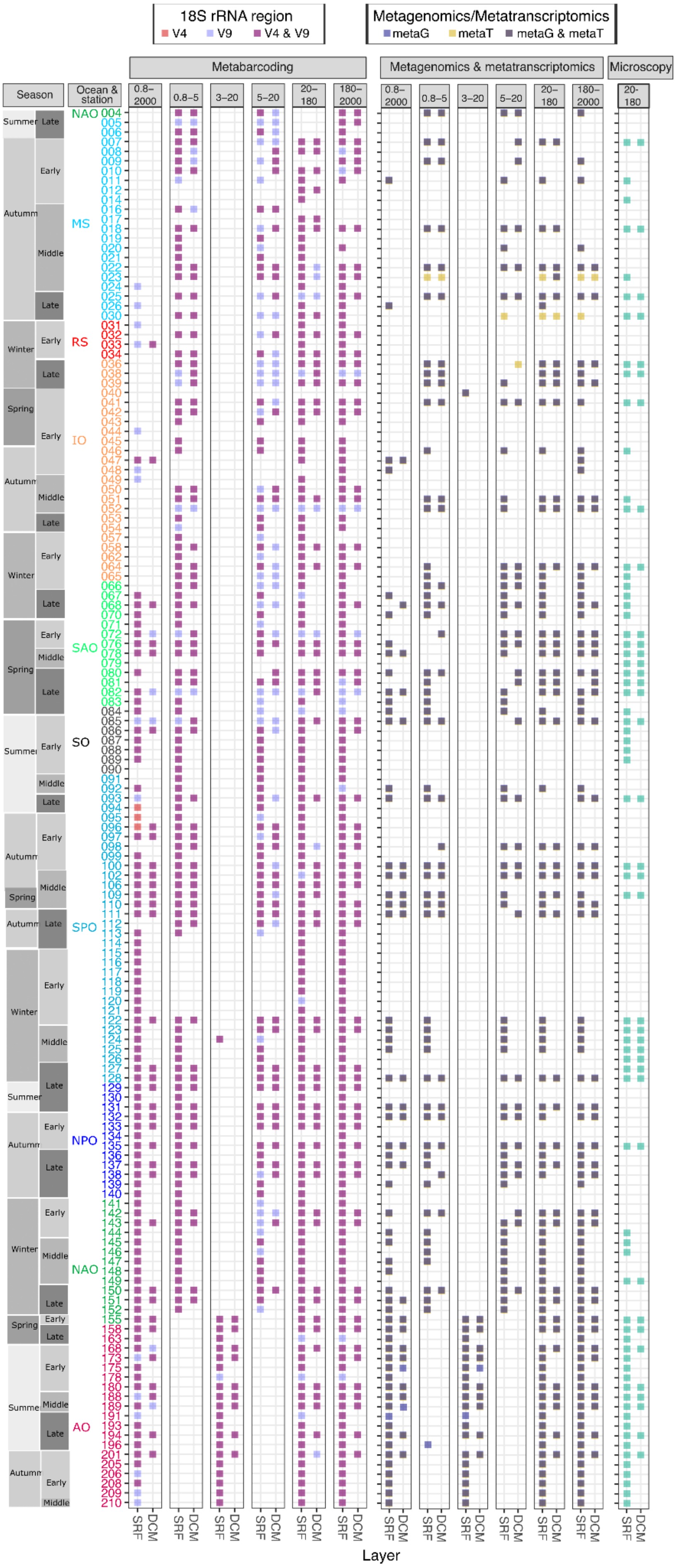

18S rRNA region
V4
V9
V4 & V9
Metagenomics/Metatranscriptomics
metaG
metaT
metaG & metaT
Metabarcoding
Metagenomics & metatranscriptomics
Microscopy
Season
Ocean & station
0.8–2000
0.8–5
3–20
5–20
20–180
180–2000
20-180
Summer
Autumn
Winter
Spring
Early
Middle
Late
NAO
MS
RS
IO
SAO
SO
SPO
NPO
AO
SRF
DCM
Layer

**Figure S2**: Samples and methods used in this study. The current analysis of global diversity and abundance of diatoms was carried out across 145 *Tara* Oceans stations where samples were taken for metabarcoding and/or metagenomics and/or metatranscriptomics and/or optical microscopy. The analyzed samples are indicated as filled boxes, with color according to the method. Two distinct depth layers were sampled: surface (SUR; 5 m) and deep chlorophyll maximum (DCM; 17-180 m). The data from the bottom of the mixed layer was collected when no DCM was observed (stations TARA_123, TARA_124, TARA_125, TARA_152 and TARA_153). Plankton communities were fractionated into four size classes: piconanoplankton (0.8 to 5 µm or 0.8 to 2000 µm), nanoplankton (5 to 20 µm or 3 to 20 µm), microplankton (20 to 180 µm), and mesoplankton (180 to 2000 µm). Season and moment of the season (early, middle, late) are displayed to the left of the panel. Station labels are coloured according to the ocean region: IO, Indian Ocean; MS, Mediterranean Sea; NAO, North Atlantic Ocean; RS, Red Sea; SAO, South Atlantic Ocean; SO, Southern Ocean; SPO, South Pacific Ocean.

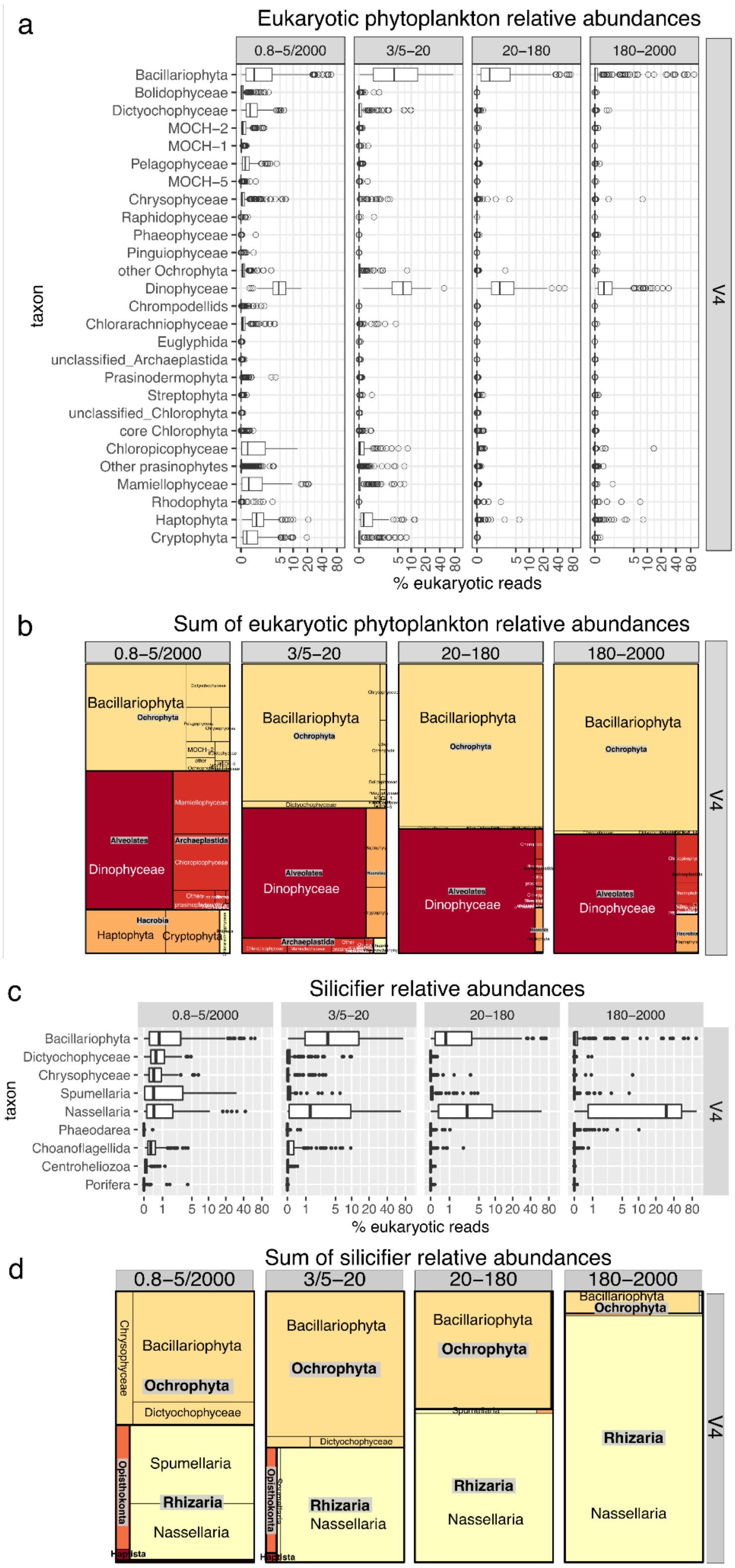


**Figure S3:** Contribution of diatoms to eukaryotic phytoplankton (a-b) and silicifiers (c-d) in surface waters of the global ocean using V4 metabarcoding data obtained from different size-fractions. a-b) Phytoplankton. We focused exclusively on the phytoplankton signal of these data sets, including dinoflagellates and chrysophytes though we acknowledge there are uncertainties in assigning the capacity for photosynthesis in these groups. a) Relative abundances (log scale). Each point is a size-fractionated sample. b) Sum of normalized reads in the overall dataset. c-d) Silicifiers. The equivalent plots for the V9 marker are displayed in Fig 1.

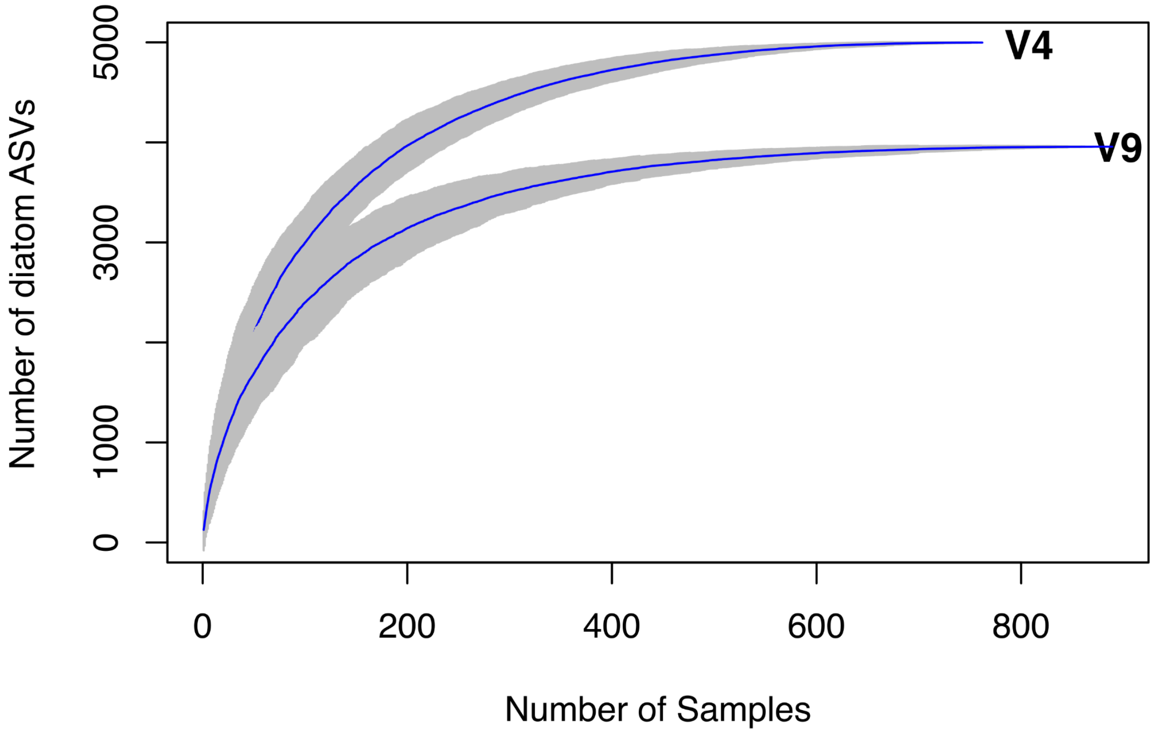


**Figure S4:** Accumulation curves for V4 and V9 18S rRNA gene metabarcoding assigned to diatoms.

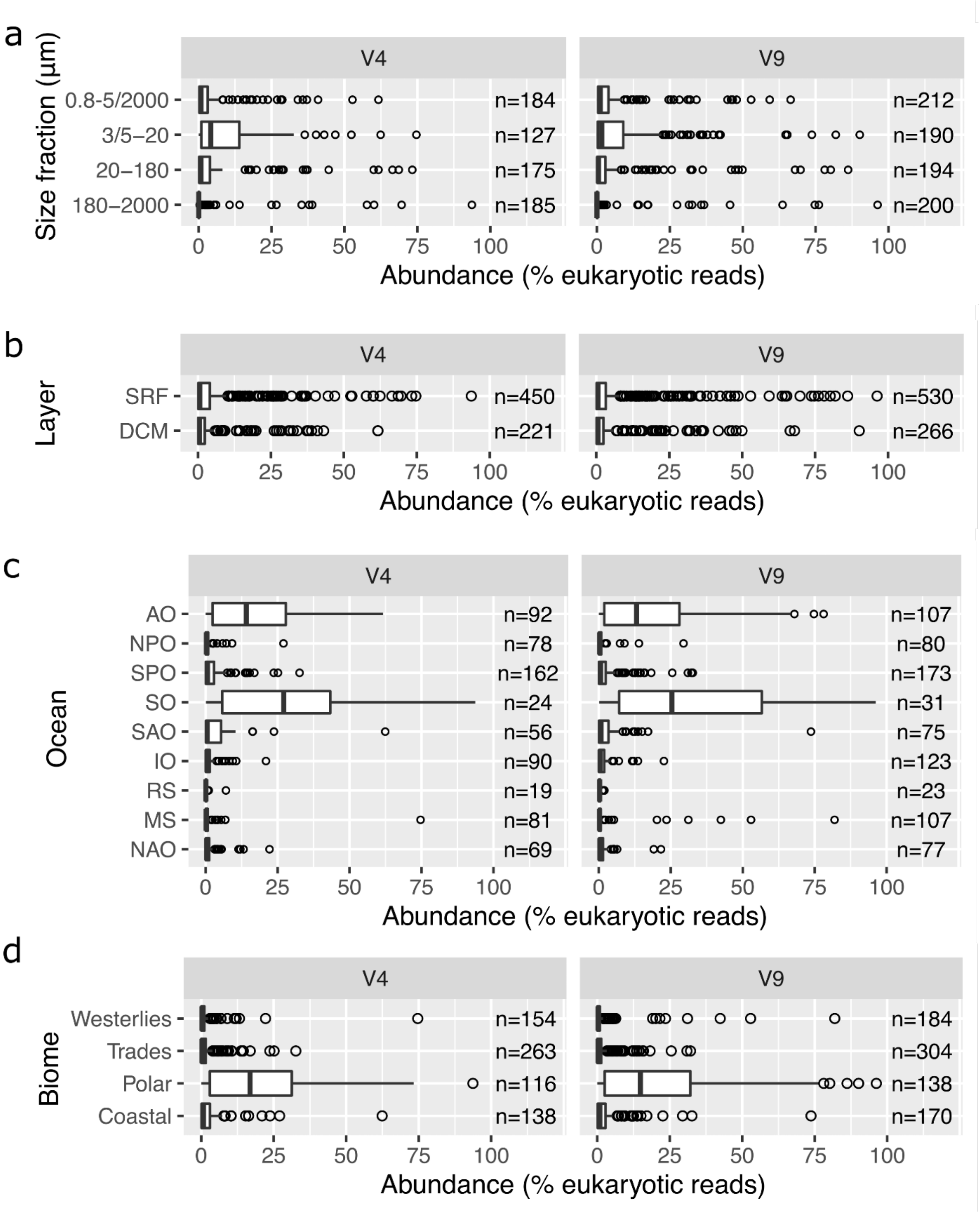


**Figure S5:** Distribution of diatom relative abundances in epipelagic seawater samples from the V9 and V4 metabarcoding data across a) size fraction. b) water layers (SRF, surface; DCM, deep chlorophyll maximum), c) ocean basins (MS, Mediterranean Sea, IO, Indian Ocean, SAO, South Atlantic Ocean, SO, Southern Ocean, SPO, South Pacific Ocean, NPO, North Pacific Ocean, NAO, North Atlantic Ocean, AO, Arctic Ocean), d) biomes.

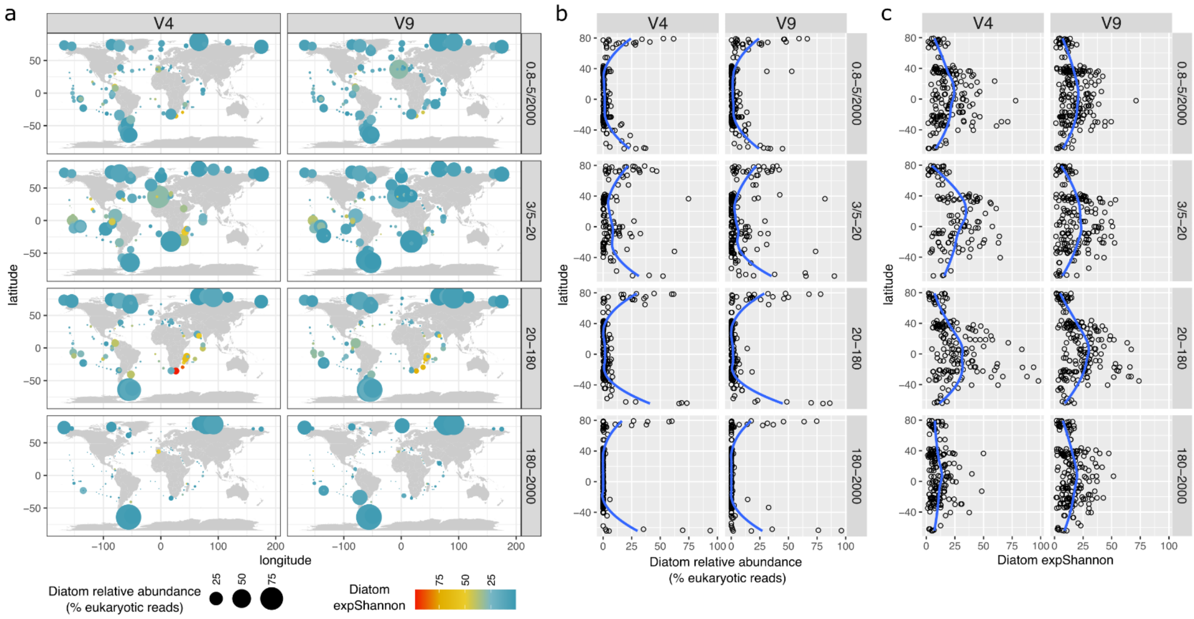


**Figure S6:** Comparison between V4 and V9 metabarcoding for the global macroecological patterns of diatom relative abundance and diversity in surface waters using data obtained from different size-fractions. a) Distribution maps. b) Latitudinal gradient for relative abundance. c) Latitudinal gradient for the exponentiated Shannon Diversity Index. The blue lines correspond to Loess smoothings.

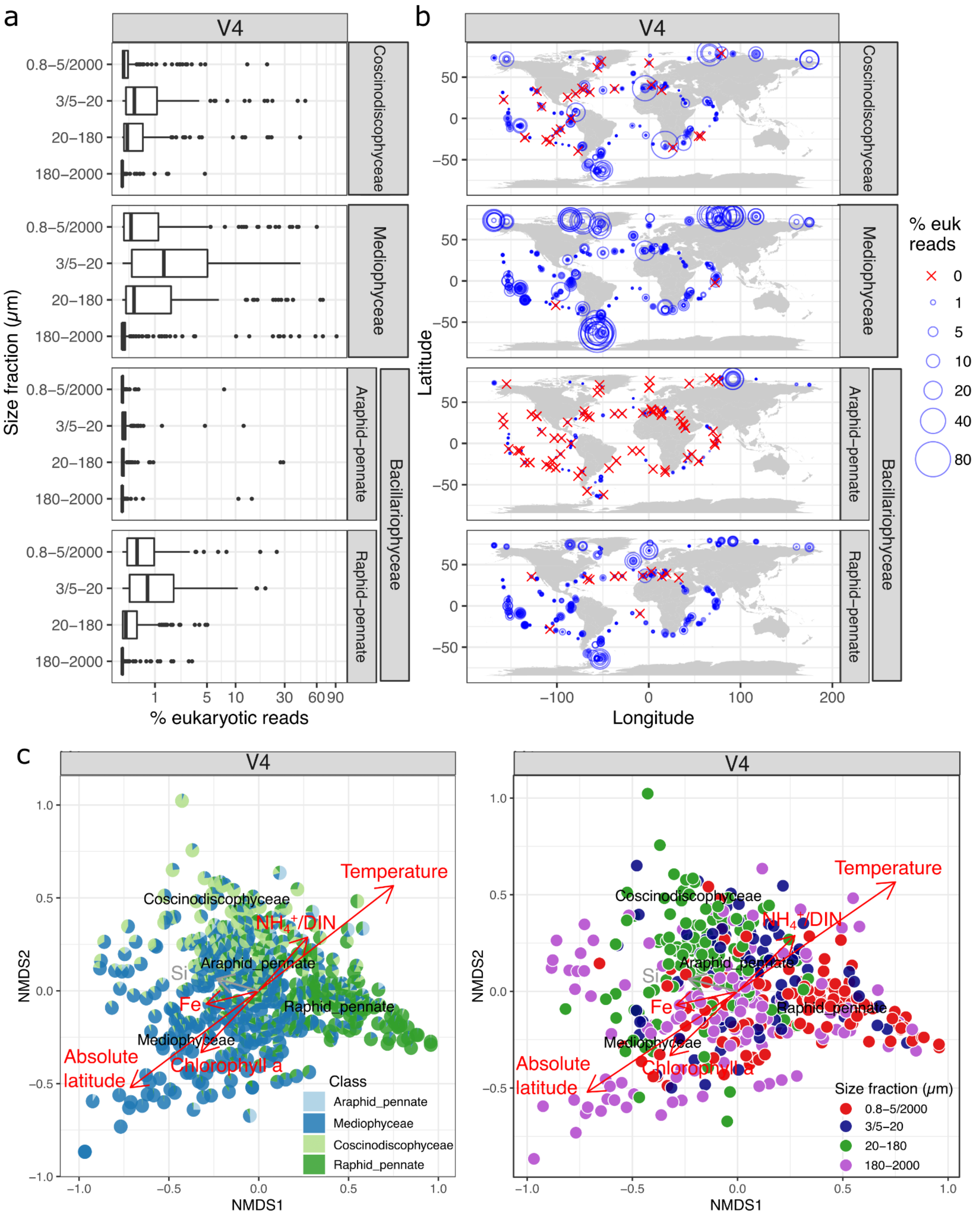


**Figure S7:** Abundance patterns of diatom classes based on V4 marker. a) Relative abundances (log scale). Each point is a size-fractionated sample. b) Biogeography. c) NMDS analysis of stations according to Bray–Curtis distance. Fitted statistically significant physico-chemical parameters are displayed in black lines when adjusted P value < 0.05 (temperature, iron, chlorophyll a concentration, absolute latitude, ratio of ammonium to total dissolved inorganic nitrogen) and in gray lines when between 0.1 and 0.05 (silicon). Each pie chart (left) or each circle (right) is a size-fractionated sample. NMDS stress value: 0.1197578. The equivalent figure for V9 is displayed in **Figure 4**.

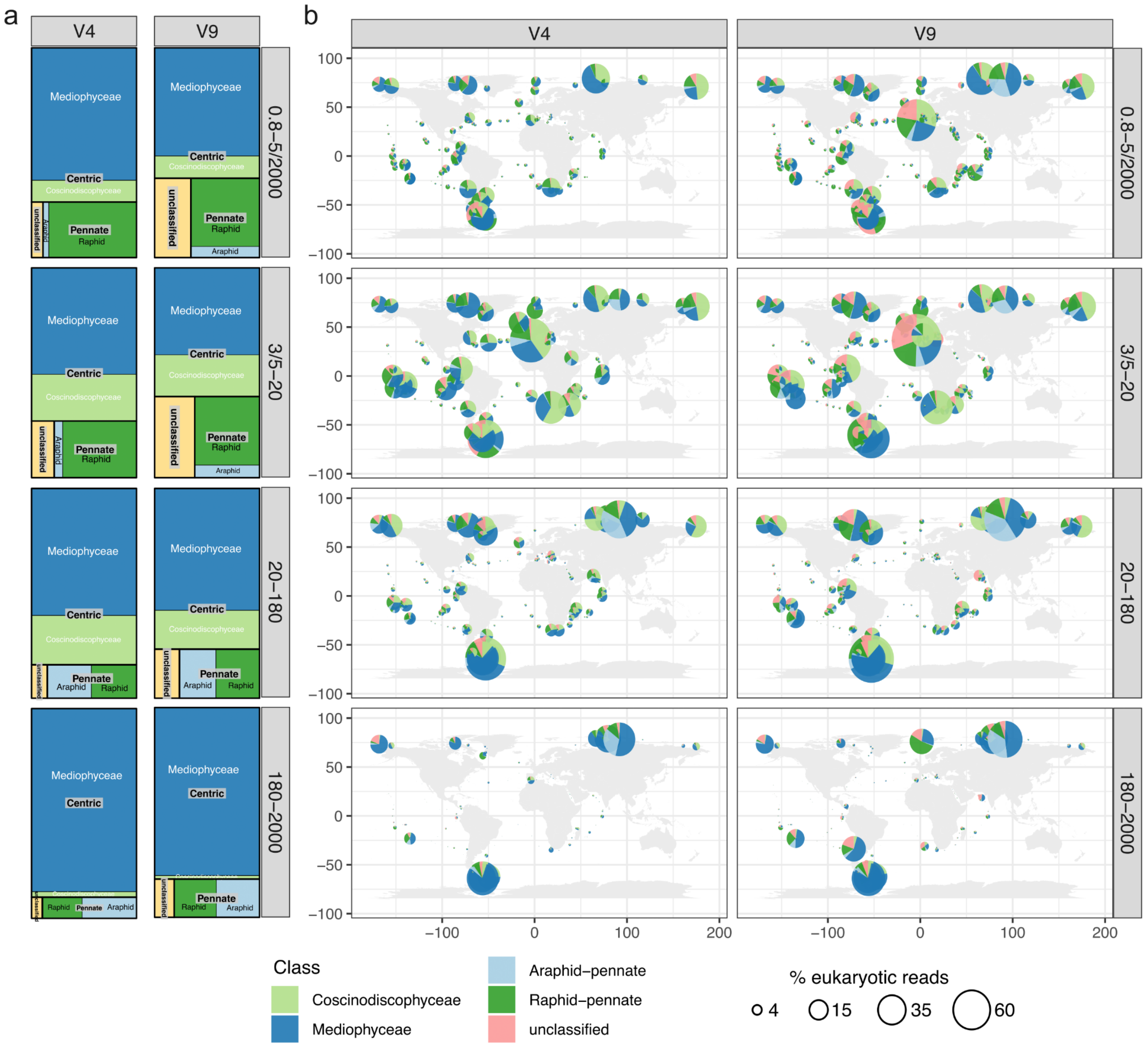


**Figure S8:** Relative contribution of the different diatom classes. a) Sum of normalized reads for V4 and V9 markers in the overall dataset. b) Biogeography. The maps separated by diatom classes are shown in **Figure S9**.

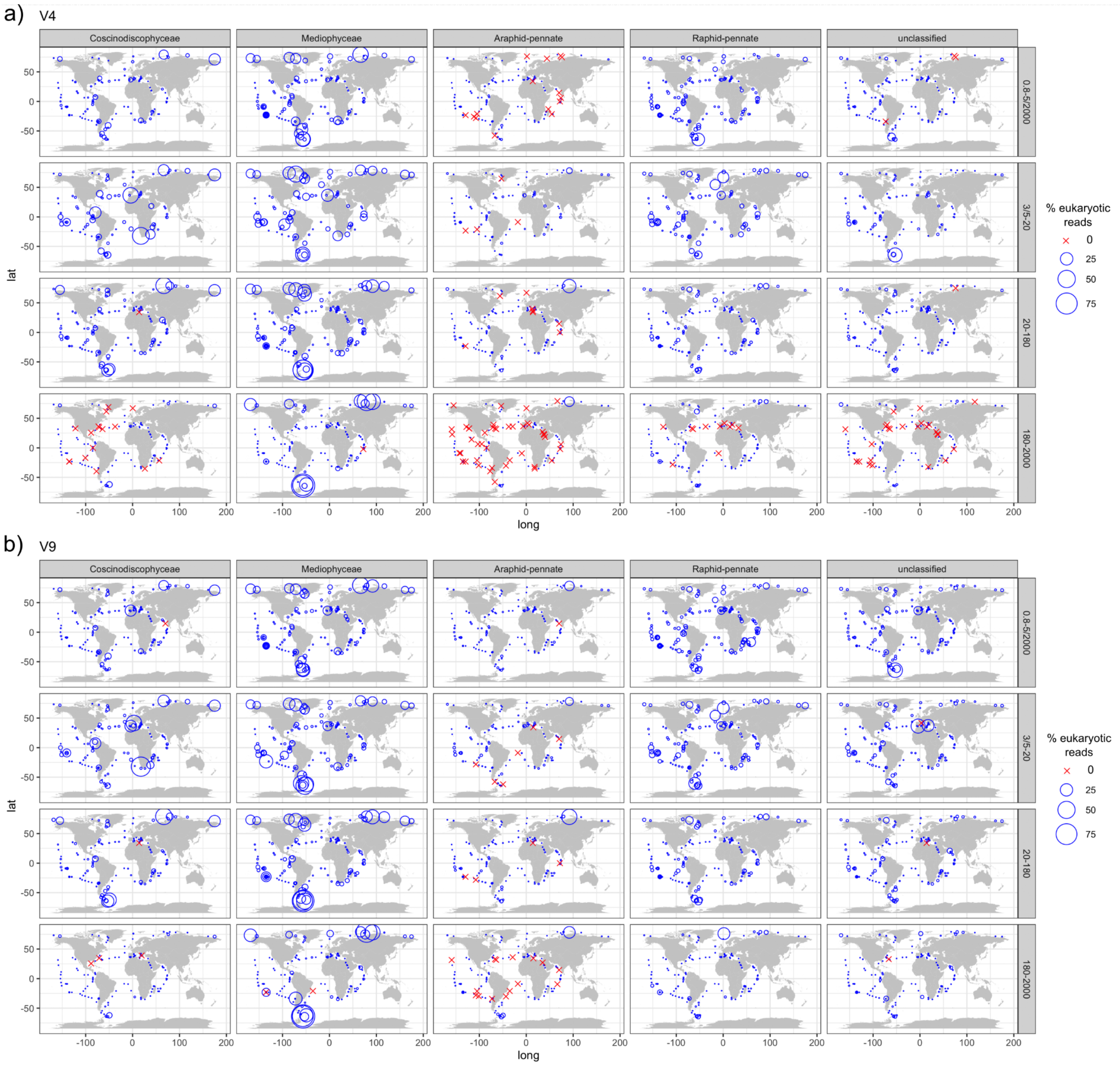


**Figure S9:** Biogeography of diatom classes in surface waters using V4 (a) and V9 (b) metabarcoding data obtained from different size-fractionated samples. The bubble sizes vary according to the percentage of diatoms among eukaryotes.

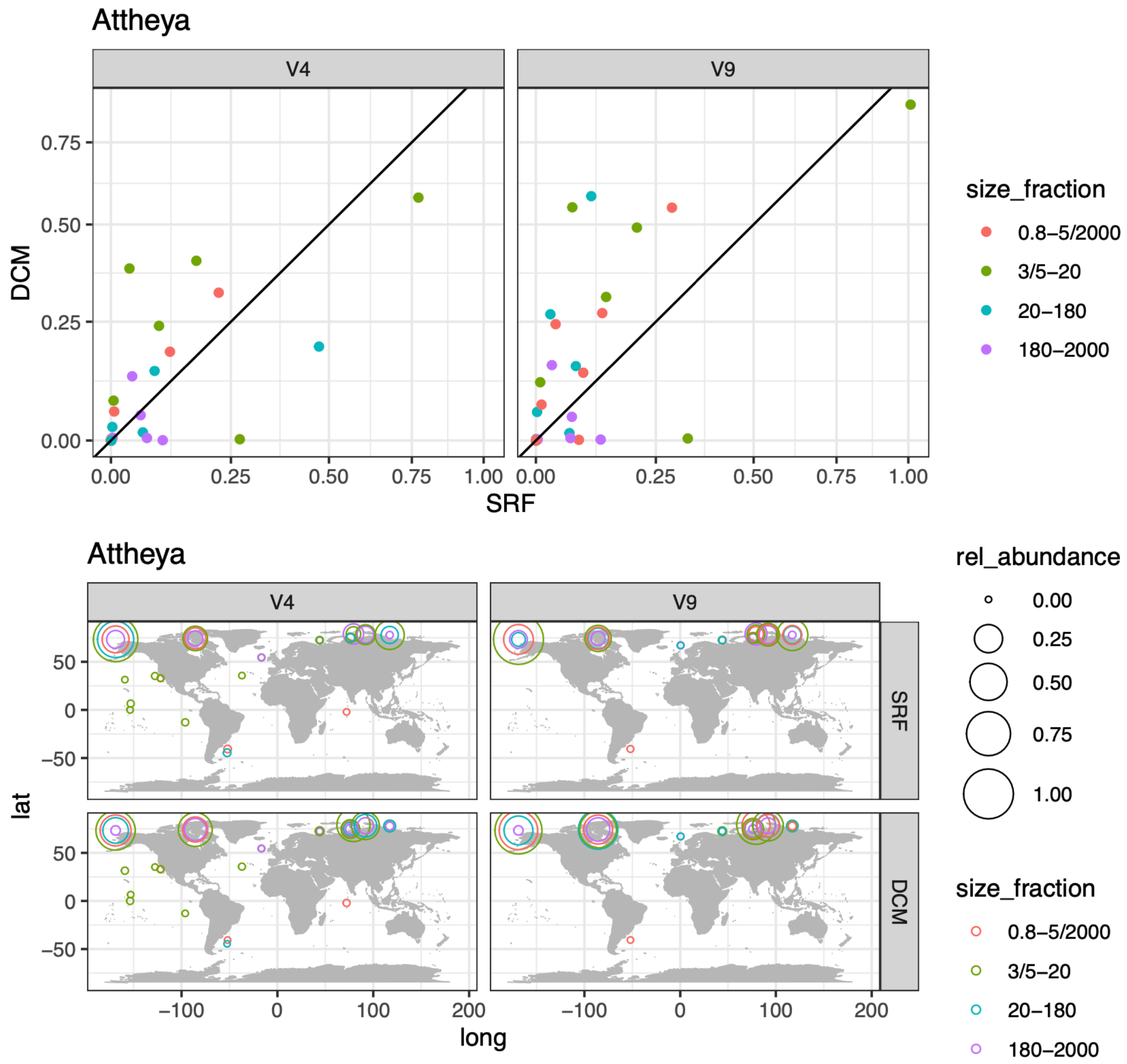


**Figure S10:** Environmental distribution of the *Attheya* genus. a) Depth distribution. The scatter plots compare the relative abundances between surface (5 m) and deep chlorophyll maximum (DCM; 17–180 m). Axes are in the same scale and the diagonal line corresponds to a 1:1 slope. b) Biogeography. The bubble sizes vary according to the percentage of diatoms among eukaryotes. The data corresponds to V4 (left panel) and V9 (right panel) metabarcoding data obtained from different size-fractionated samples (indicated in colors).

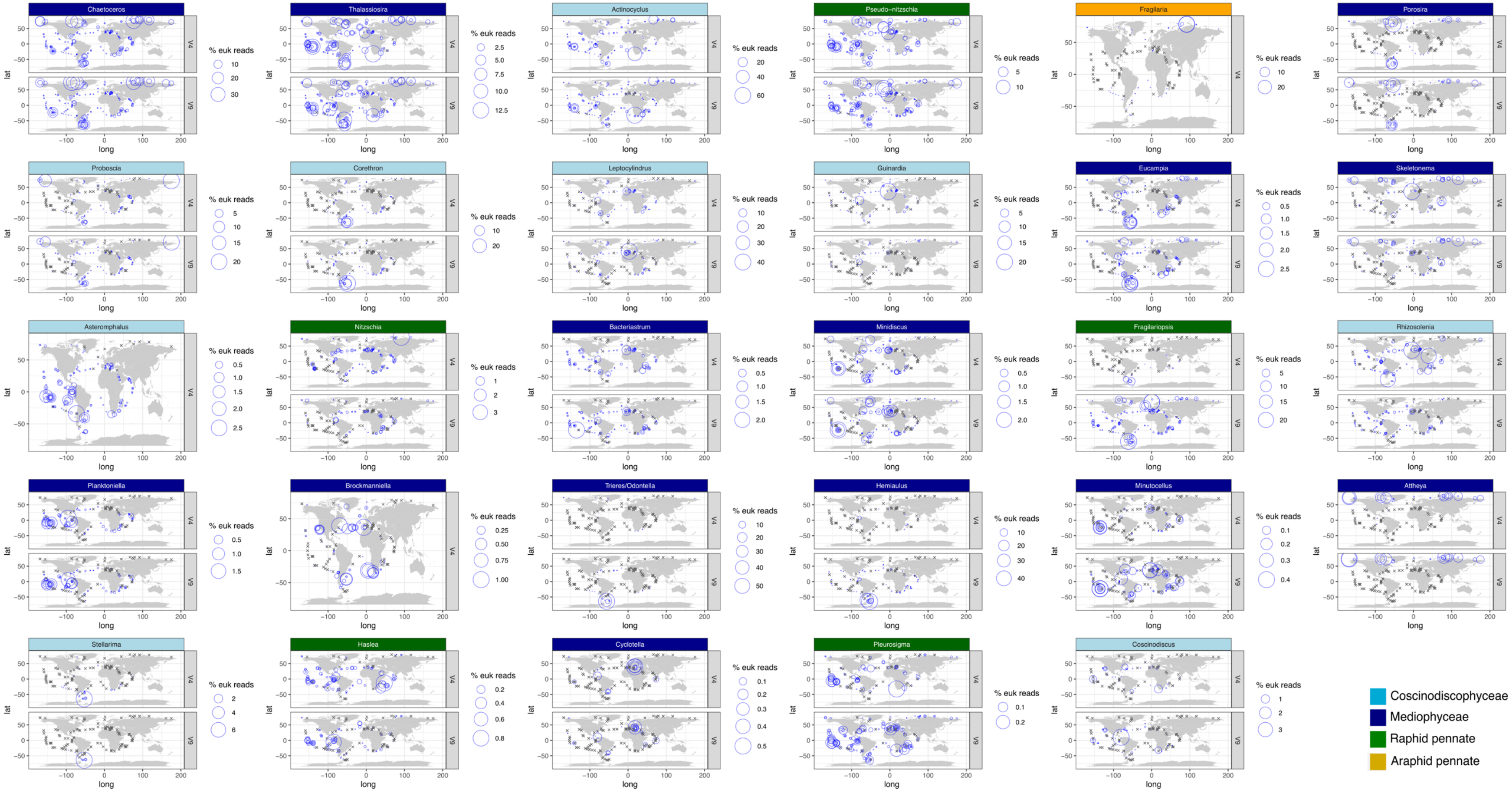

Chaetoceros
Thalassiosira
Actinocyclus
Pseudo−nitzschia
Fragilaria
Porosira
Proboscia
Corethron
Leptocylindrus
Guinardia
Eucampia
Skeletonema
Asteromphalus
Nitzschia
Bacteriastrum
Minidiscus
Fragilariopsis
Rhizosolenia
Planktoniella
Brockmanniella
Trieres/Odontella
Hemiaulus
Minutocellus
Attheya
Stellarima
Haslea
Cyclotella
Pleurosigma
Coscinodiscus
% euk reads
lat
long
V4
V9
Coscinodiscophyceae
Mediophyceae
Raphid pennate
Araphid pennate

**Figure S11:** Global distribution of the 20 most abundant genera in the V9 and V4 datasets. All genera in the list of Fig 6a are displayed. This figure is analogous to Figure 7, but differentiates by marker region. Bubble areas represent the percentage of reads for each genus among eukaryotic reads at each station location, with crosses indicating absence of detection. The maps depict surface samples from the size fraction where the genus was most prevalent: 20-180 µm for *Chaetoceros*, *Fragilaria*, *Porosira*, *Proboscia*, *Corethron*, Eucampia, Asteromphalus, Planktoniella and *Trieres*/*Odontella, and Stellarima*; 3/5-20 µm for *Thalassiosira*, *Actinocyclus*, *Pseudo-nitzschia*, *Guinardia*, *Leptocylindrus*, *Fragilariopsis*, Skeletonema, Bacteriastrum, Rhizosolenia, Attheya, Haslea, Pleurosigma, Coscinodiscus, and *Hemiaulus; 0.8-5/2000 µm for Nitzschia, Minidiscus, Brockmanniella, Minutocellus, and Cyclotella*. The maps including all size fractions and covering the top 50 most abundant genera are available in Supplementary File 1.

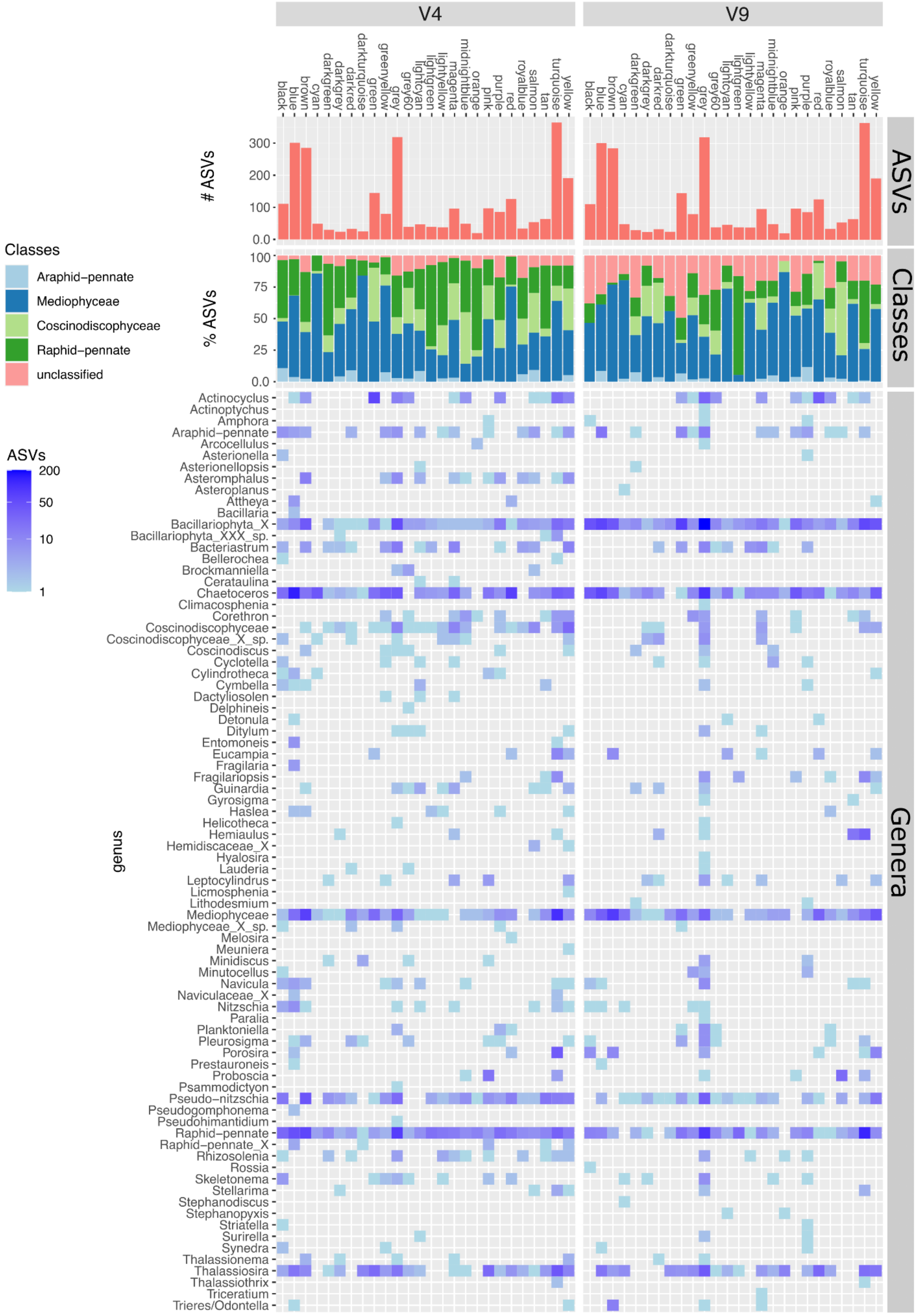


**Figure S12:** Taxonomic composition of the diatom ASV modules obtained by WGCNA.

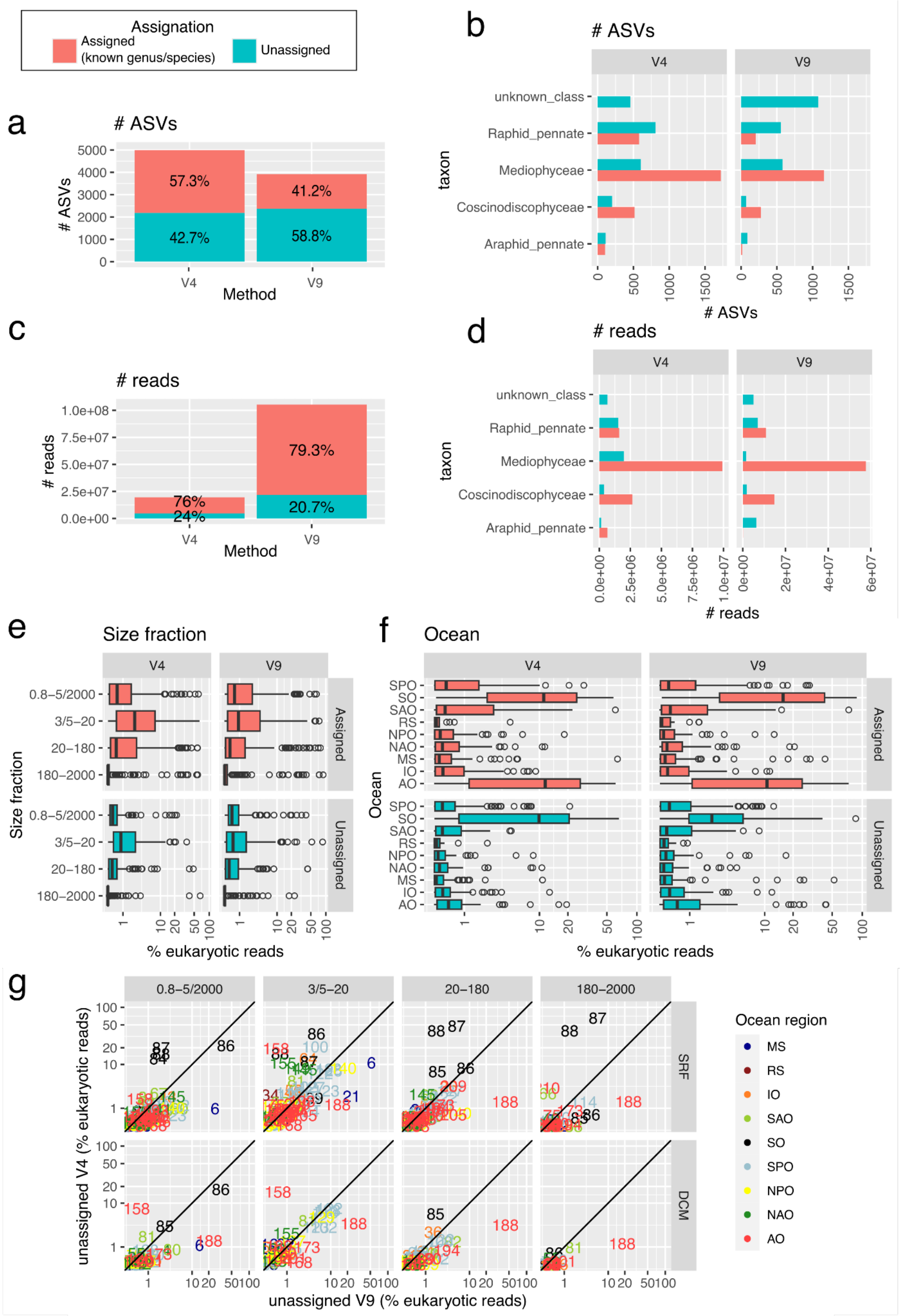

Assignation
Assigned (known genus/species)
Unassigned
a
### ASVs
57.3%
42.7%
41.2%
58.8%
Method
b
### ASVs
V4
V9
unknown_class
Raphid_pennate
Mediophyceae
Coscinodiscophyceae
Araphid_pennate
taxon
c
### reads
76%
24%
79.3%
20.7%
d
### reads
e
Size fraction
0.8–5/2000
3/5–20
20–180
180–2000
Assigned
Unassigned
% eukaryotic reads
f
Ocean
SPO
SO
SAO
RS
NPO
NAO
MS
IO
AO
g
0.8–5/2000
3/5–20
20–180
180–2000
SRF
DCM
unassigned V4 (% eukaryotic reads)
unassigned V9 (% eukaryotic reads)
Ocean region
MS
RS
IO
SAO
SO
SPO
NPO
NAO
AO

**Figure S13:** Unassigned ASVs in the *Tara* Oceans V4 and V9 metabarcoding datasets from diatoms. The ASVs considered to be unassigned were those that could not be unambiguously assigned to any diatom genus/species but could be classified only as araphid or raphid pennate, Mediophyceae, Coscinodiscophyceae, or unassigned diatom. a-b) Total number of assigned (red) and unassigned (blue) ASVs, and their class-level distribution. c-d) Total number of assigned (red) and unassigned (blue) reads, and their class-level distribution. e) Read abundances of assigned and unassigned ASVs per size class. f) Read abundances of assigned and unassigned ASVs by ocean region. g) Comparison of the read abundance of unassigned ASVs in those samples that are represented by both V4 and V9 metabarcoding. Axes are in the same logarithmic scale and the diagonal line corresponds to a 1:1 slope. Station labels are indicated, with the color according to the ocean region: MS Mediterranean Sea, IO Indian Ocean, SAO South Atlantic Ocean, SO Southern Ocean, SPO South Pacific Ocean, NPO North Pacific Ocean, NAO North Atlantic Ocean, AO Arctic Ocean. Each point in e-g panels corresponds to a size-fractionated sample.

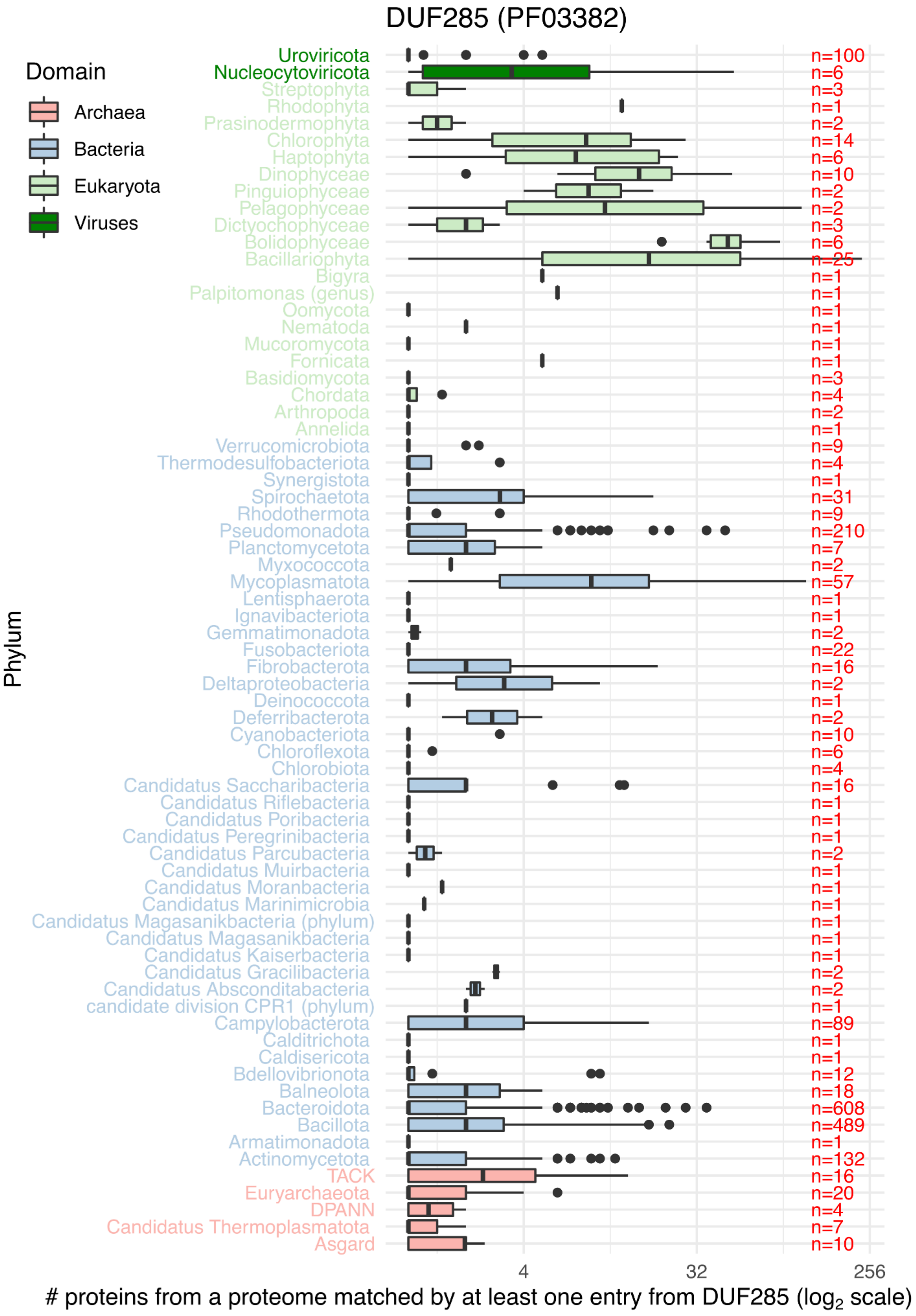


**Figure S14:** Phylogenetic distribution of DUF285 (PF03382) in reference proteomes from UniprotKB database (https://www.uniprot.org/uniprotkb). The number of analysed species is displayed in red. The x axis corresponds to the number of unique DUF285 sequences per proteome (log2 scale)

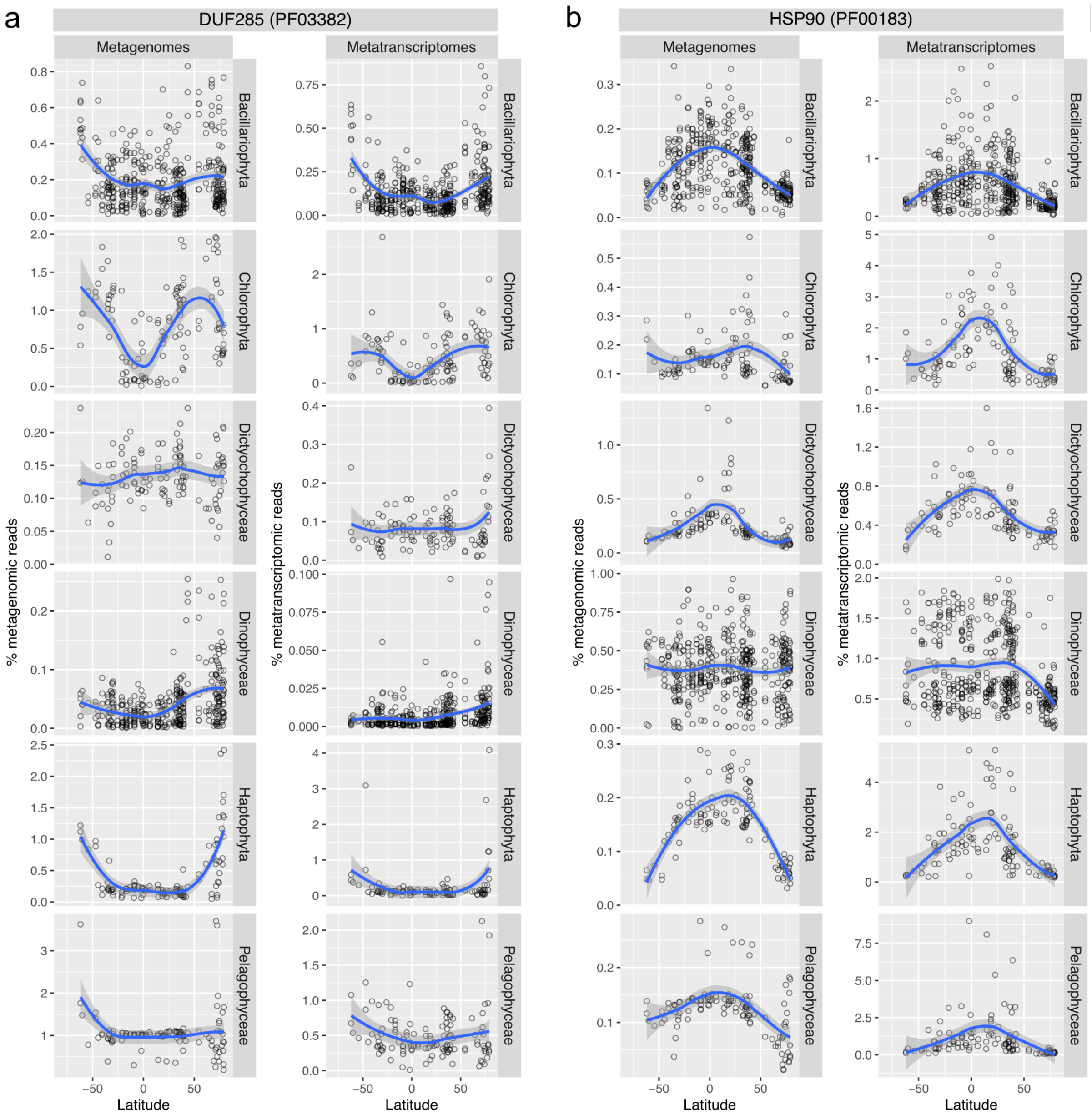


**Figure S15:** Latitudinal abundance gradient for genes and transcripts coding for Domain of Unknown Function 285 (DUF285) and Heat Shock Protein 90 (HSP90) in diatoms and other eukaryotic phytoplankton. a) DUF285 (PF03382). b) HSP90 (PF00183). The scatter plots correspond to samples from the size fraction where the taxon was most prevalent: 0.8-5/2000, 3/5-20, 20-180, and 180-2000 µm for diatoms and dinoflagellates, and 0.8-5/2000 µm for the rest. Y axis corresponds to the proportion of metagenomic or metatranscriptomic reads for the given function among the total metagenomic or metatranscriptomic reads from the corresponding taxon. Note that we were not able to discard heterotrophic species from dinoflagellates due to the small number of reference gene catalogs for this group.

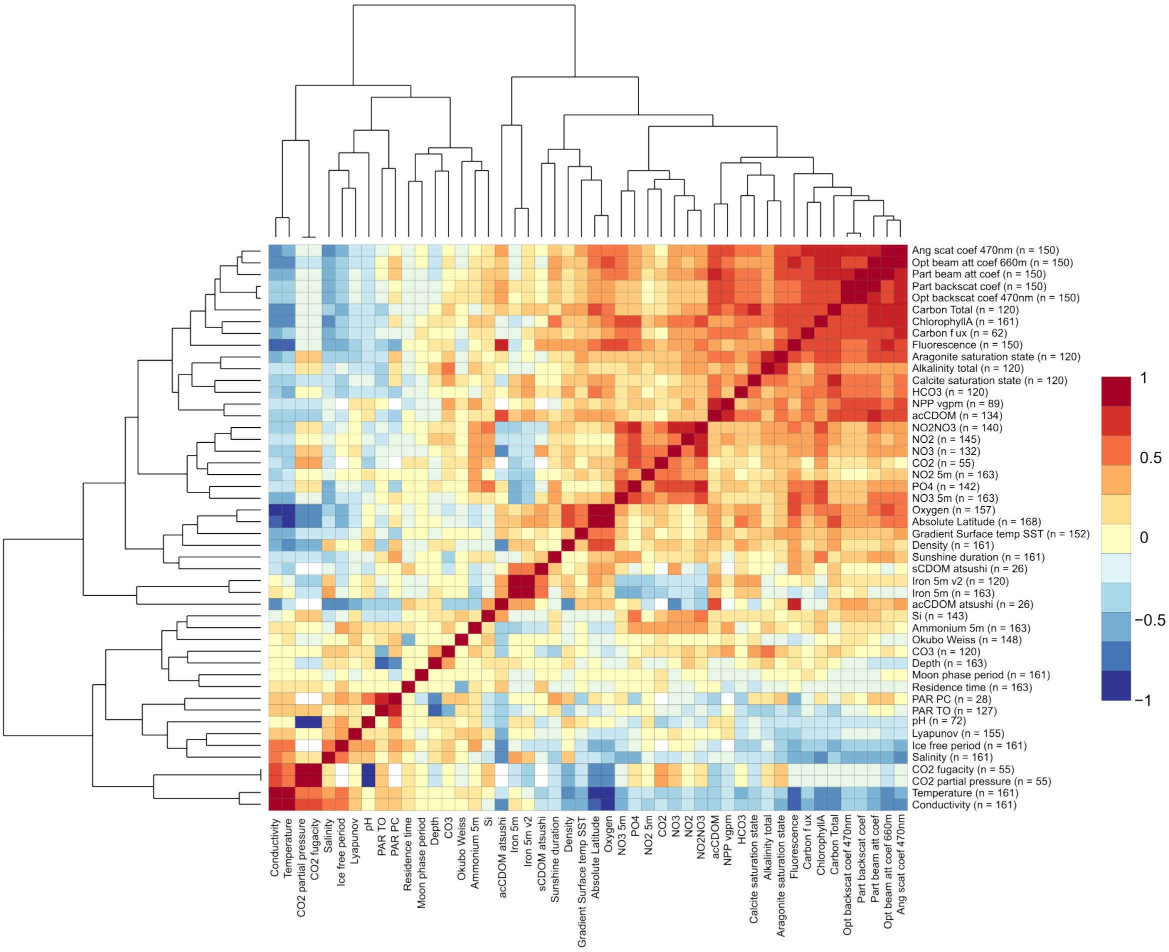


**Figure S16:** Correlation matrix for the environmental variables analyzed in the current work. Spearman rho correlation values are displayed. Sample size is indicated for each variable.
