## Supplementary File 1 for "Patterns and drivers of diatom diversity and abundance in the global ocean"

**Supplementary File 1**
**Patterns and drivers of diatom diversity and abundance in the global ocean**
**Pierella Karlusich, et al.**

Global distribution of the diatom genera detected in V4 and V9 datasets. Bubble areas are scaled to the % reads for the genus among eukaryotic reads at each station location, whereas crosses indicate absence of detection. Colors indicate the exponentiated Shannon Diversity Index. Note that some genera are only detected in V4 or V9 dataset.

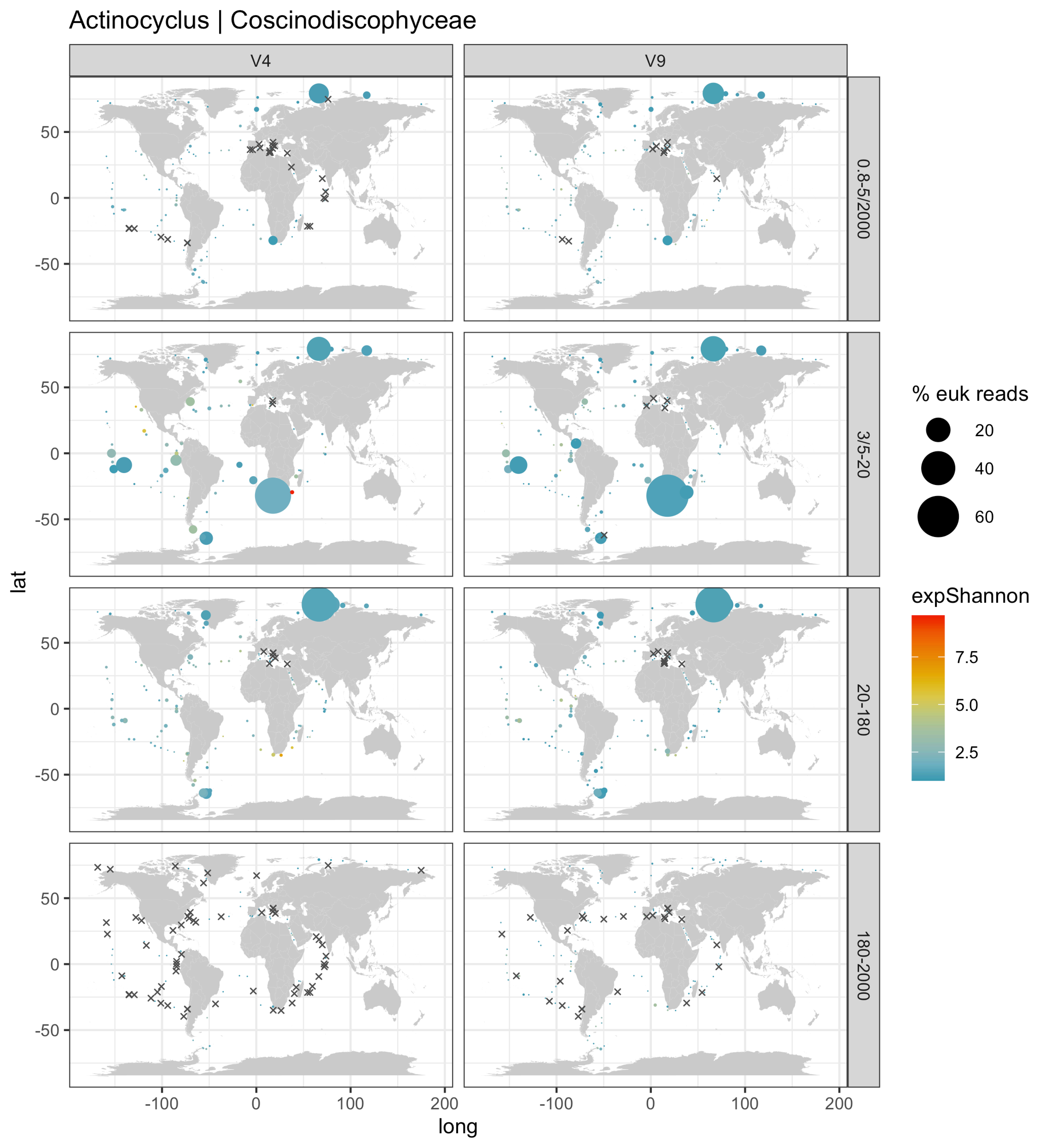

Actinocyclus | Coscinodiscophyceae
V4
V9
0.8-5/2000
3/5-20
20-180
180-2000
lat
long
50
0
-50
-100
0
100
200
% euk reads
20
40
60
expShannon
7.5
5.0
2.5

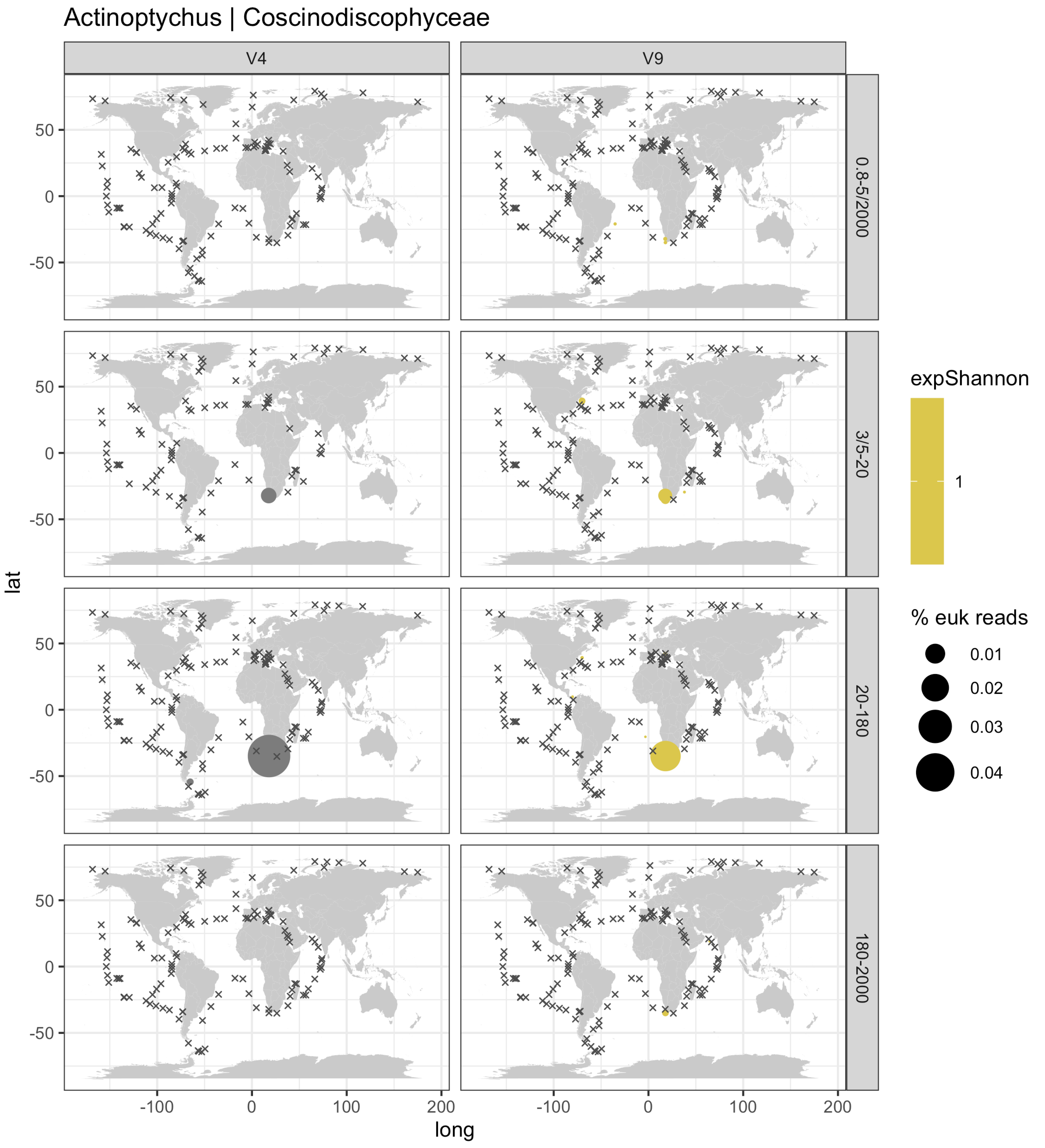

Actinoptychus | Coscinodiscophyceae
V4
V9
0.8-5/2000
3/5-20
20-180
180-2000
lat
long
50
0
-50
-100
100
200
expShannon
1
% euk reads
0.01
0.02
0.03
0.04

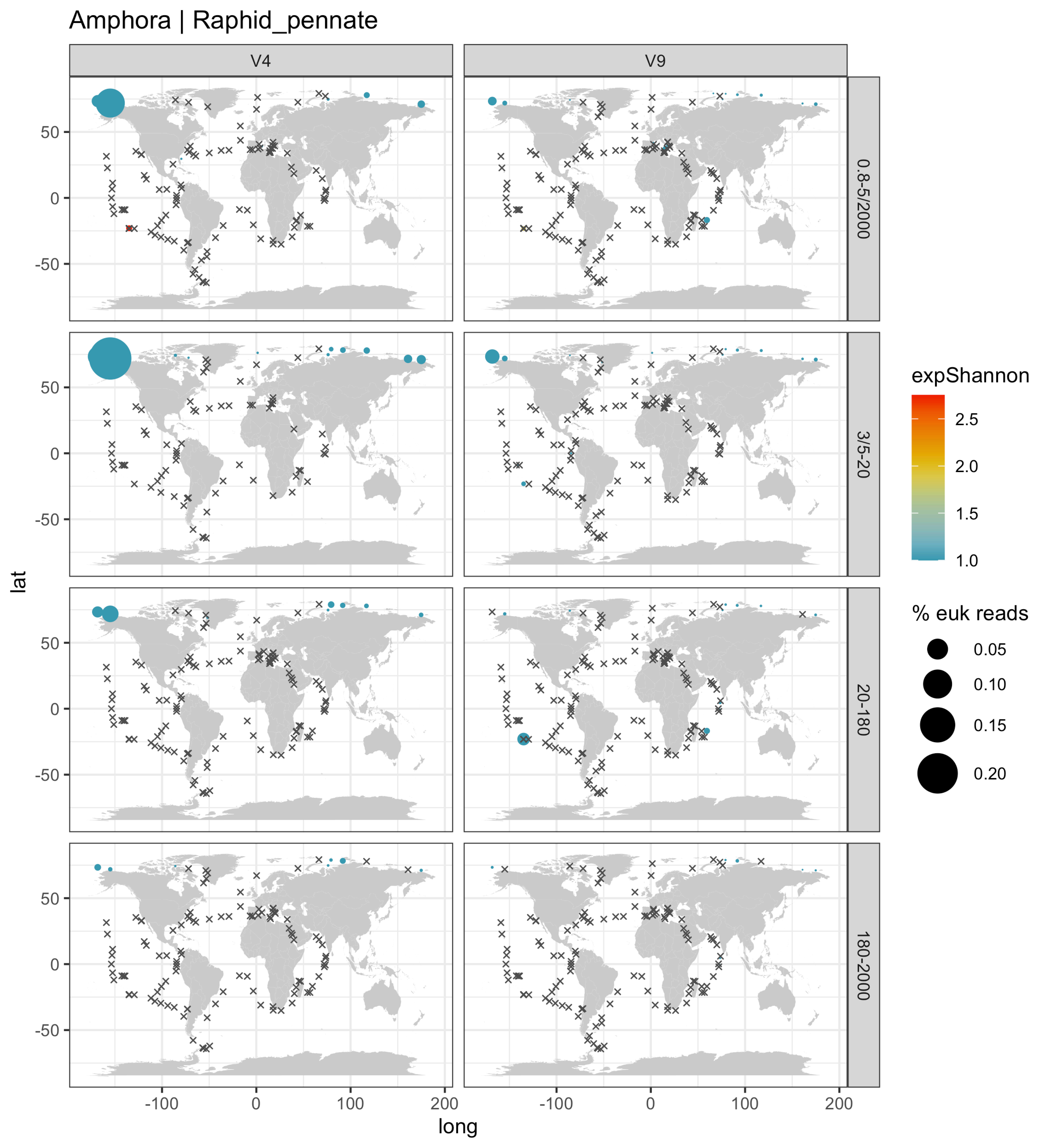

Amphora | Raphid_pennate
V4
V9
0.8-5/2000
3/5-20
20-180
180-2000
lat
long
50
0
-50
-100
100
200
expShannon
2.5
2.0
1.5
1.0
% euk reads
0.05
0.10
0.15
0.20

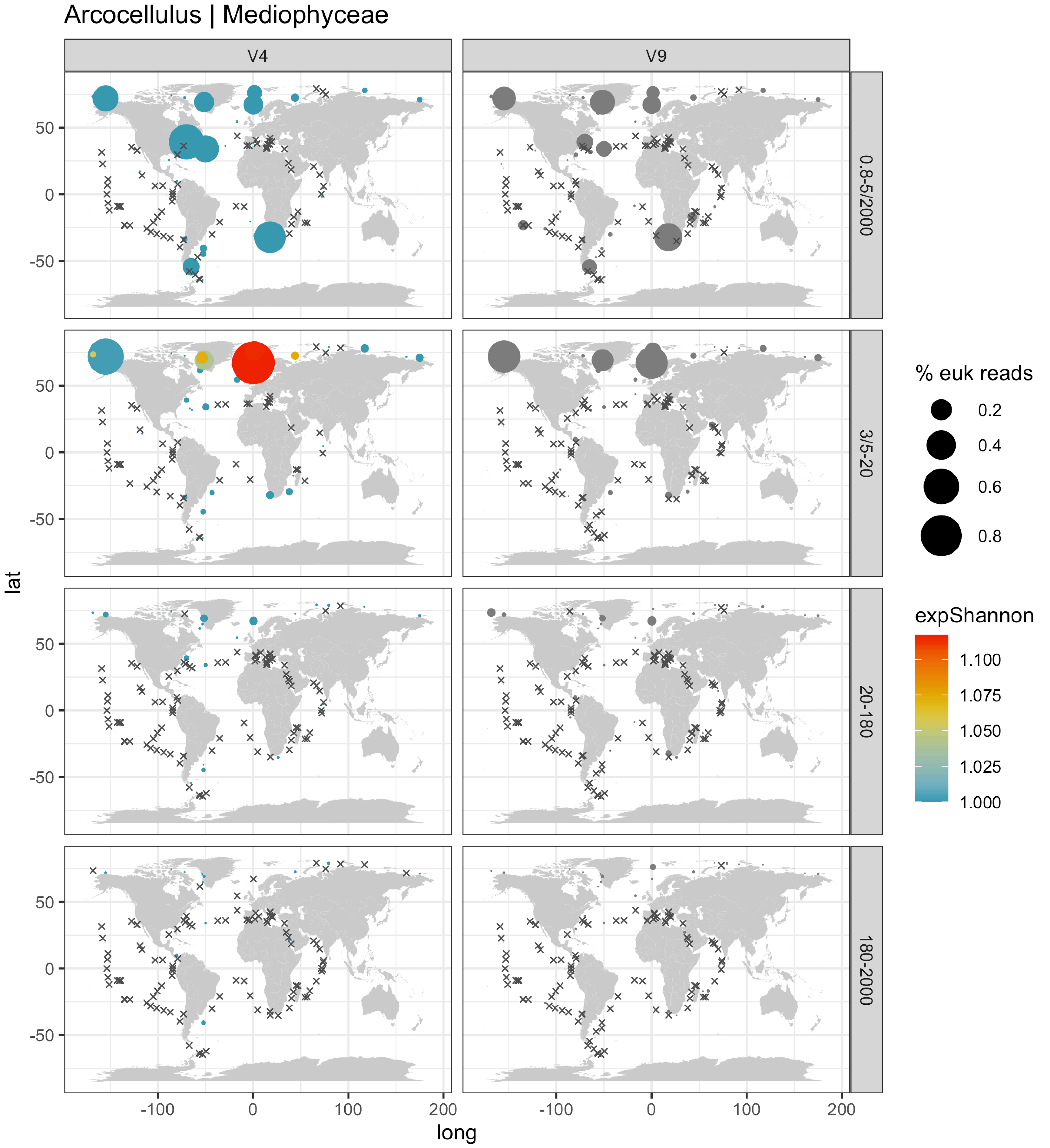

Arcocellulus | Mediophyceae
V4
V9
0.8-5/2000
3/5-20
20-180
180-2000
lat
long
% euk reads
0.2
0.4
0.6
0.8
expShannon
1.100
1.075
1.050
1.025
1.000

Ardissonea | Mediophyceae

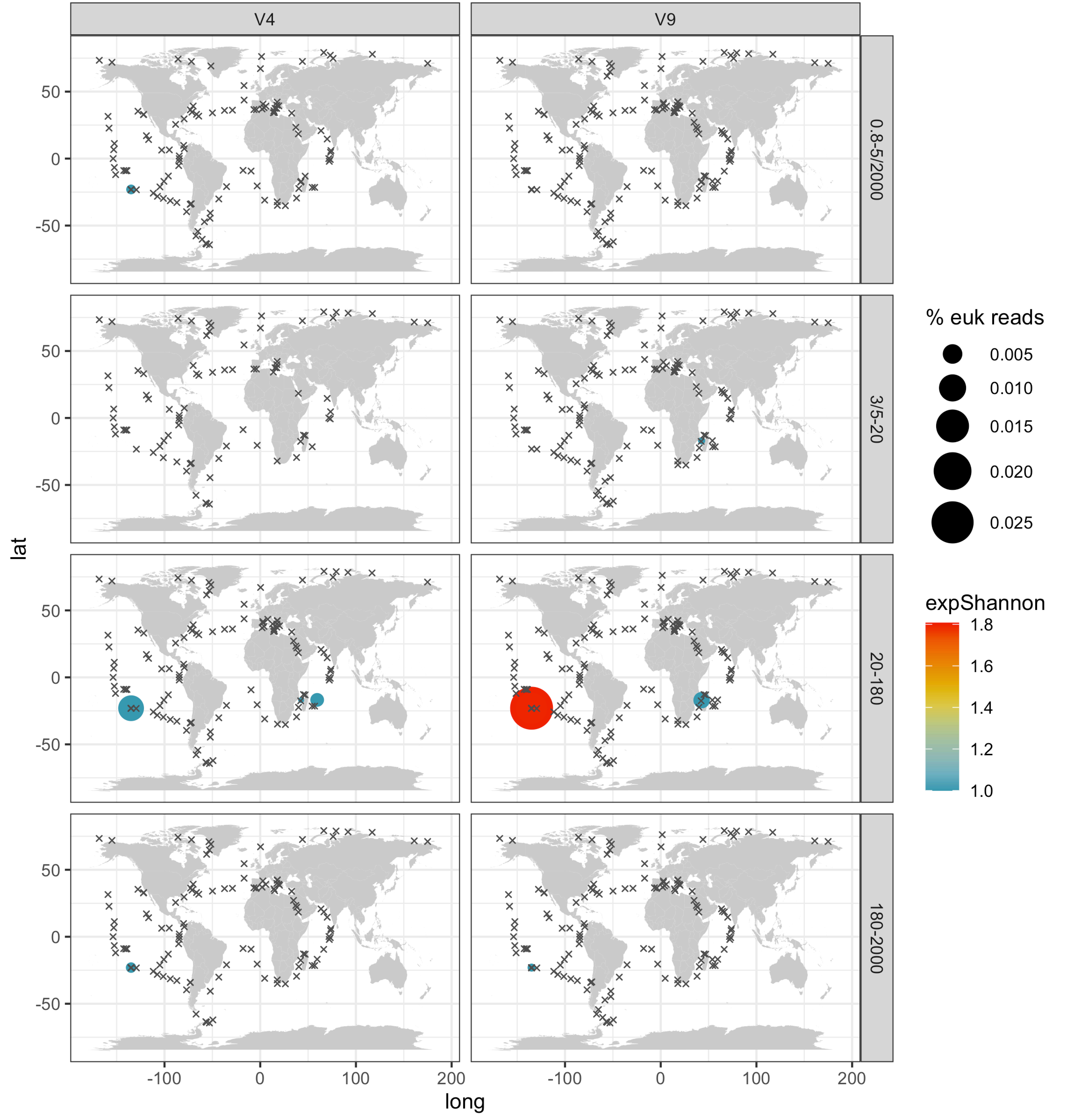

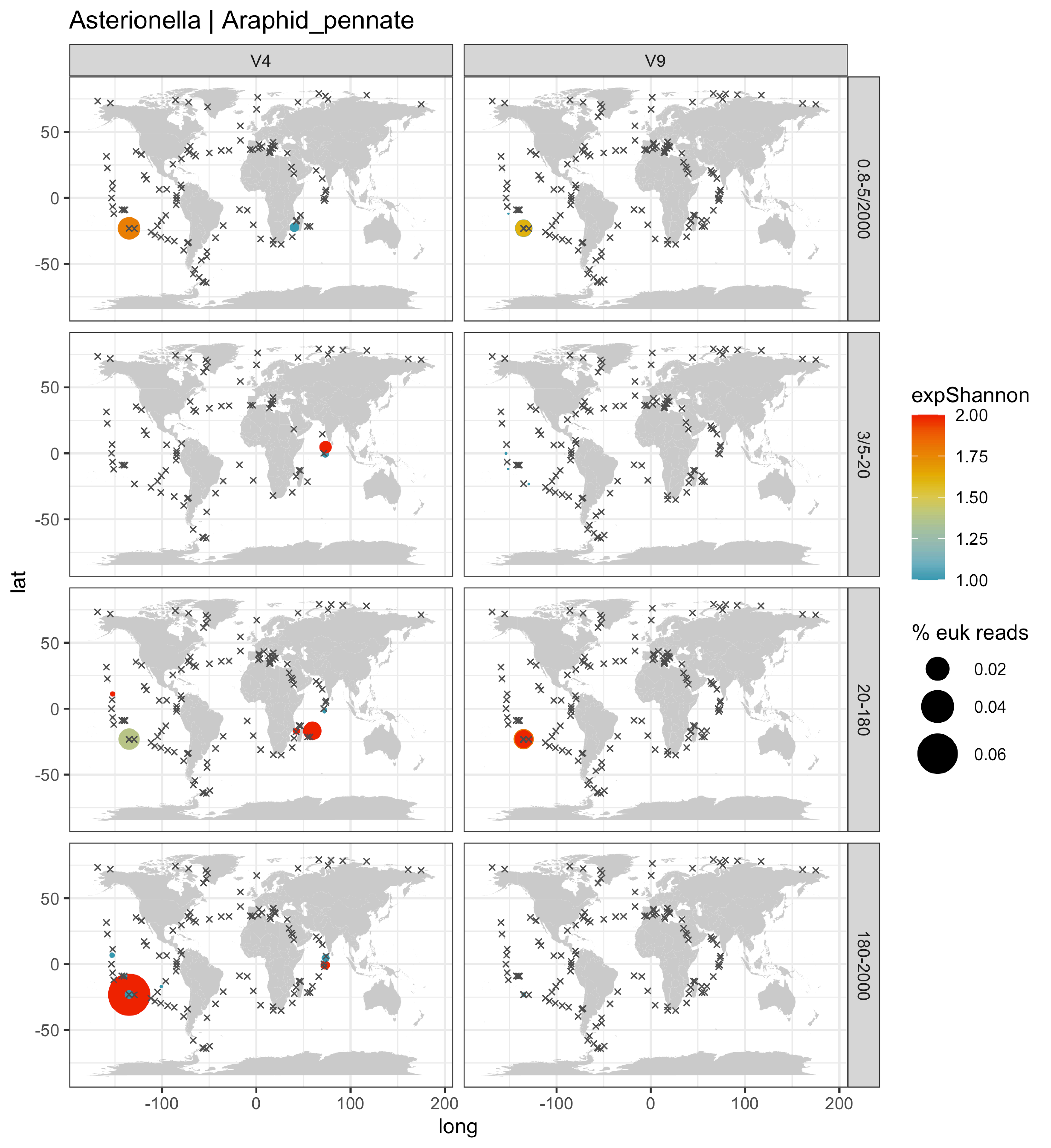

Asterionella | Araphid_pennate
V4
V9
0.8-5/2000
3/5-20
20-180
180-2000
lat
long
50
0
-50
-100
0
100
200
expShannon
2.00
1.75
1.50
1.25
1.00
% euk reads
0.02
0.04
0.06

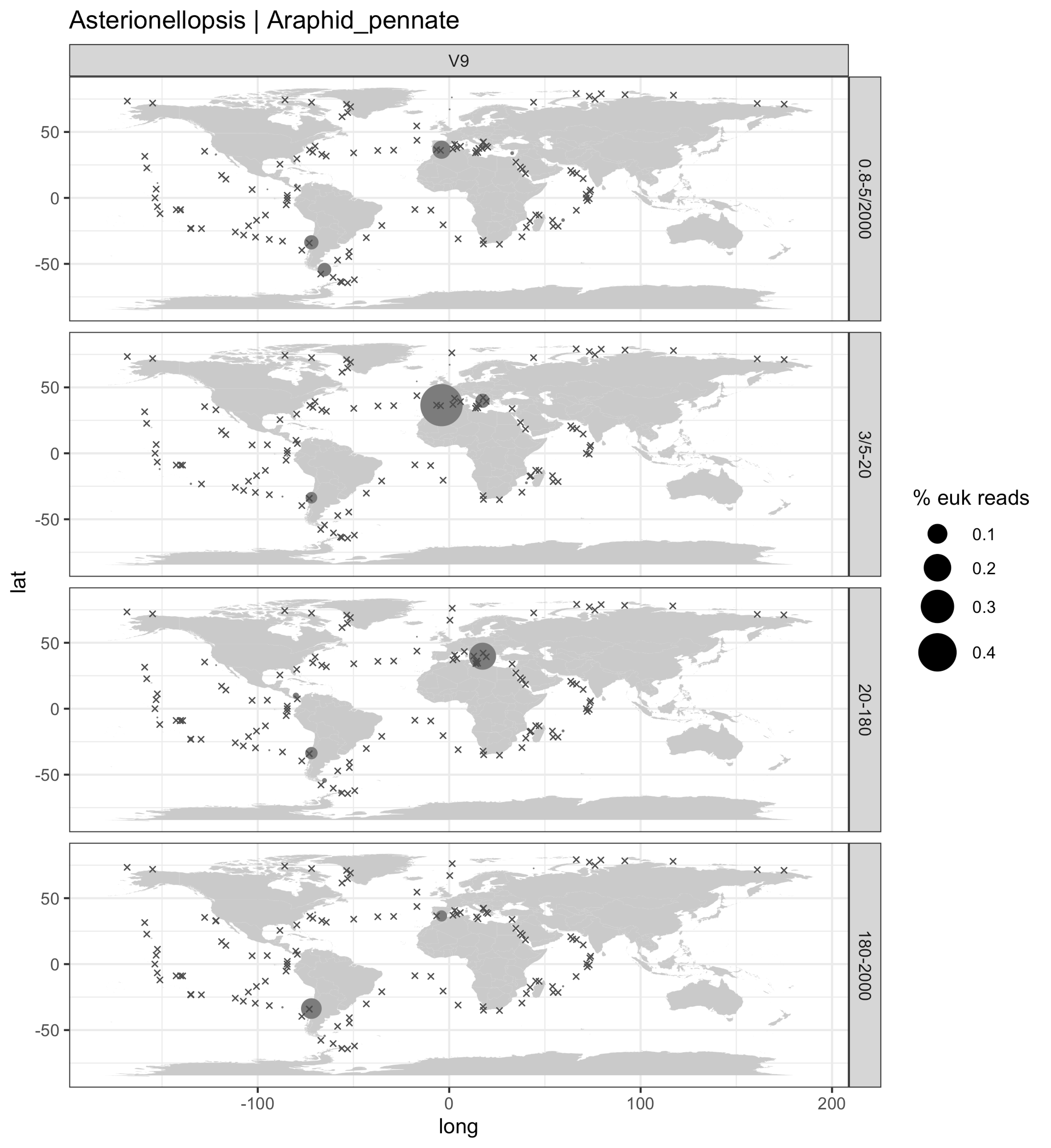

Asterionellopsis | Araphid_pennate
V9
0.8-5/2000
3/5-20
20-180
180-2000
lat
long
50
0
-50
-100
0
100
200
% euk reads
0.1
0.2
0.3
0.4

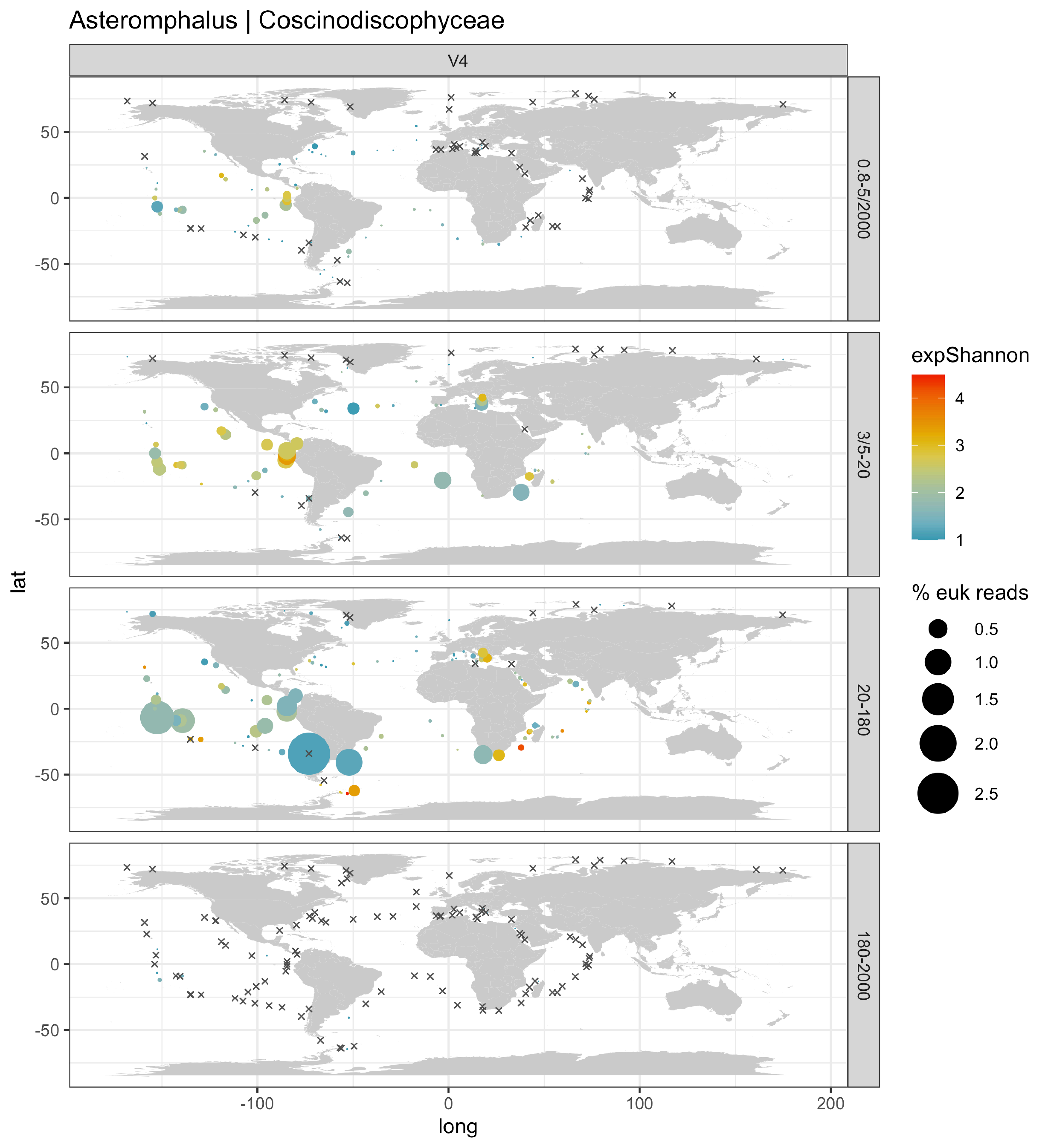
Asteromphalus | Coscinodiscophyceae
V4
0.8-5/2000
3/5-20
20-180
180-2000
lat
long
-100
0
100
200
50
0
-50
expShannon
4
3
2
1
% euk reads
0.5
1.0
1.5
2.0
2.5

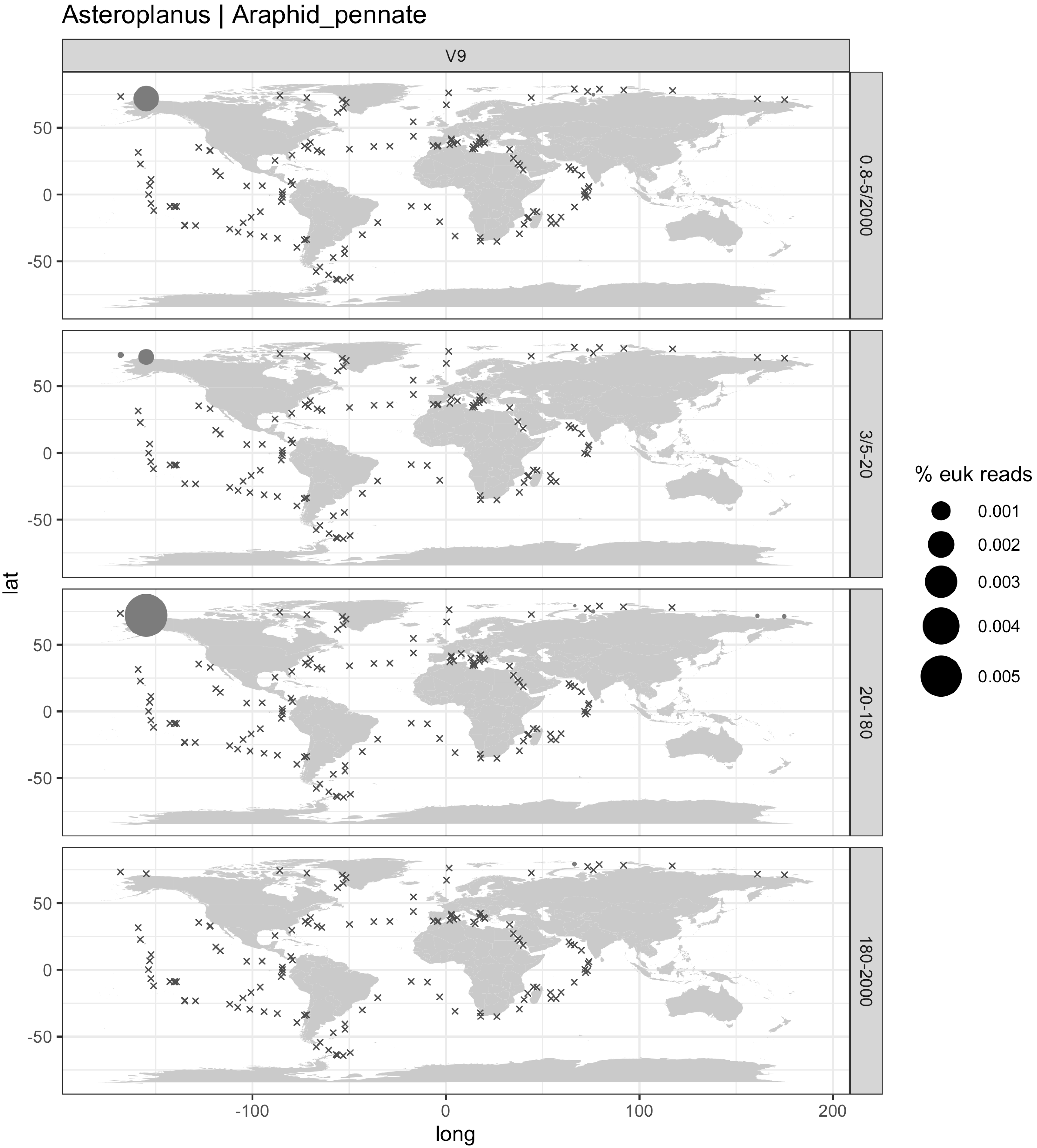

Asteroplanus | Araphid_pennate
V9
0.8-5/2000
3/5-20
20-180
180-2000
lat
long
50
0
-50
-100
100
200
% euk reads
0.001
0.002
0.003
0.004
0.005

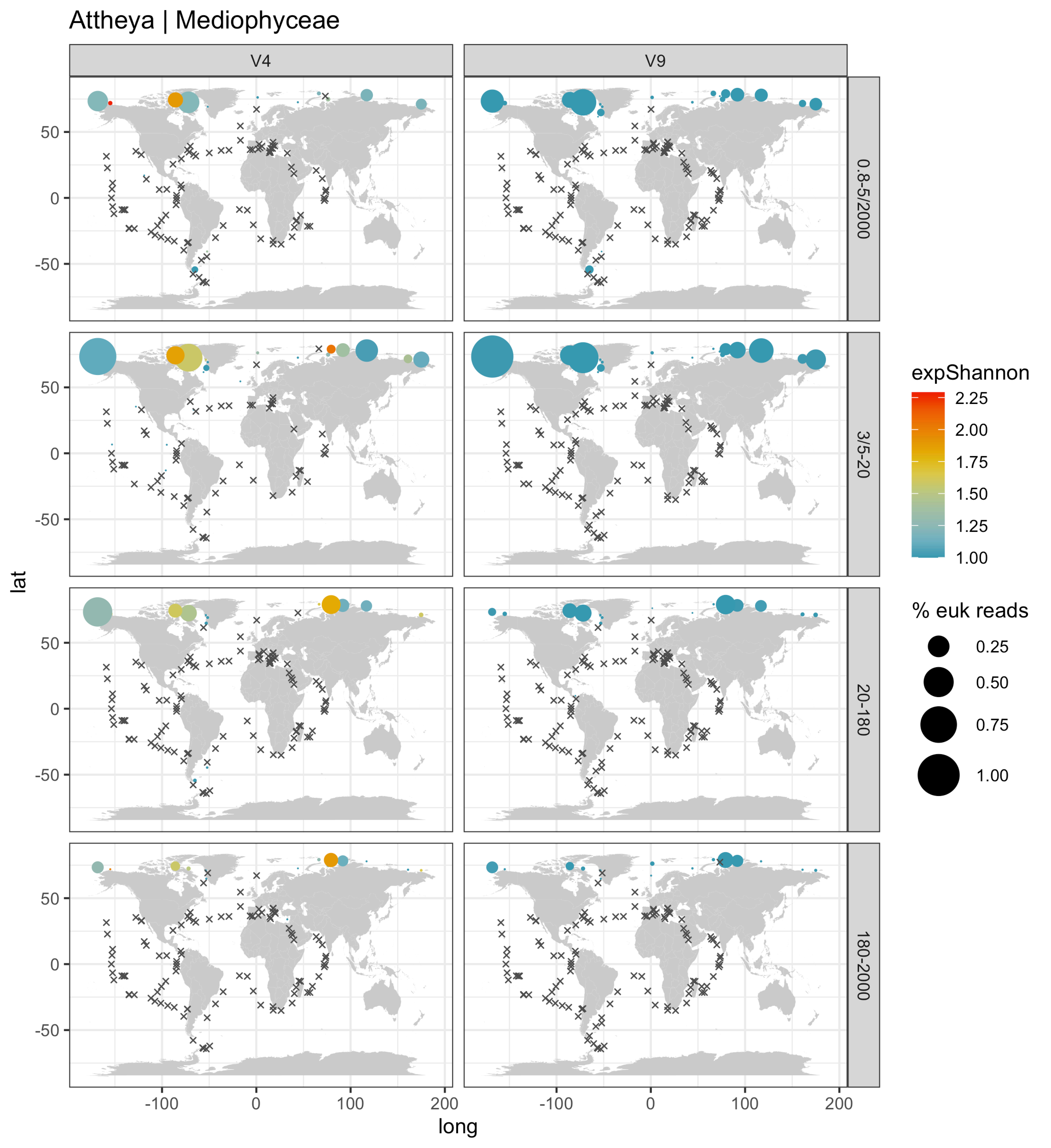

Attheya | Mediophyceae
V4
V9
0.8-5/2000
3/5-20
20-180
180-2000
lat
long
50
0
-50
-100
100
200
expShannon
2.25
2.00
1.75
1.50
1.25
1.00
% euk reads
0.25
0.50
0.75
1.00

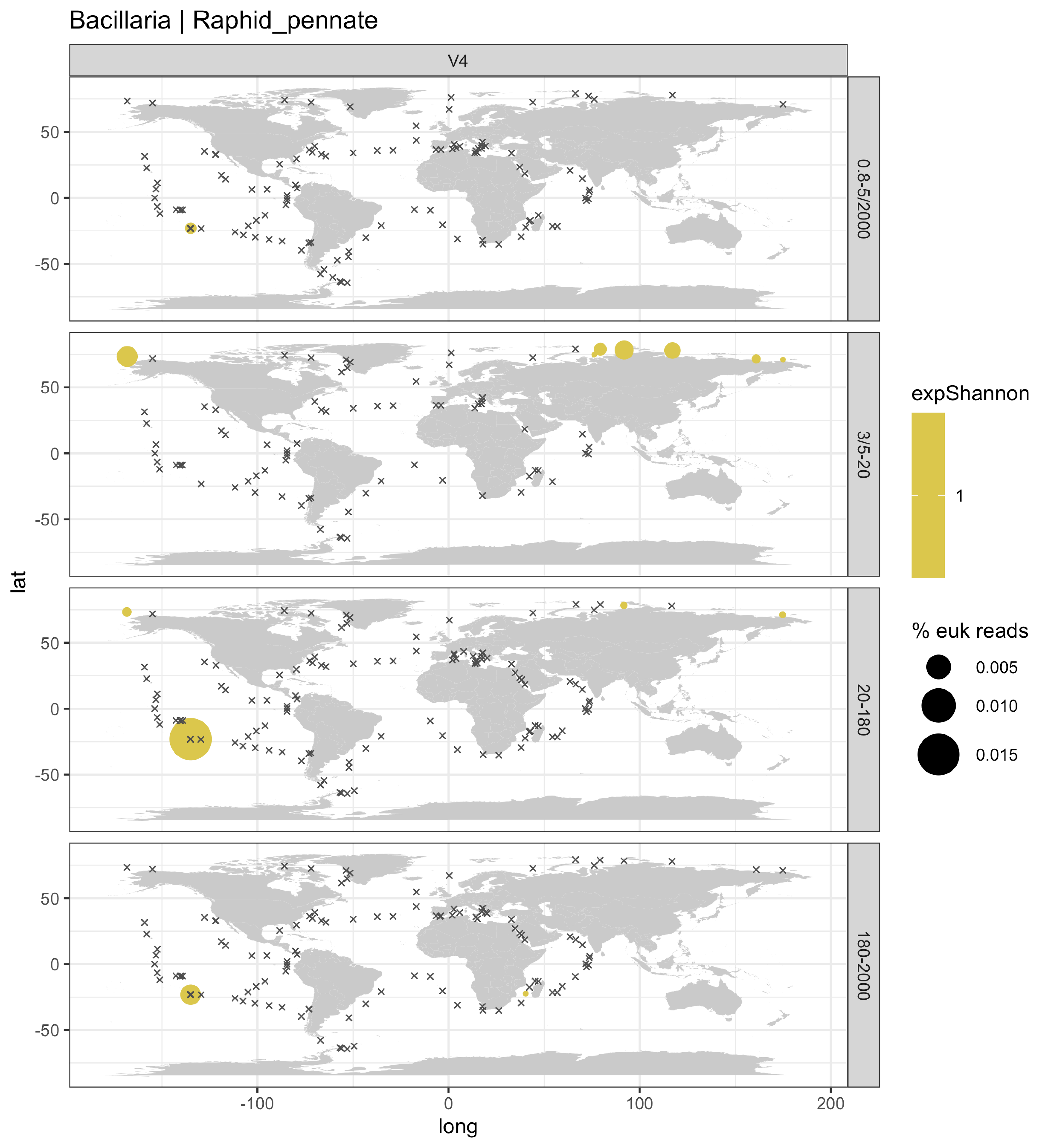

Bacillaria | Raphid_pennate
V4
0.8-5/2000
3/5-20
20-180
180-2000
lat
long
50
0
-50
-100
0
100
200
expShannon
1
% euk reads
0.005
0.010
0.015

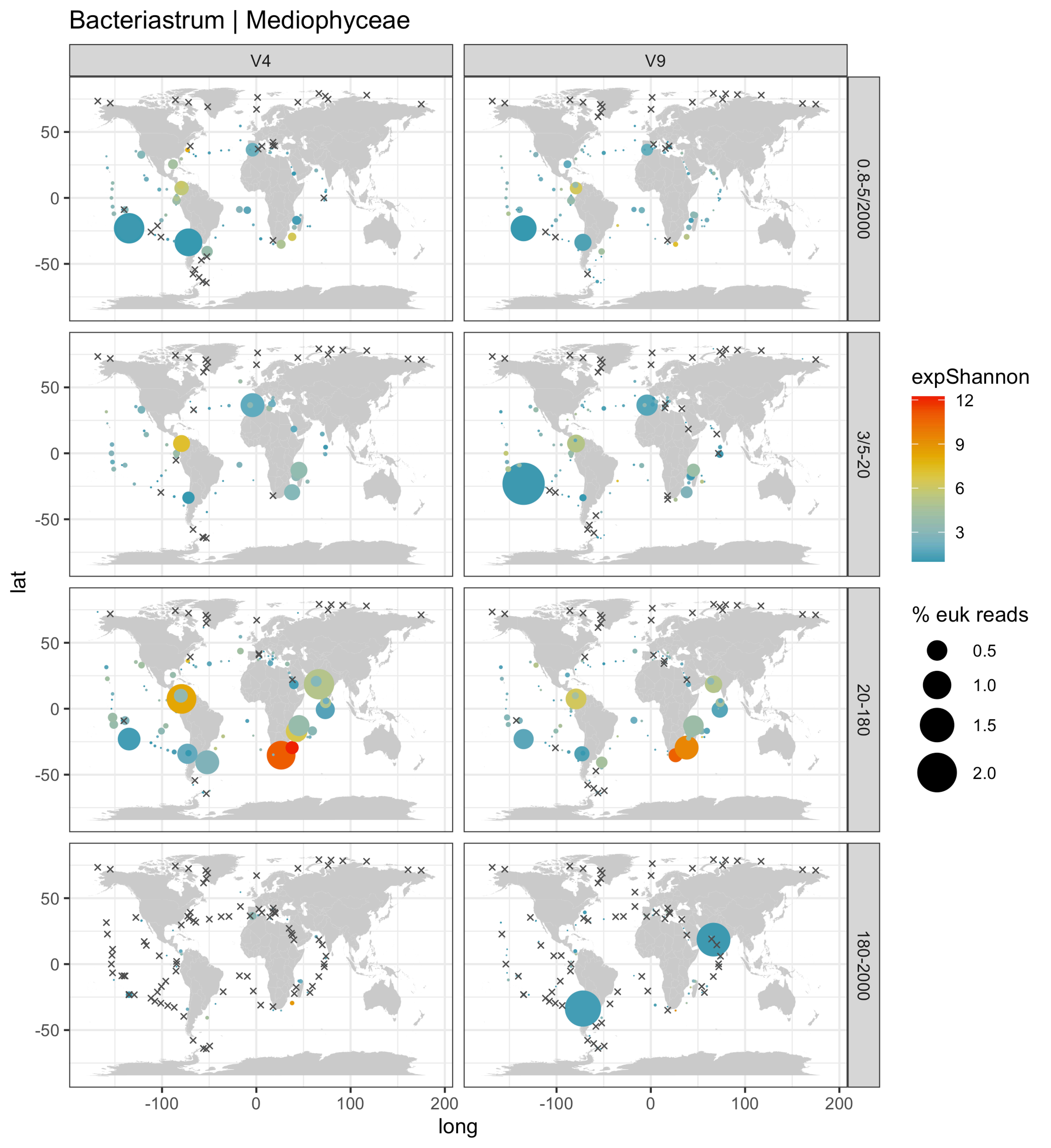

Bacteriastrum | Mediophyceae
V4
V9
0.8-5/2000
3/5-20
20-180
180-2000
lat
long
50
0
-50
-100
100
200
expShannon
12
9
6
3
% euk reads
0.5
1.0
1.5
2.0

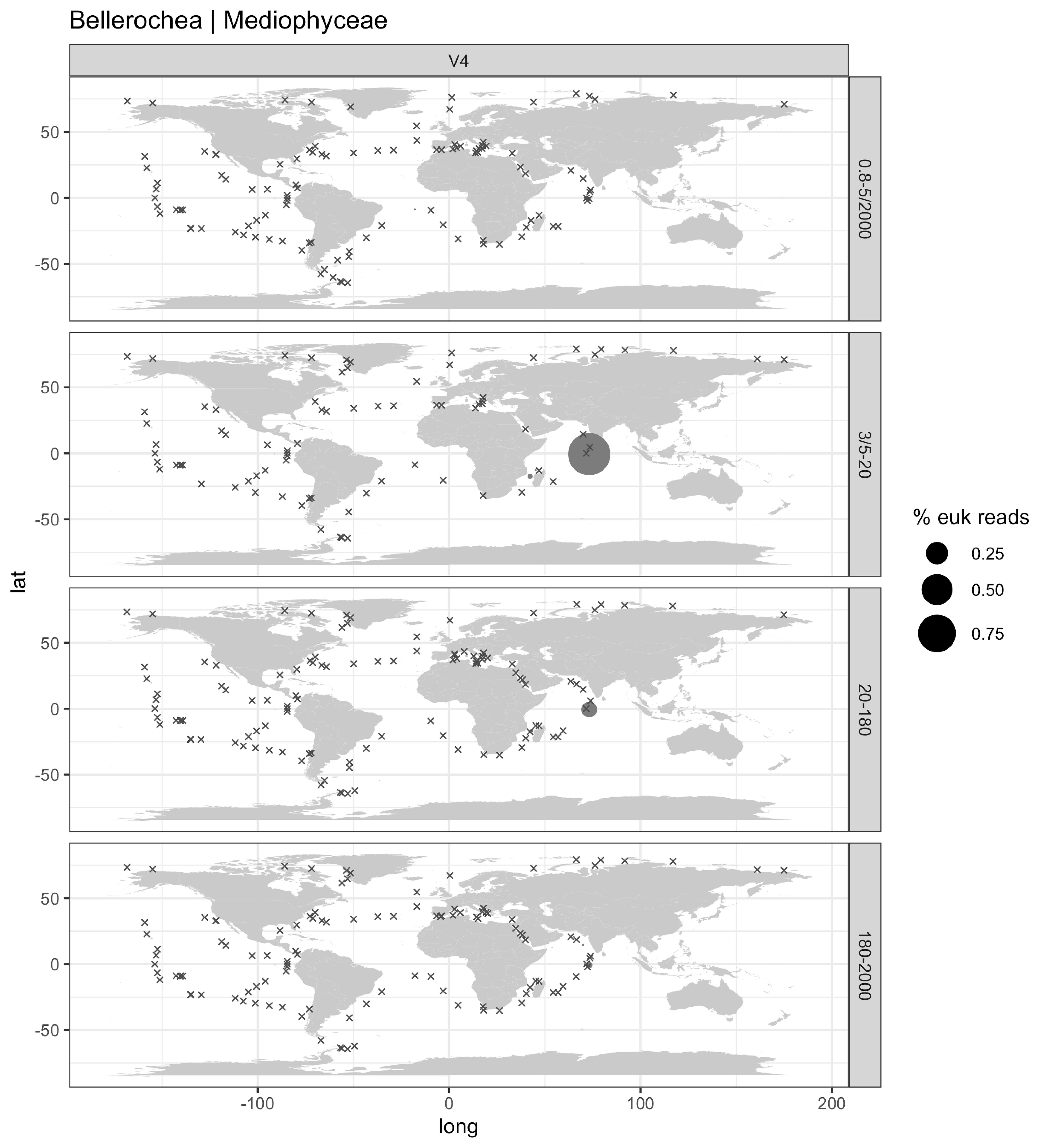
Bellerochea | Mediophyceae
V4
0.8-5/2000
3/5-20
20-180
180-2000
lat
long
50
0
-50
-100
0
100
200
% euk reads
0.25
0.50
0.75

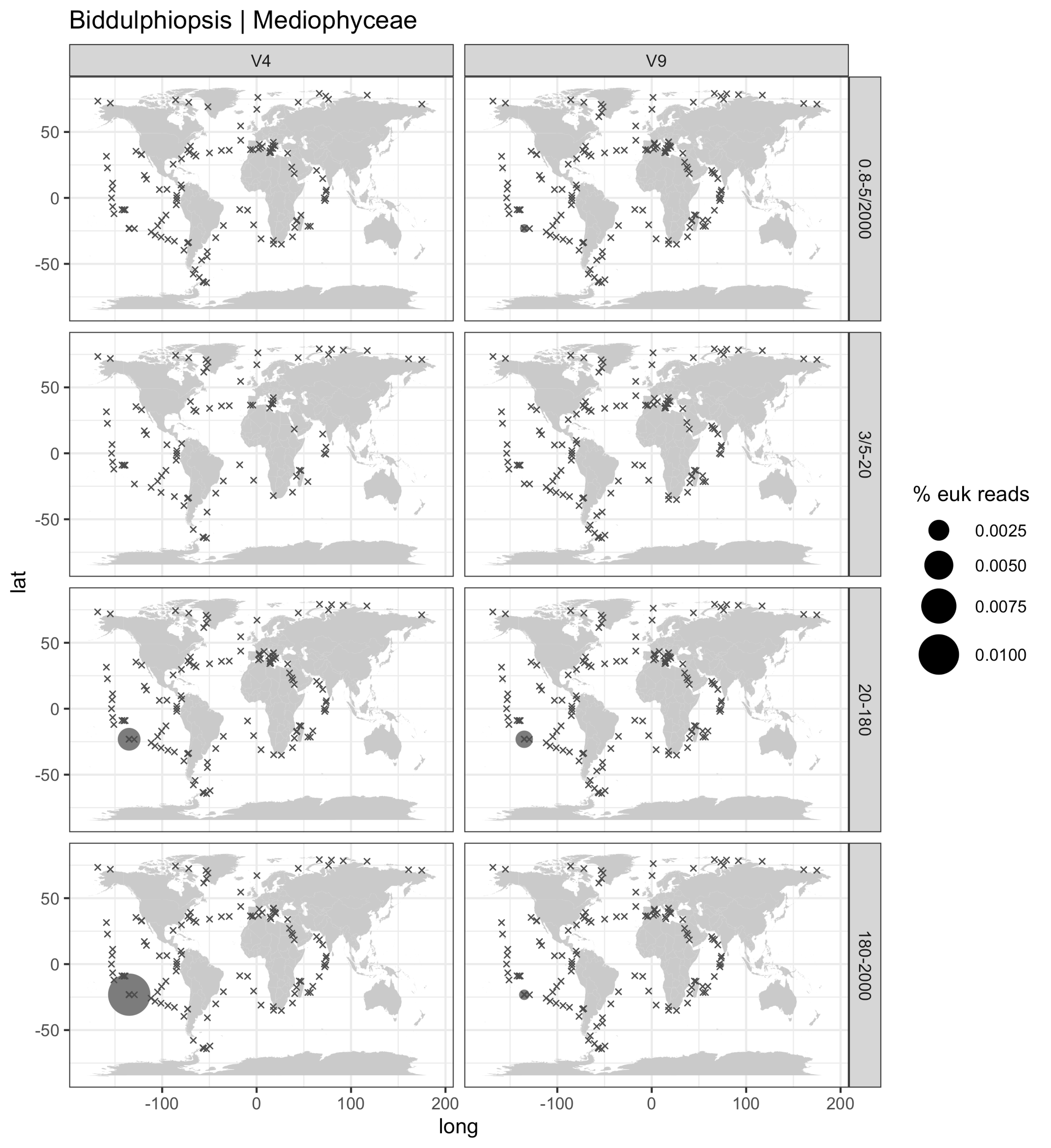

Biddulphiopsis | Mediophyceae
V4
V9
0.8-5/2000
3/5-20
20-180
180-2000
lat
long
-100
0
100
200
50
0
-50
% euk reads
0.0025
0.0050
0.0075
0.0100

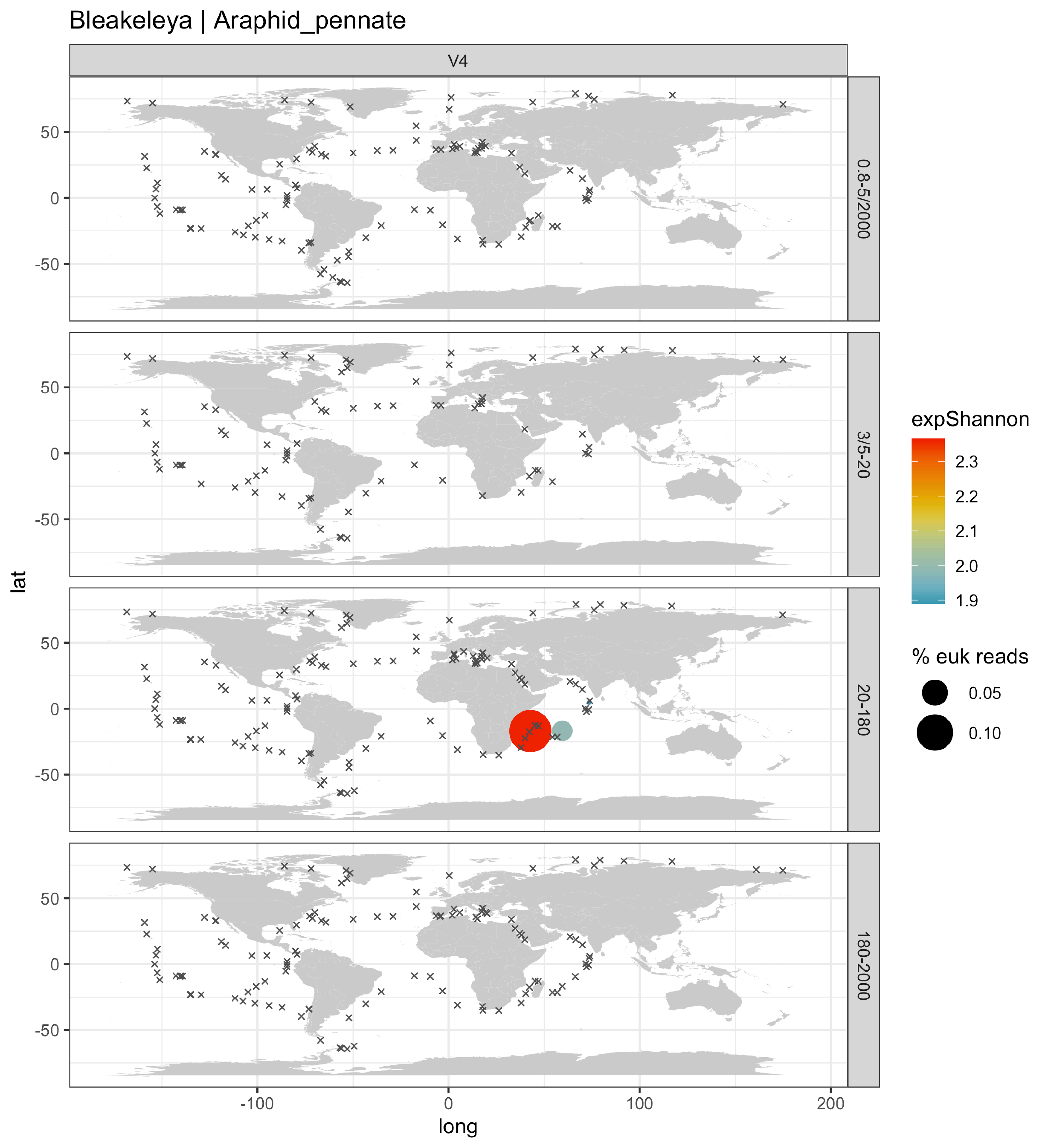
Bleakeleya | Araphid_pennate
V4
0.8-5/2000
3/5-20
20-180
180-2000
lat
long
50
0
-50
-100
100
200
expShannon
2.3
2.2
2.1
2.0
1.9
% euk reads
0.05
0.10

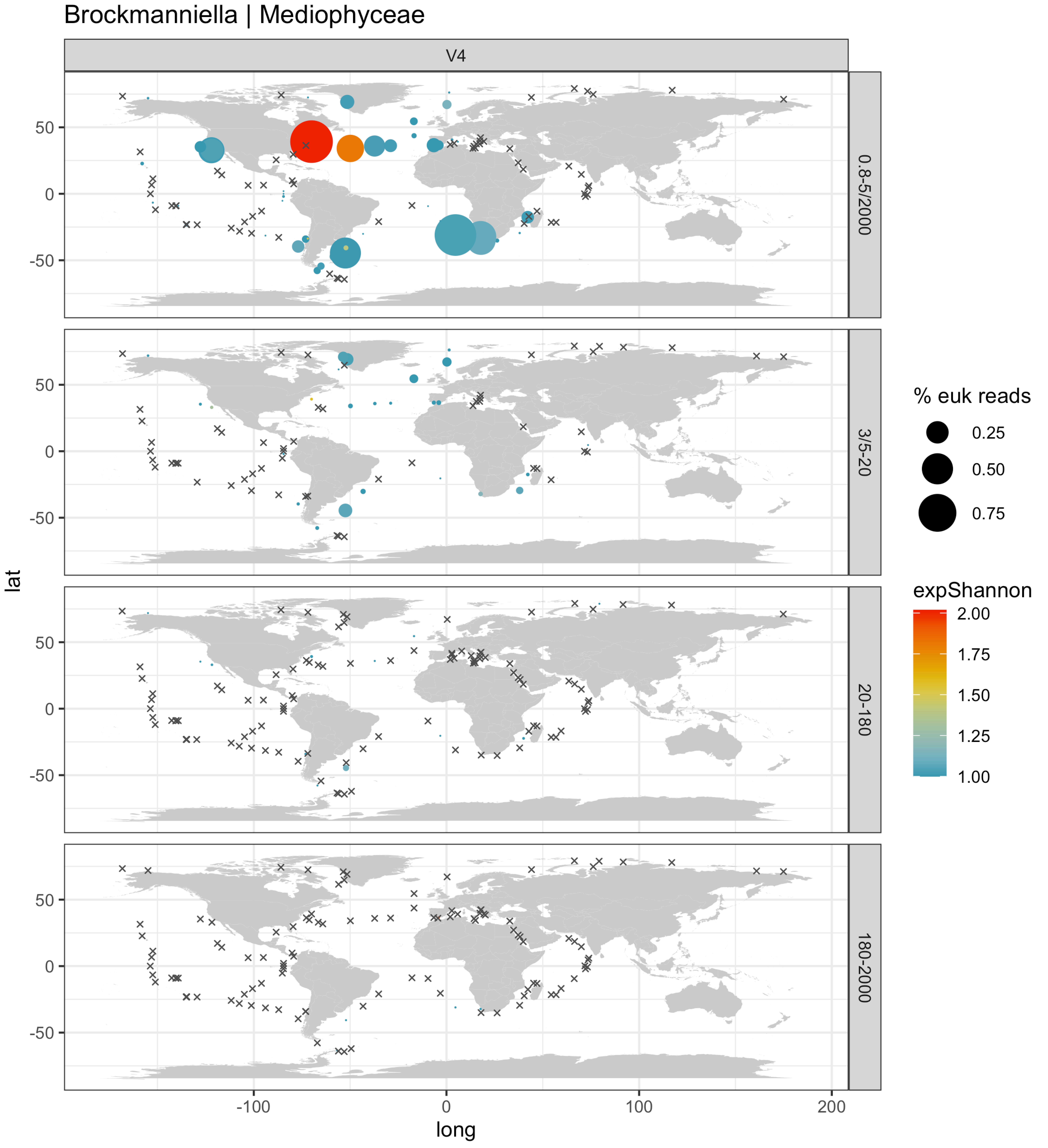

Brockmanniella | Mediophyceae
V4
0.8-5/2000
3/5-20
20-180
180-2000
lat
long
50
0
-50
-100
0
100
200
% euk reads
0.25
0.50
0.75
expShannon
2.00
1.75
1.50
1.25
1.00

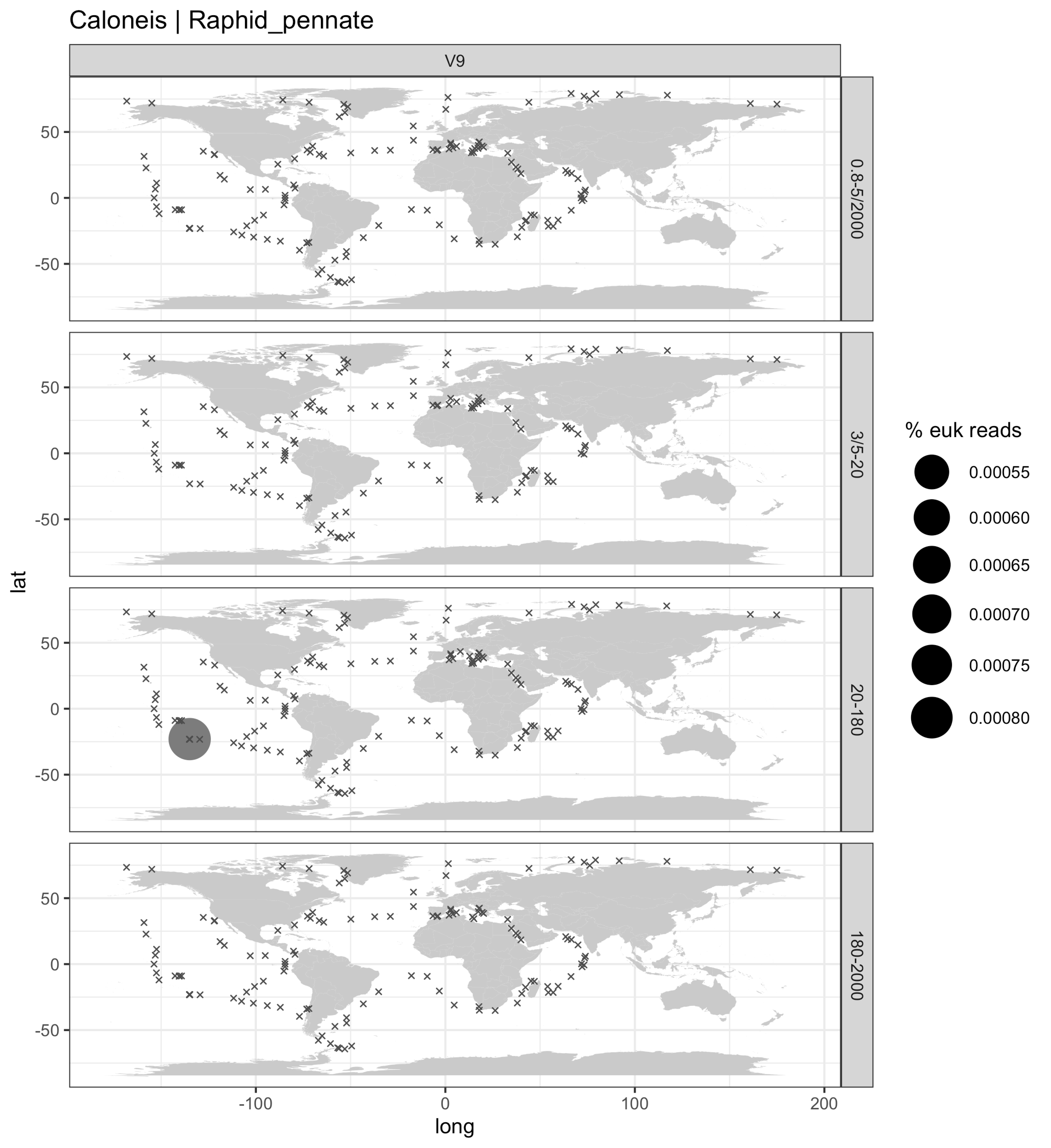
Caloneis | Raphid_pennate
V9
0.8-5/2000
3/5-20
20-180
180-2000
lat
long
50
0
-50
-100
100
200
% euk reads
0.00055
0.00060
0.00065
0.00070
0.00075
0.00080

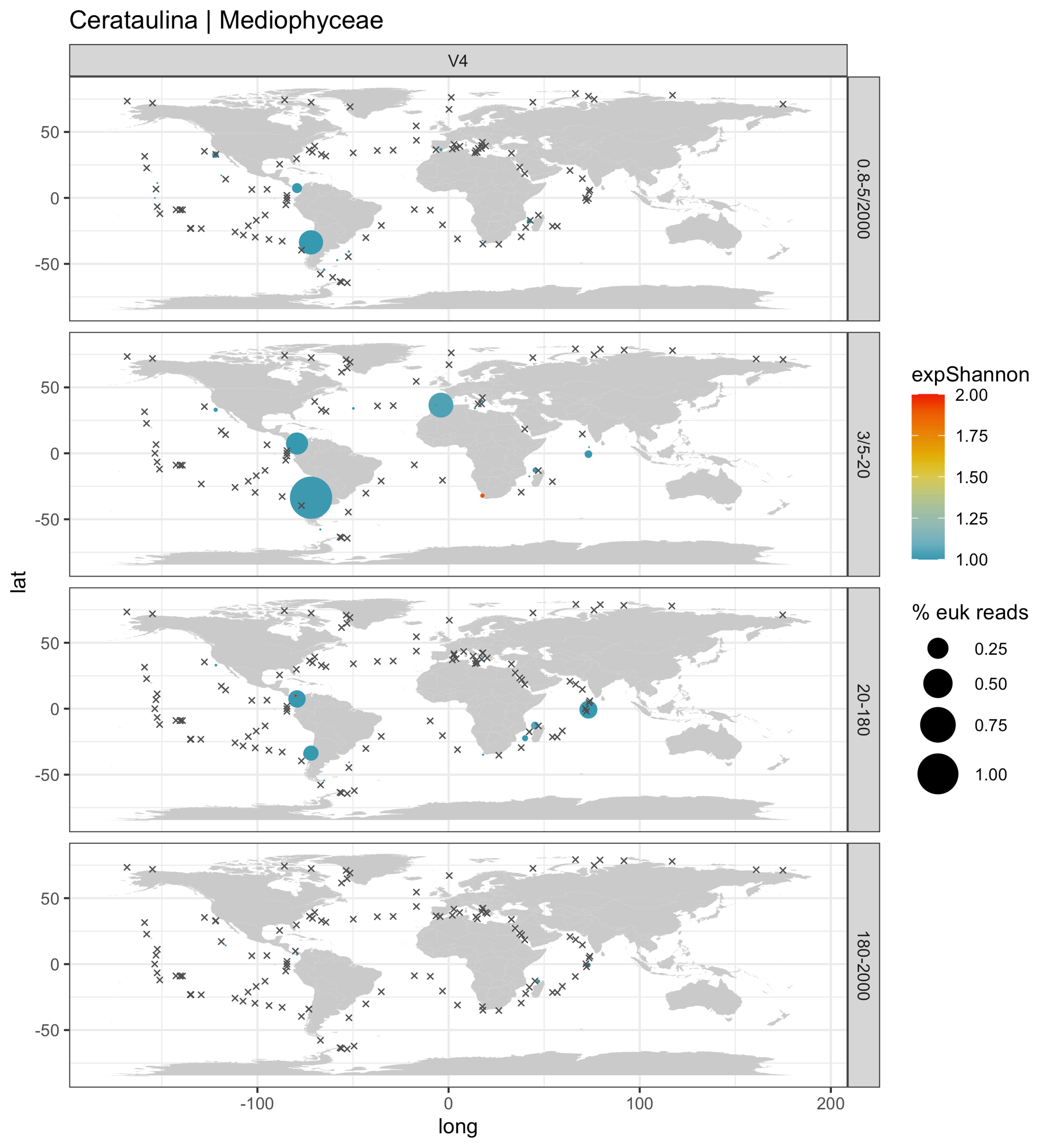
Cerataulina | Mediophyceae
V4
0.8-5/2000
3/5-20
20-180
180-2000
lat
long
expShannon
2.00
1.75
1.50
1.25
1.00
% euk reads
0.25
0.50
0.75
1.00
50
0
-50
-100
100
200

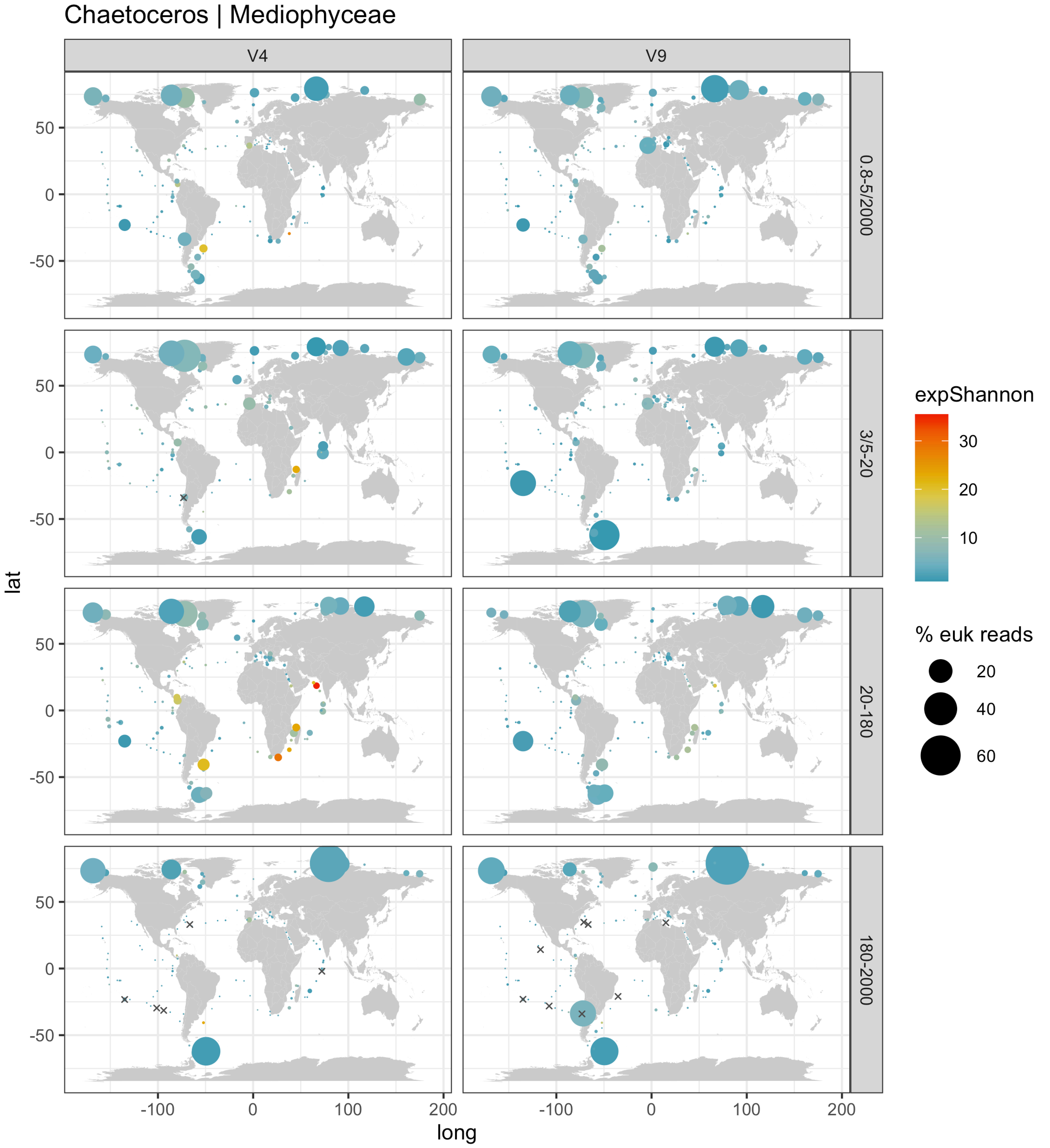

Chaetoceros | Mediophyceae
V4
V9
0.8-5/2000
3/5-20
20-180
180-2000
lat
long
50
0
-50
-100
0
100
200
expShannon
30
20
10
% euk reads
20
40
60

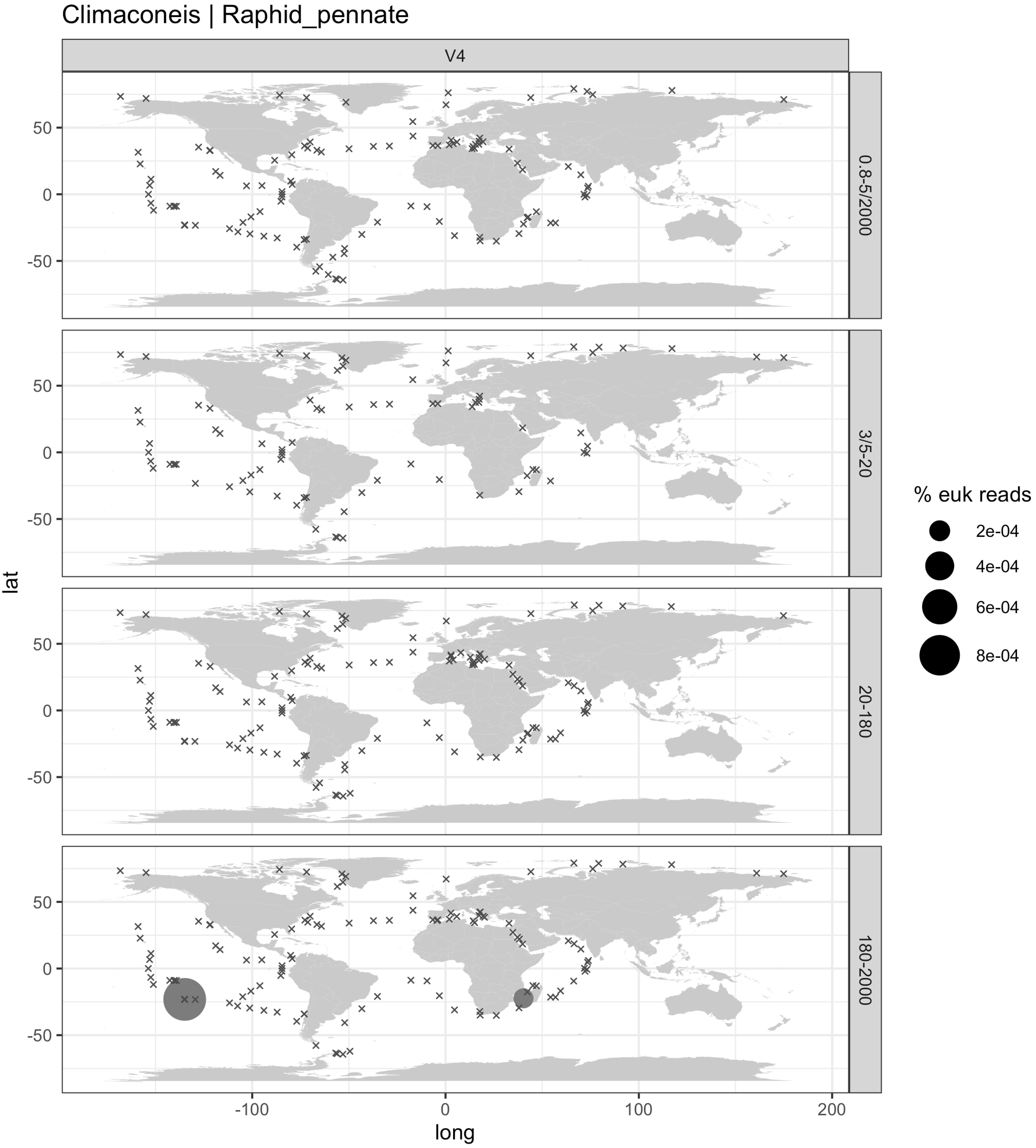

Climaconeis | Raphid_pennate
V4
0.8-5/2000
3/5-20
20-180
180-2000
lat
long
50
0
-50
-100
100
200
% euk reads
2e-04
4e-04
6e-04
8e-04

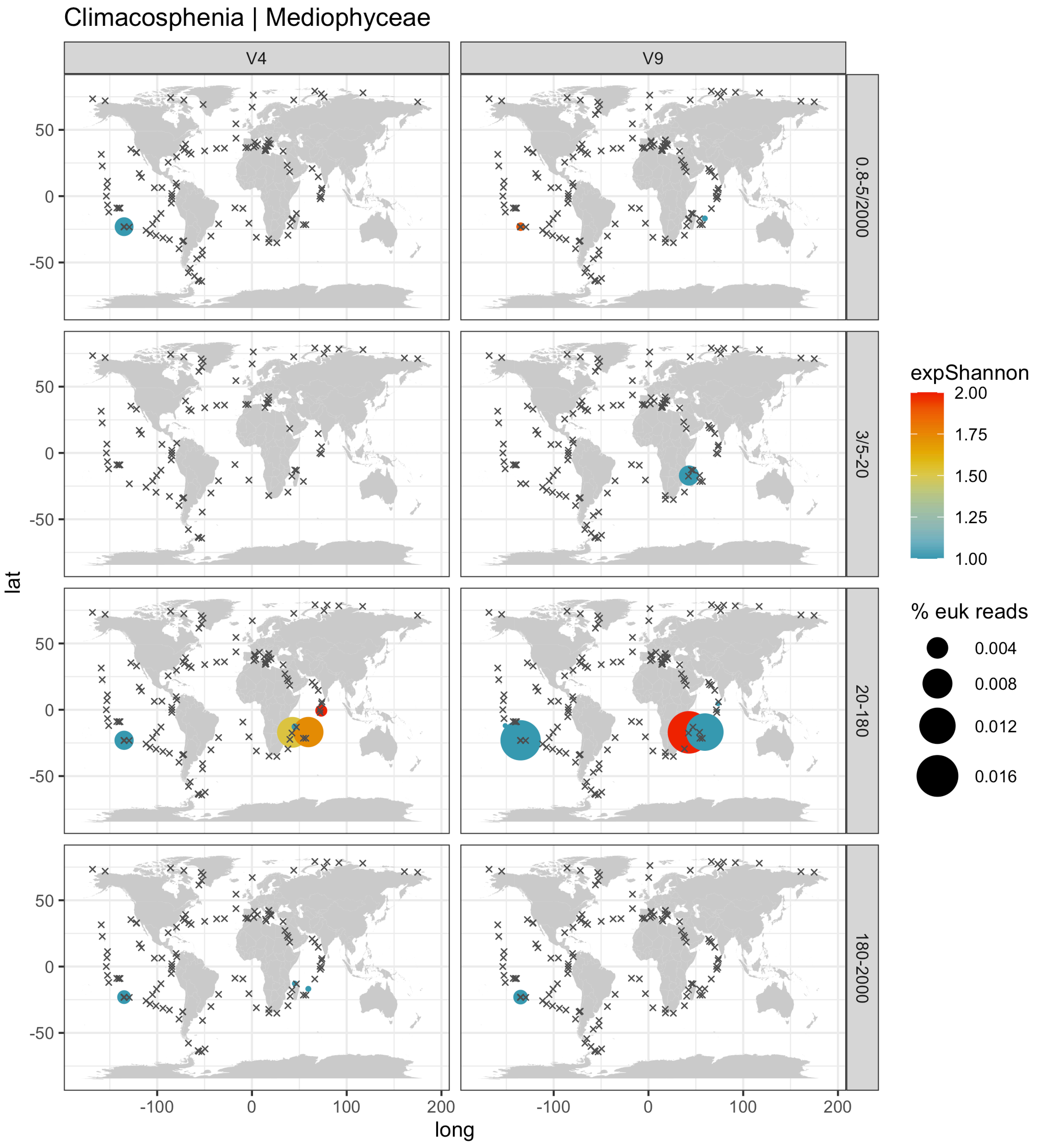

Climacosphenia | Mediophyceae
V4
V9
0.8-5/2000
3/5-20
20-180
180-2000
lat
long
50
0
-50
-100
0
100
200
expShannon
2.00
1.75
1.50
1.25
1.00
% euk reads
0.004
0.008
0.012
0.016

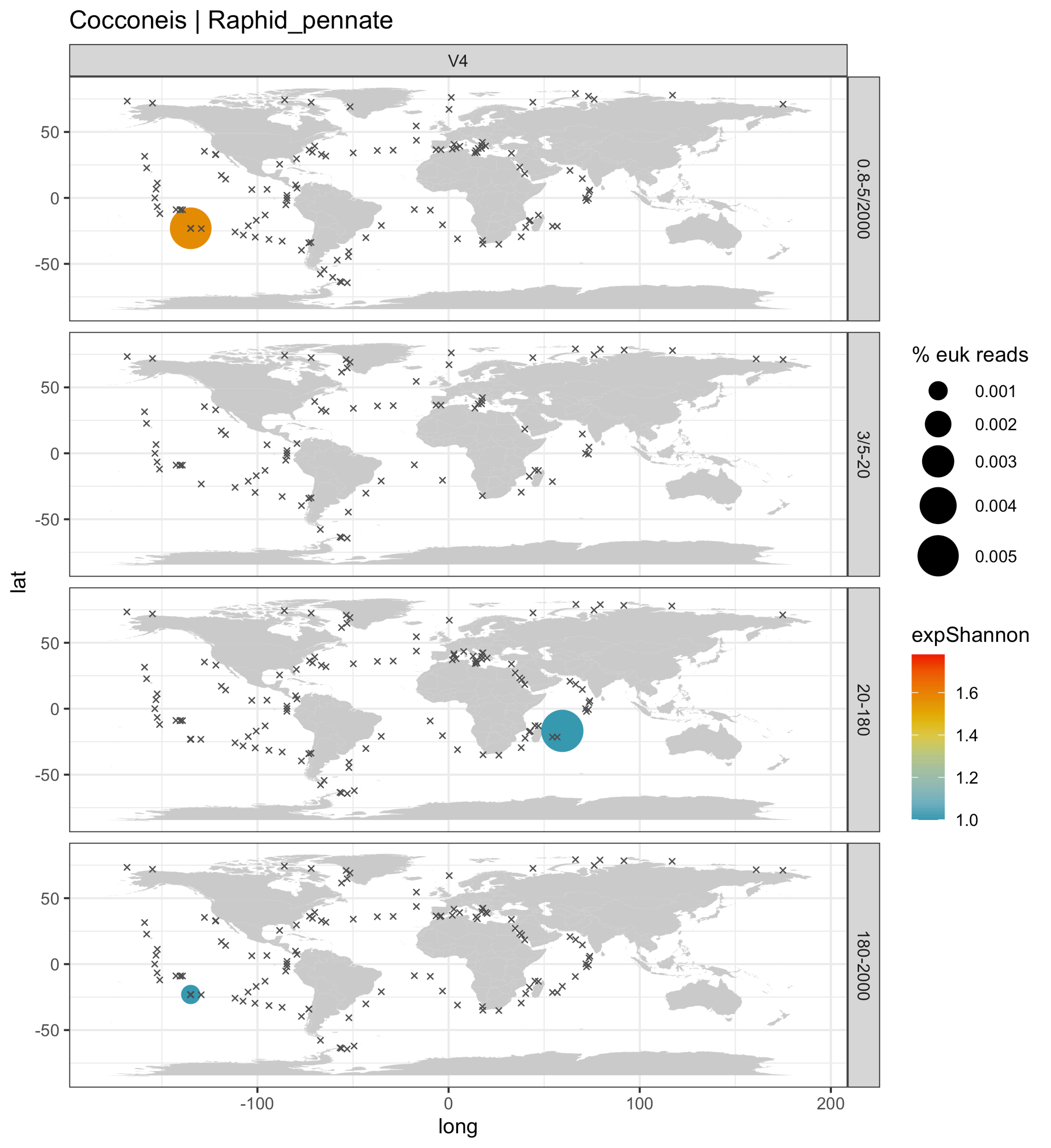

Cocconeis | Raphid_pennate
V4
0.8-5/2000
3/5-20
20-180
180-2000
lat
long
% euk reads
0.001
0.002
0.003
0.004
0.005
expShannon
1.6
1.4
1.2
1.0

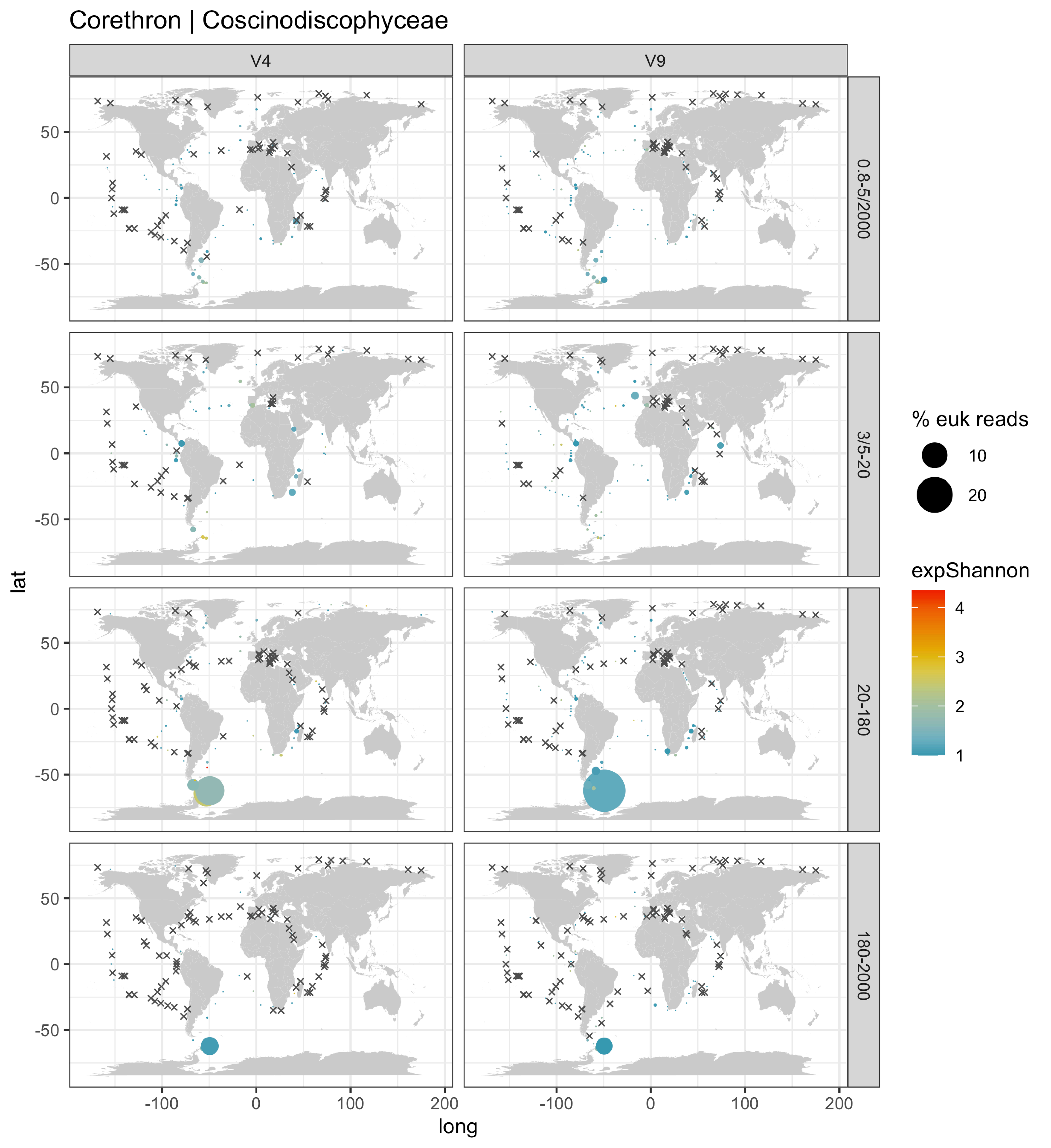
Corethron | Coscinodiscophyceae
V4
V9
0.8-5/2000
3/5-20
20-180
180-2000
lat
long
50
0
-50
-100
0
100
200
% euk reads
10
20
expShannon
4
3
2
1

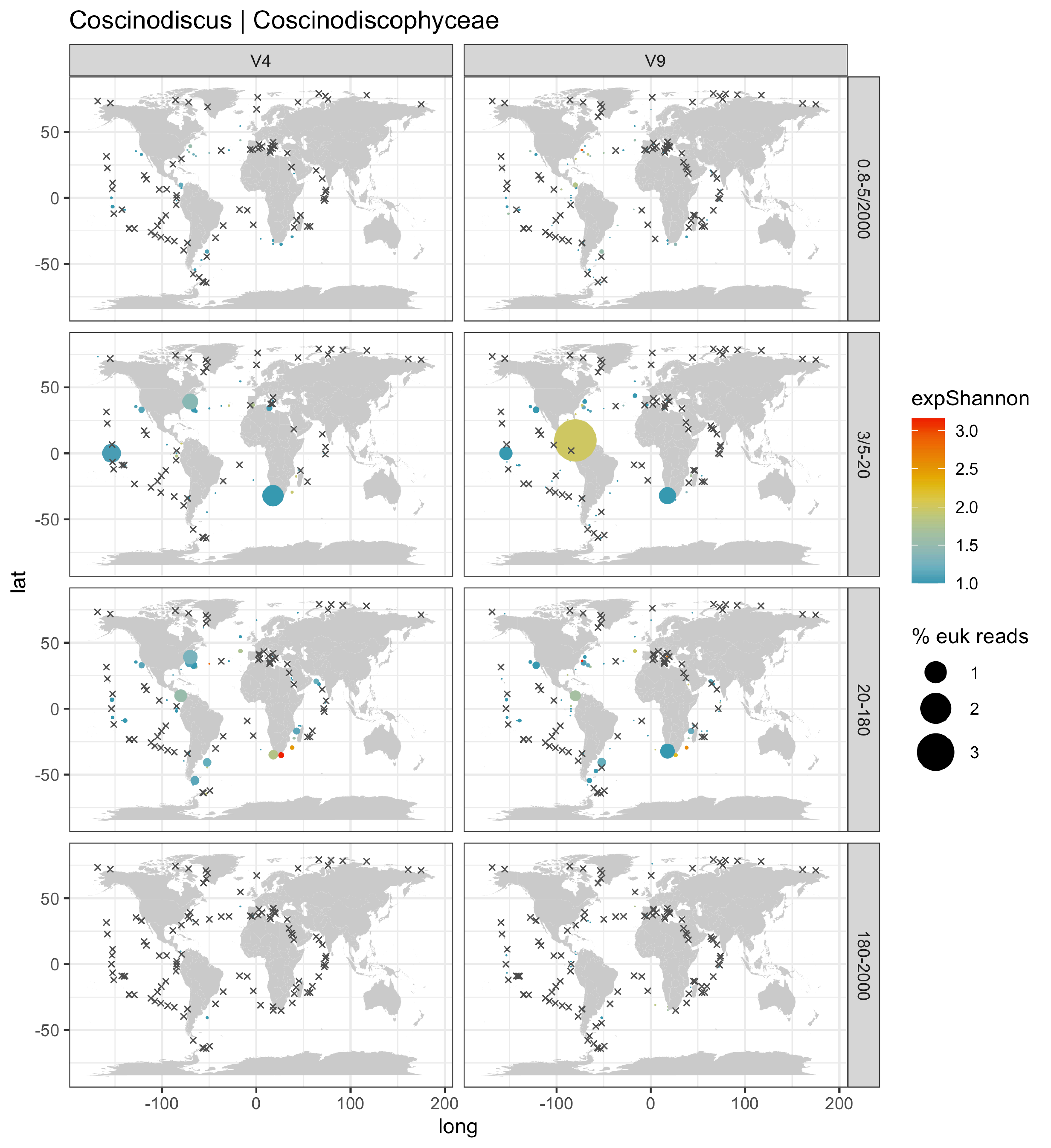

Coscinodiscus | Coscinodiscophyceae
V4
V9
0.8-5/2000
3/5-20
20-180
180-2000
lat
long
50
0
-50
-100
0
100
200
expShannon
3.0
2.5
2.0
1.5
1.0
% euk reads
1
2
3

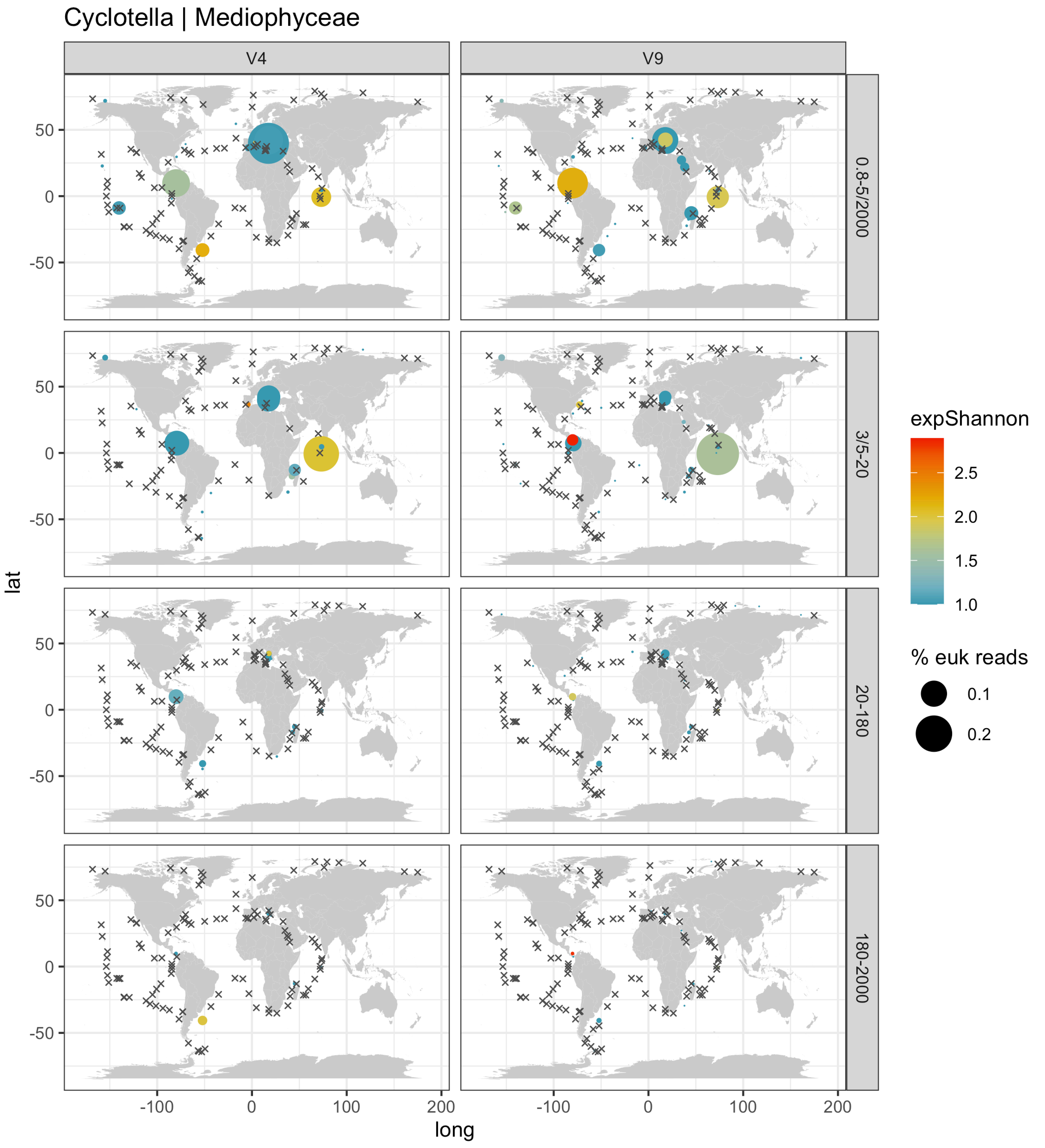
Cyclotella | Mediophyceae
V4
V9
0.8-5/2000
3/5-20
20-180
180-2000
lat
long
50
0
-50
-100
0
100
200
expShannon
2.5
2.0
1.5
1.0
% euk reads
0.1
0.2

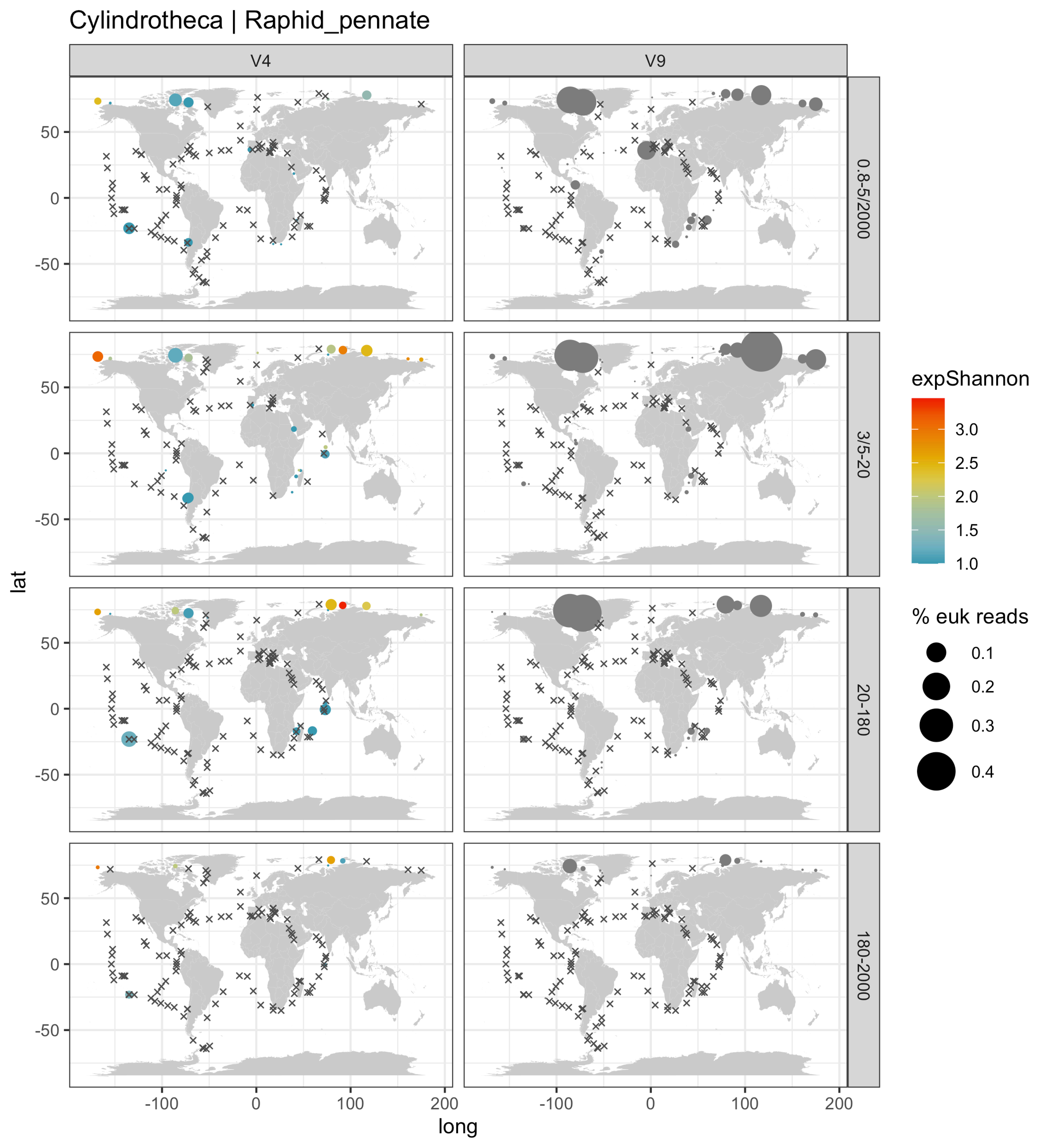

Cylindrotheca | Raphid_pennate
V4
V9
0.8-5/2000
3/5-20
20-180
180-2000
lat
long
-100
0
100
200
50
0
-50
expShannon
3.0
2.5
2.0
1.5
1.0
% euk reads
0.1
0.2
0.3
0.4

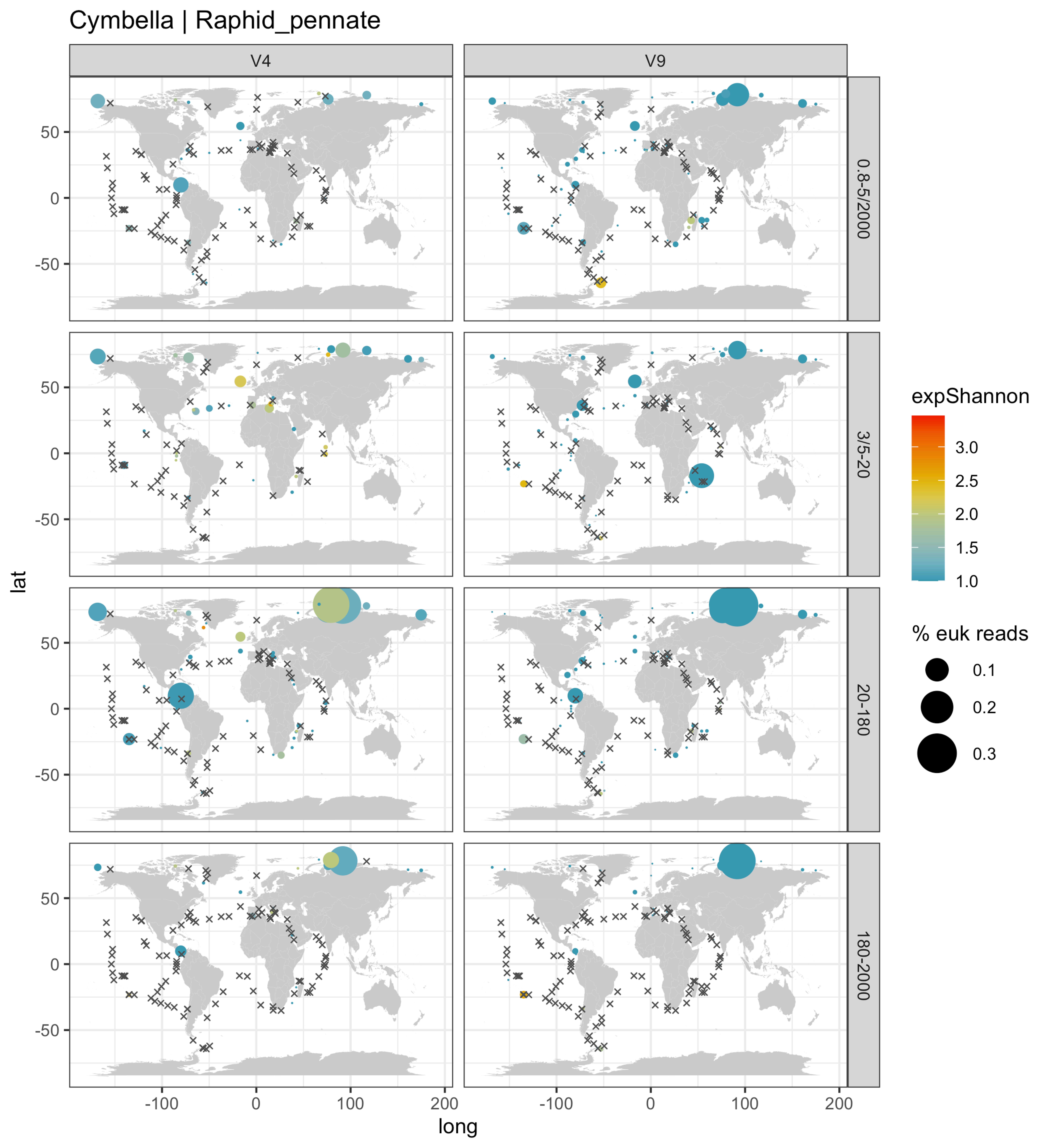
Cymbella | Raphid_pennate
V4
V9
0.8-5/2000
3/5-20
20-180
180-2000
lat
long
50
0
-50
-100
100
200
expShannon
3.0
2.5
2.0
1.5
1.0
% euk reads
0.1
0.2
0.3

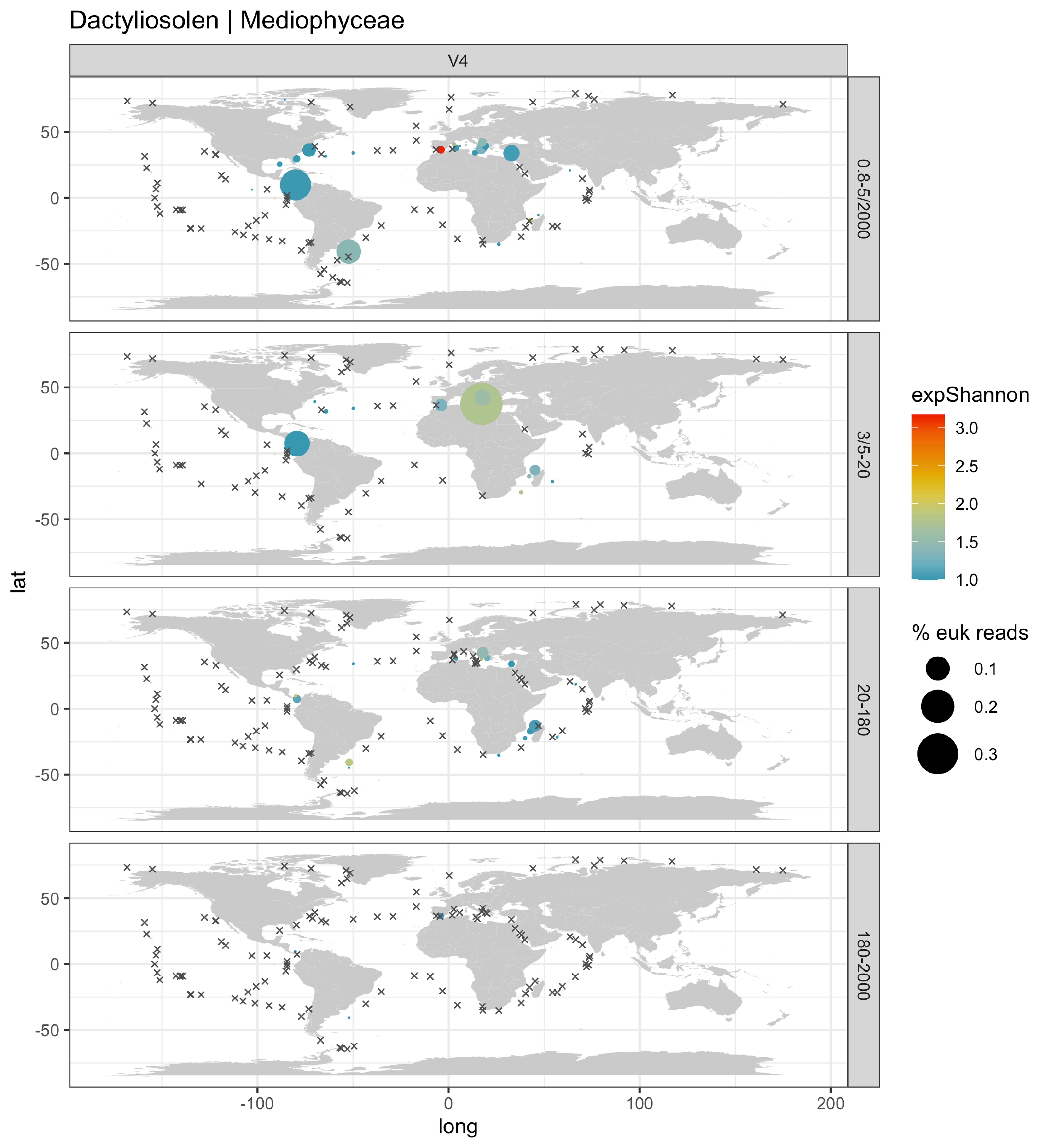

Dactyliosolen | Mediophyceae
V4
0.8-5/2000
3/5-20
20-180
180-2000
lat
long
-100
0
100
200
50
0
-50
expShannon
3.0
2.5
2.0
1.5
1.0
% euk reads
0.1
0.2
0.3

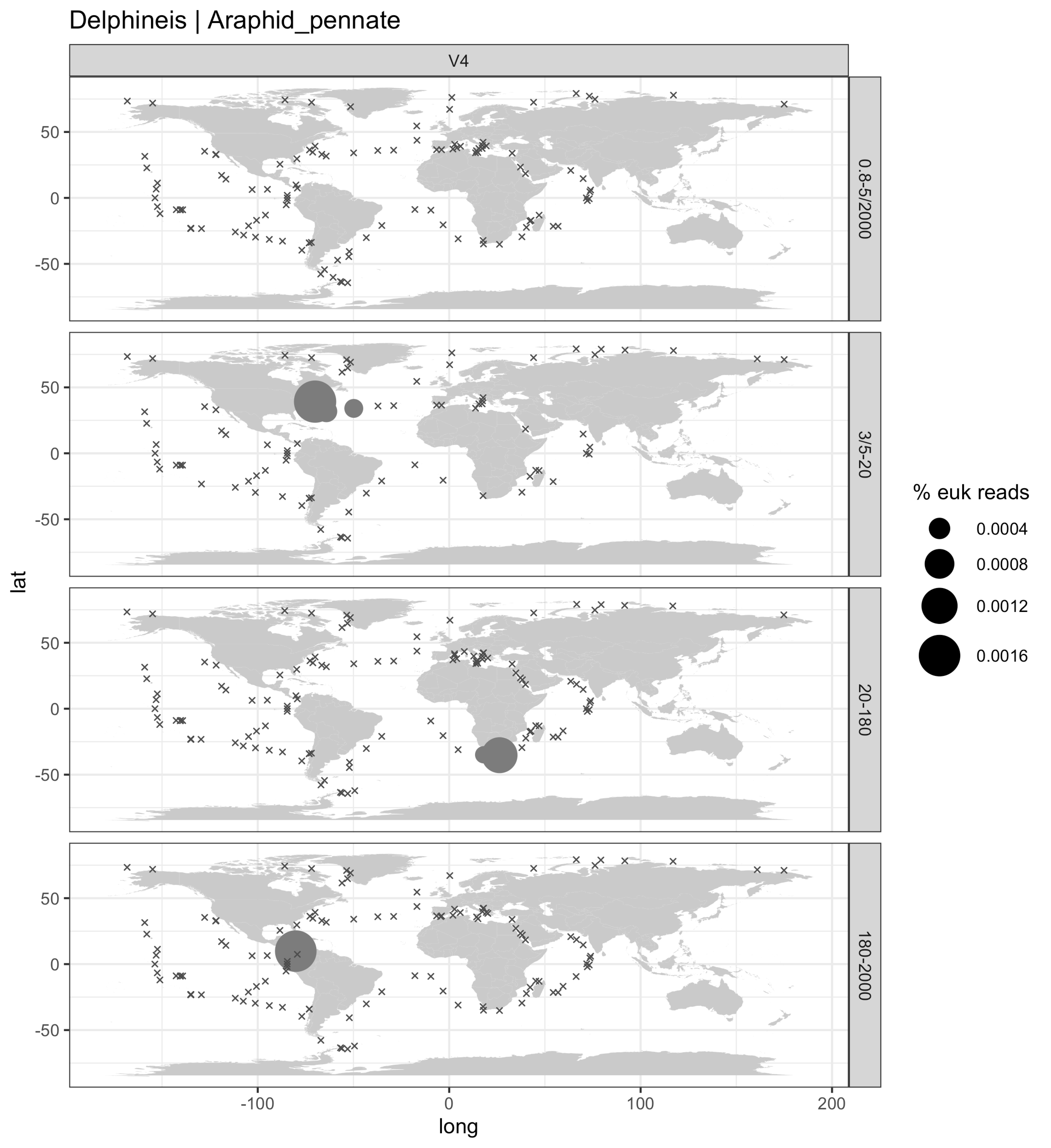

Delphineis | Araphid_pennate
V4
0.8-5/2000
3/5-20
20-180
180-2000
lat
long
50
0
-50
-100
0
100
200
% euk reads
0.0004
0.0008
0.0012
0.0016

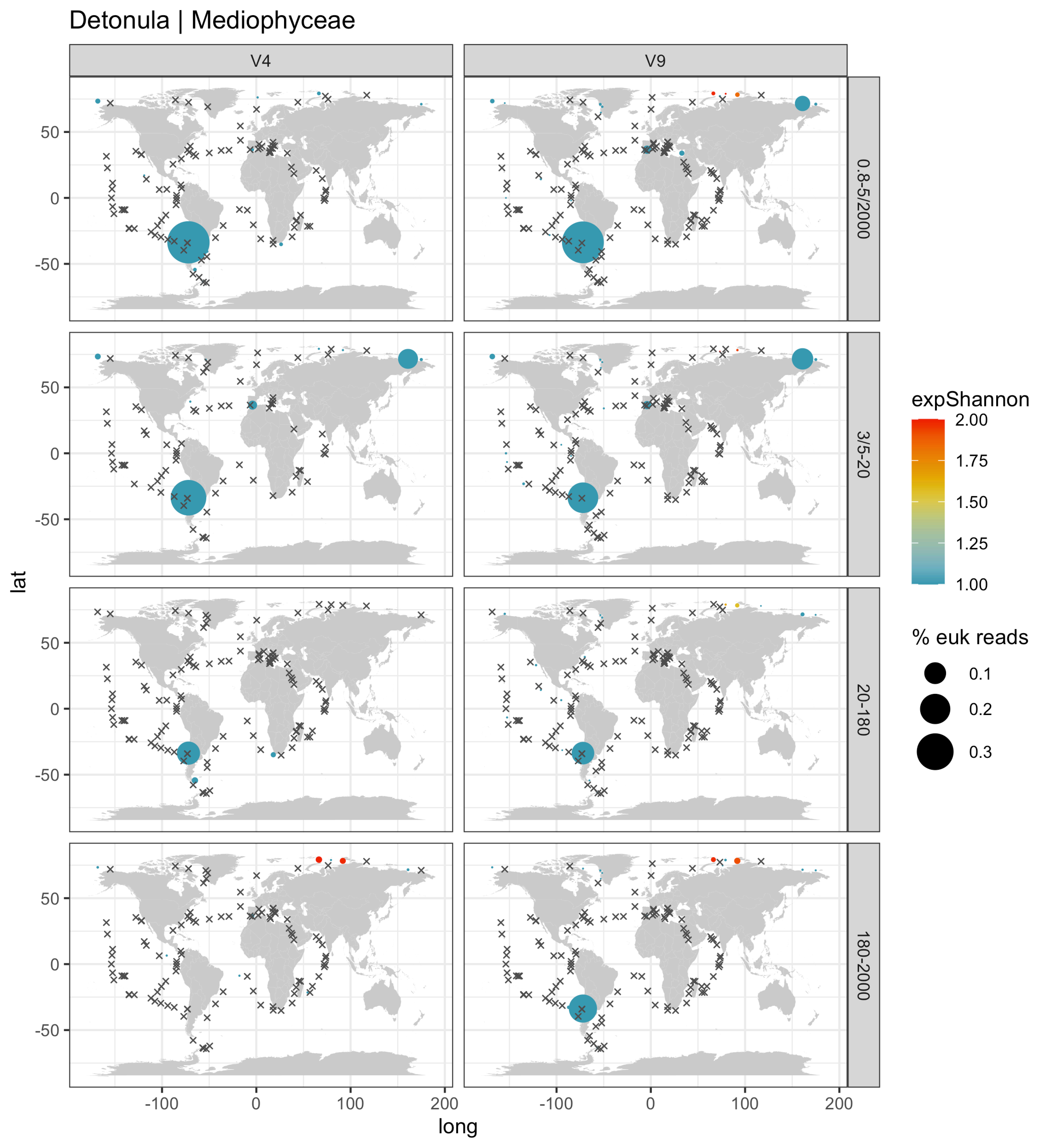

Detonula | Mediophyceae
V4
V9
0.8-5/2000
3/5-20
20-180
180-2000
lat
long
50
0
-50
-100
100
200
expShannon
2.00
1.75
1.50
1.25
1.00
% euk reads
0.1
0.2
0.3

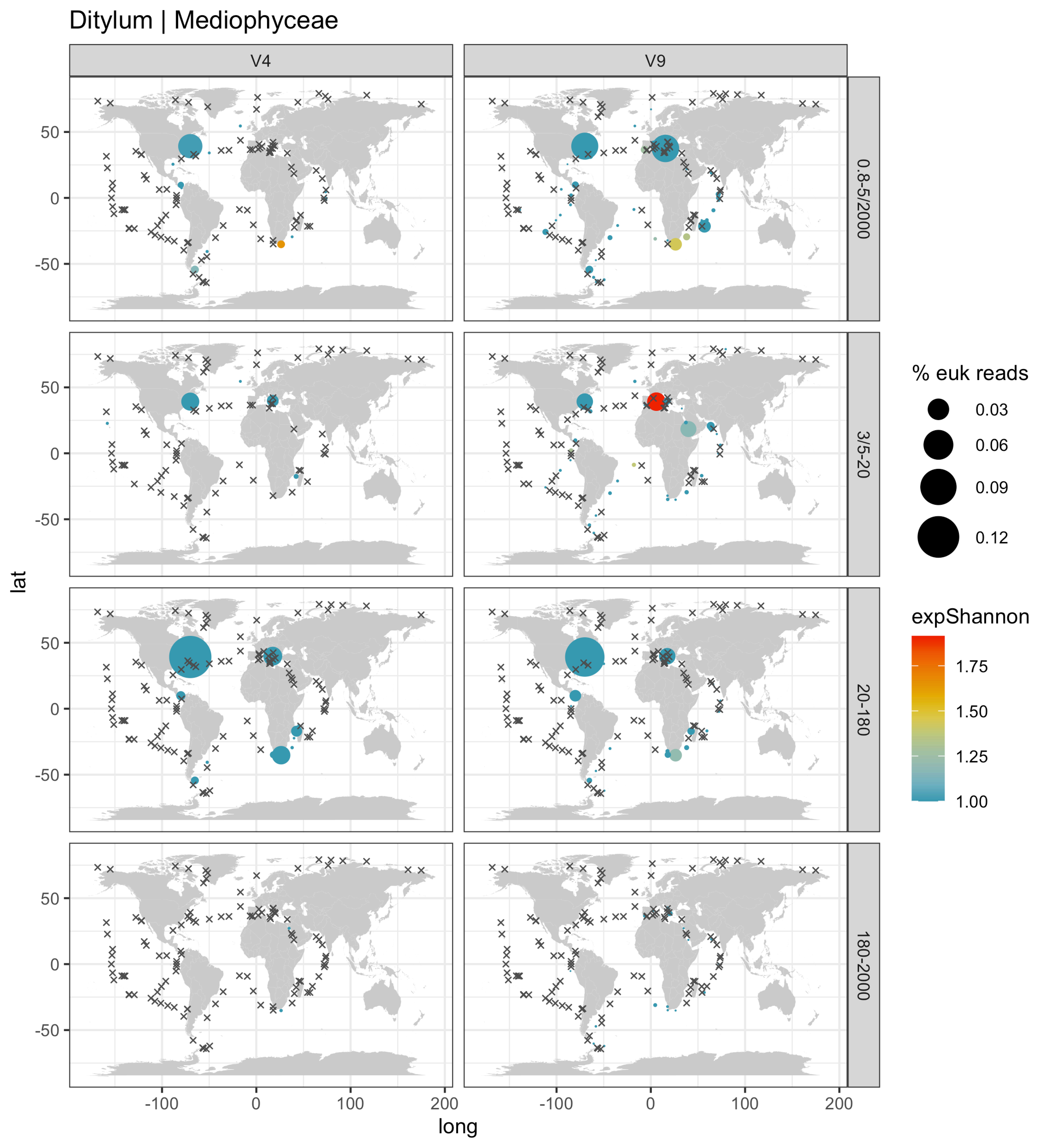

Ditylum | Mediophyceae
V4
V9
0.8-5/2000
3/5-20
20-180
180-2000
lat
long
50
0
-50
-100
0
100
200
% euk reads
0.03
0.06
0.09
0.12
expShannon
1.75
1.50
1.25
1.00

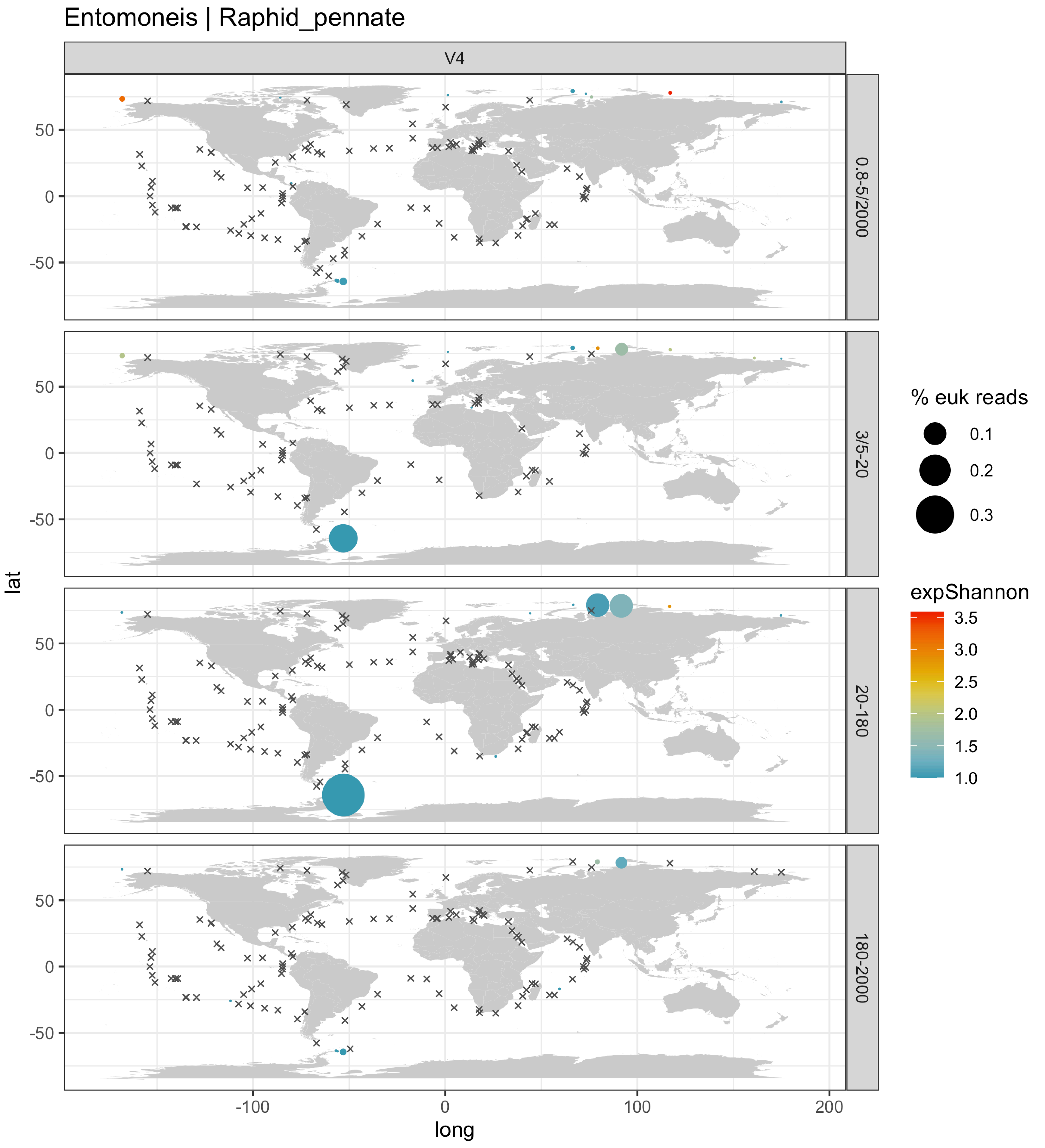
Entomoneis | Raphid_pennate
V4
0.8-5/2000
3/5-20
20-180
180-2000
lat
long
% euk reads
0.1
0.2
0.3
expShannon
3.5
3.0
2.5
2.0
1.5
1.0
50
0
-50
-100
0
100
200

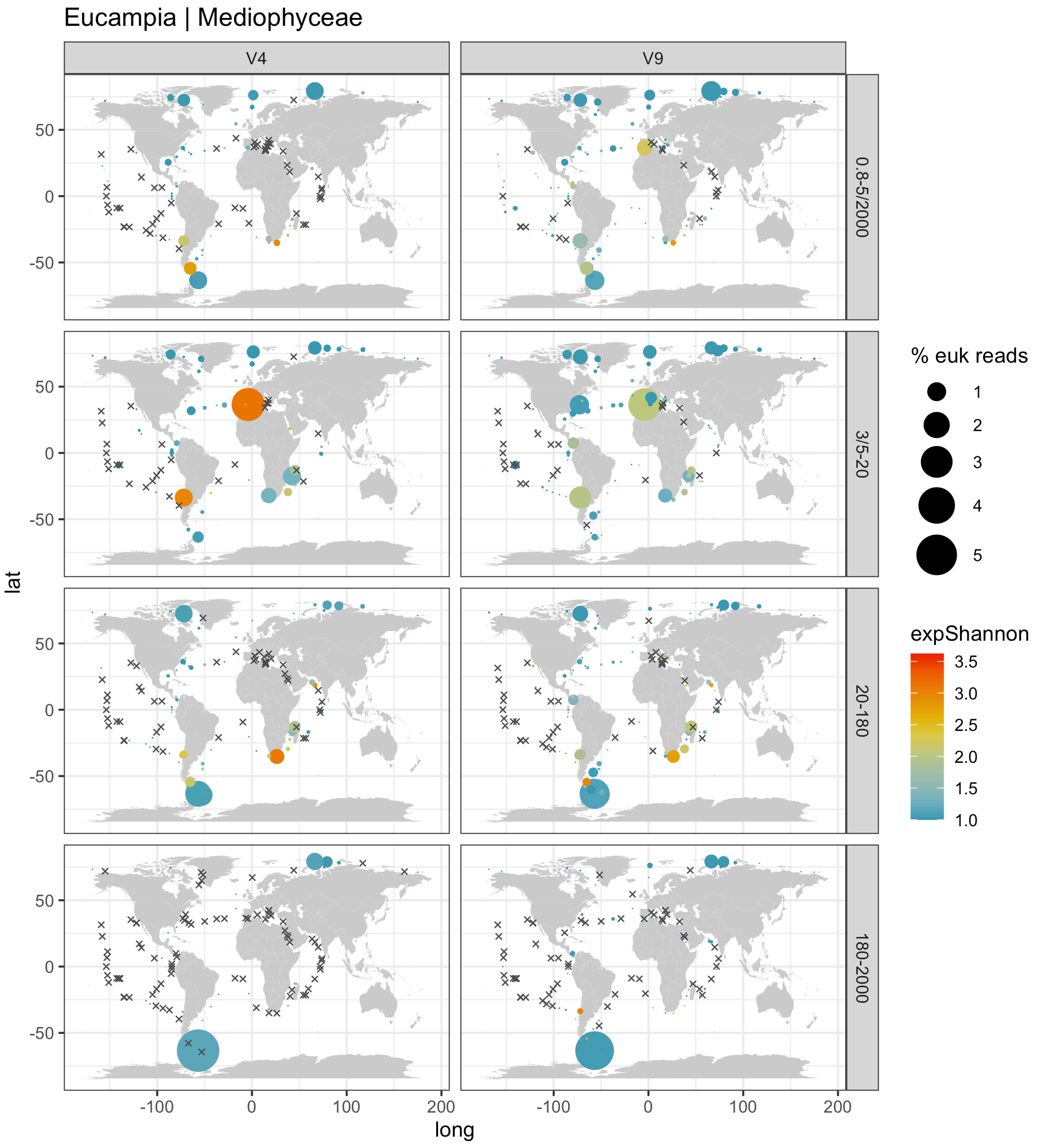

Eucampia | Mediophyceae
V4
V9
0.8-5/2000
3/5-20
20-180
180-2000
lat
long
50
0
-50
-100
0
100
200
% euk reads
1
2
3
4
5
expShannon
3.5
3.0
2.5
2.0
1.5
1.0

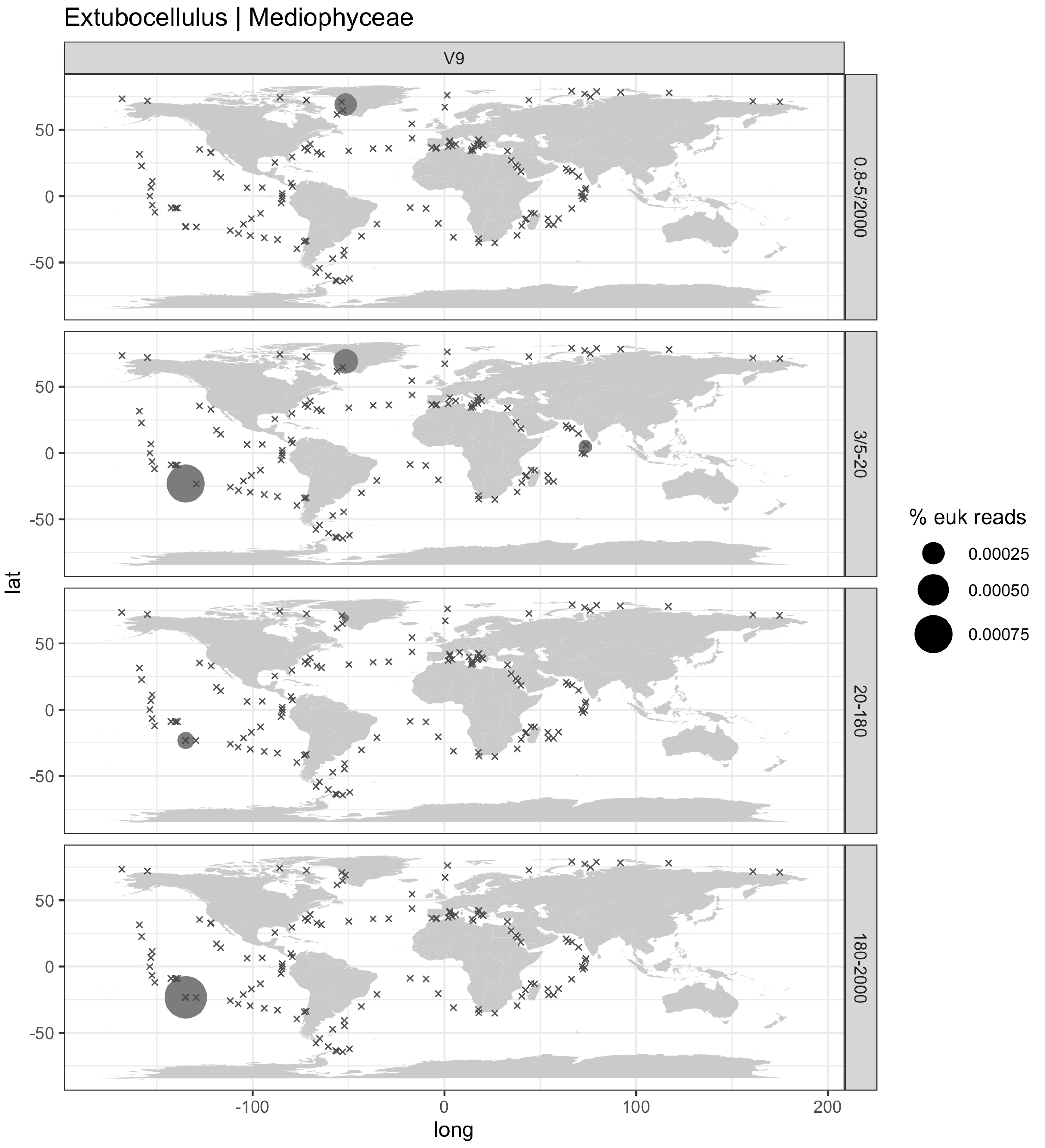

Extubocellulus | Mediophyceae
V9
0.8-5/2000
3/5-20
20-180
180-2000
lat
long
50
0
-50
-100
100
200
% euk reads
0.00025
0.00050
0.00075

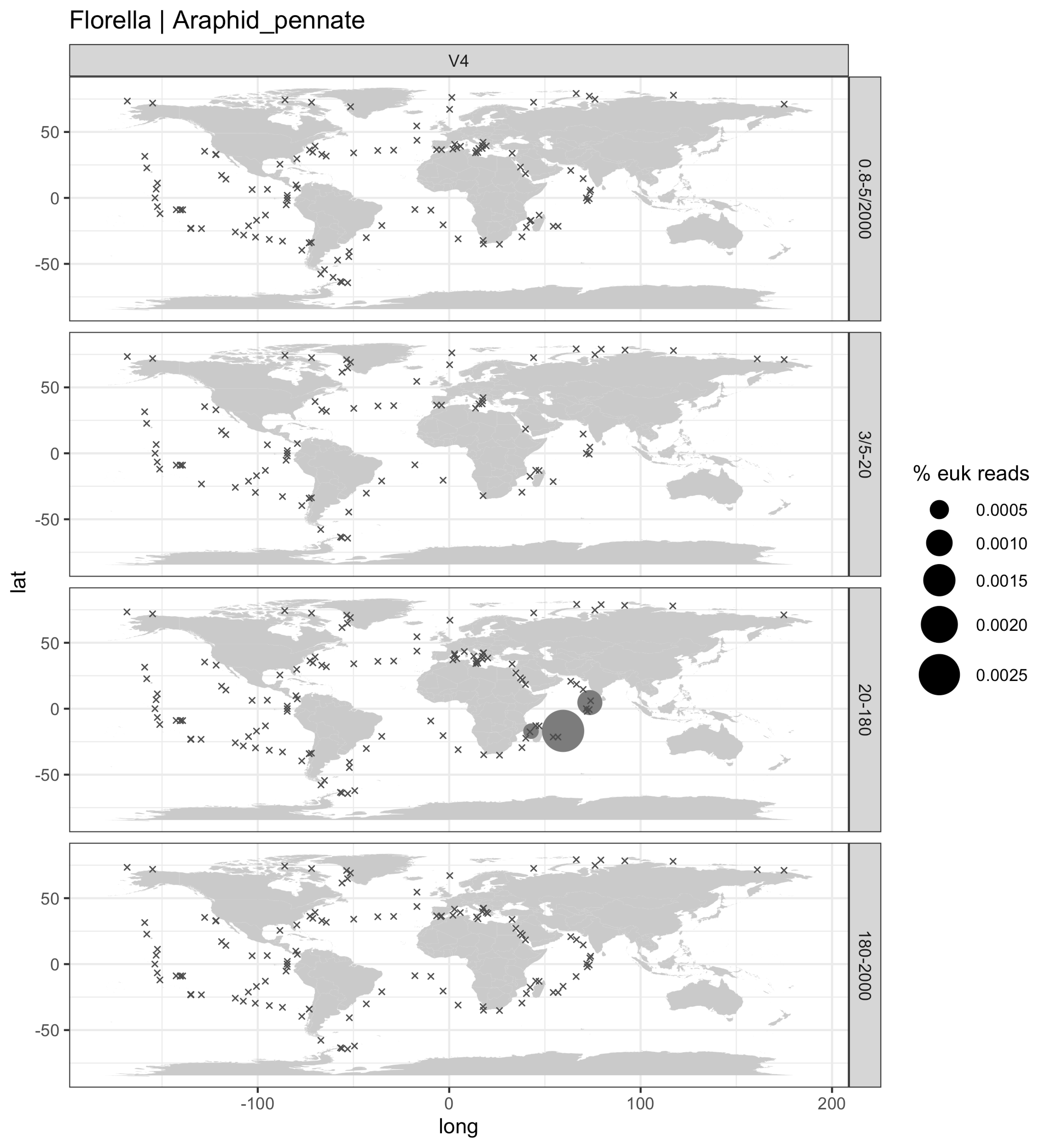

Florella | Araphid_pennate
V4
0.8-5/2000
3/5-20
20-180
180-2000
lat
long
% euk reads
0.0005
0.0010
0.0015
0.0020
0.0025
50
0
-50
-100
0
100
200

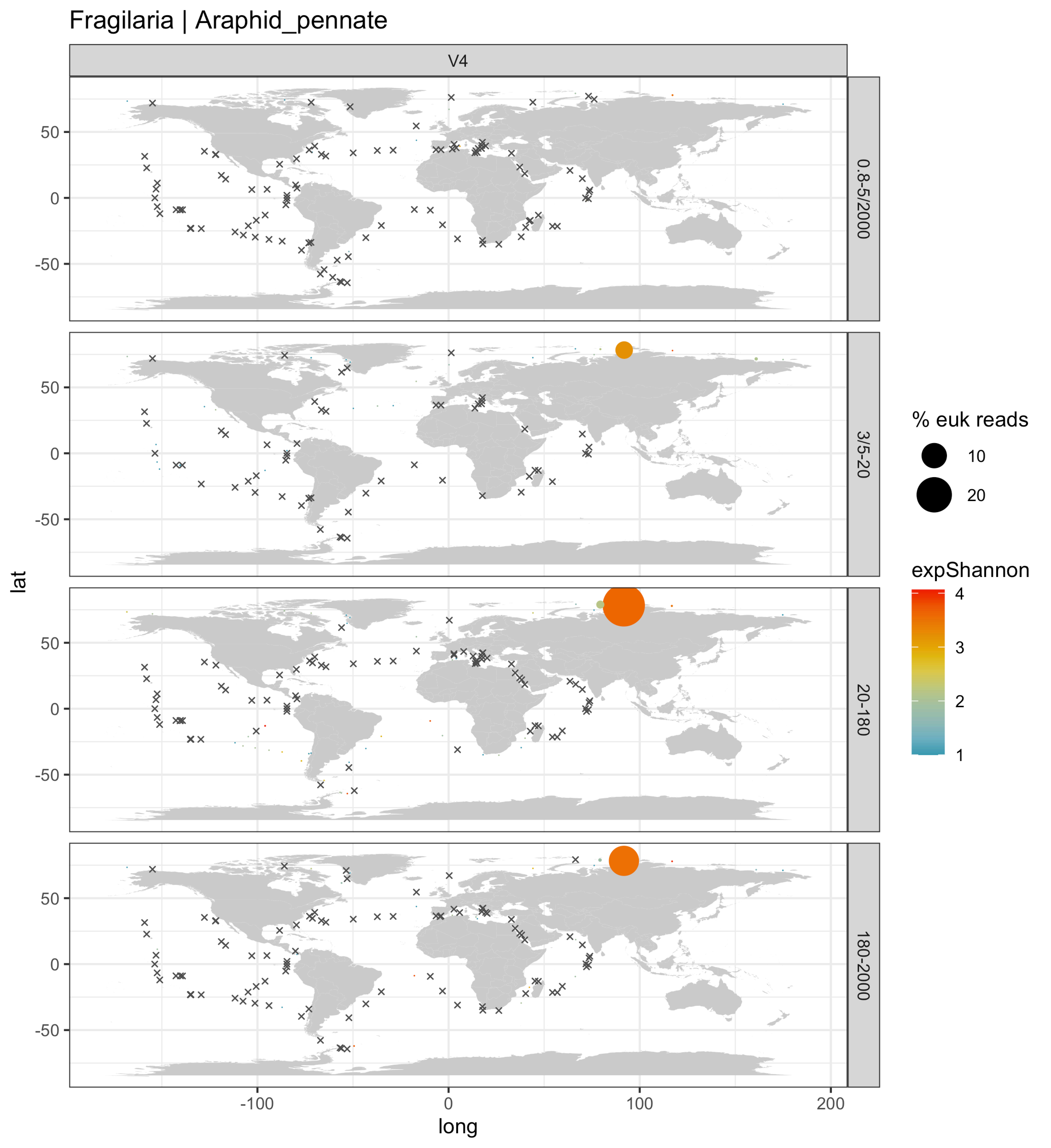
Fragilaria | Araphid_pennate
V4
0.8-5/2000
3/5-20
20-180
180-2000
lat
long
50
0
-50
-100
100
200
% euk reads
10
20
expShannon
4
3
2
1

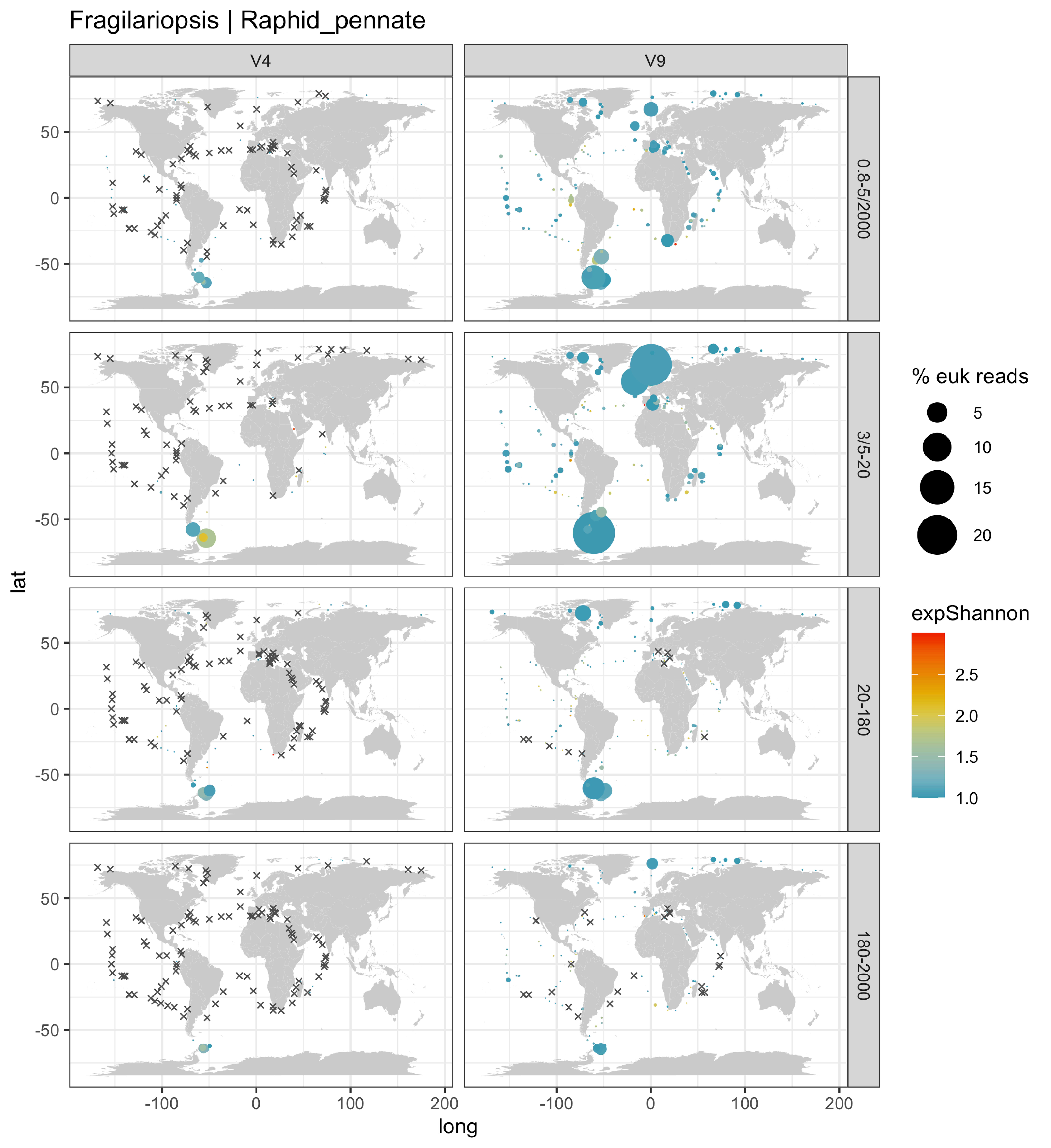
Fragilariopsis | Raphid_pennate
V4
V9
0.8-5/2000
3/5-20
20-180
180-2000
lat
long
% euk reads
5
10
15
20
expShannon
2.5
2.0
1.5
1.0
50
0
-50
-100
100
200

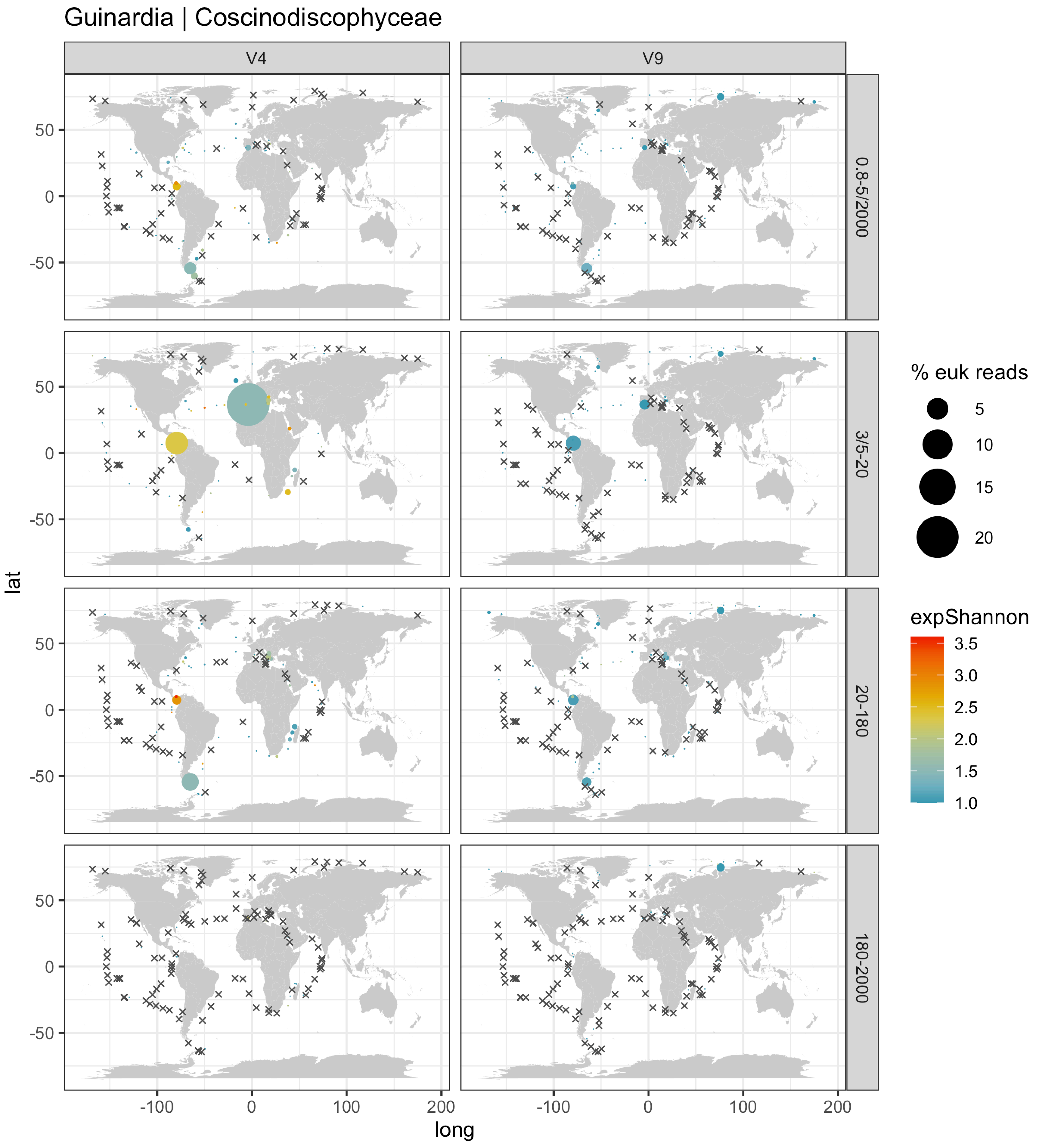
Guinardia | Coscinodiscophyceae
V4
V9
0.8-5/2000
3/5-20
20-180
180-2000
% euk reads
5
10
15
20
expShannon
3.5
3.0
2.5
2.0
1.5
1.0
lat
long
50
0
-50
-100
0
100
200

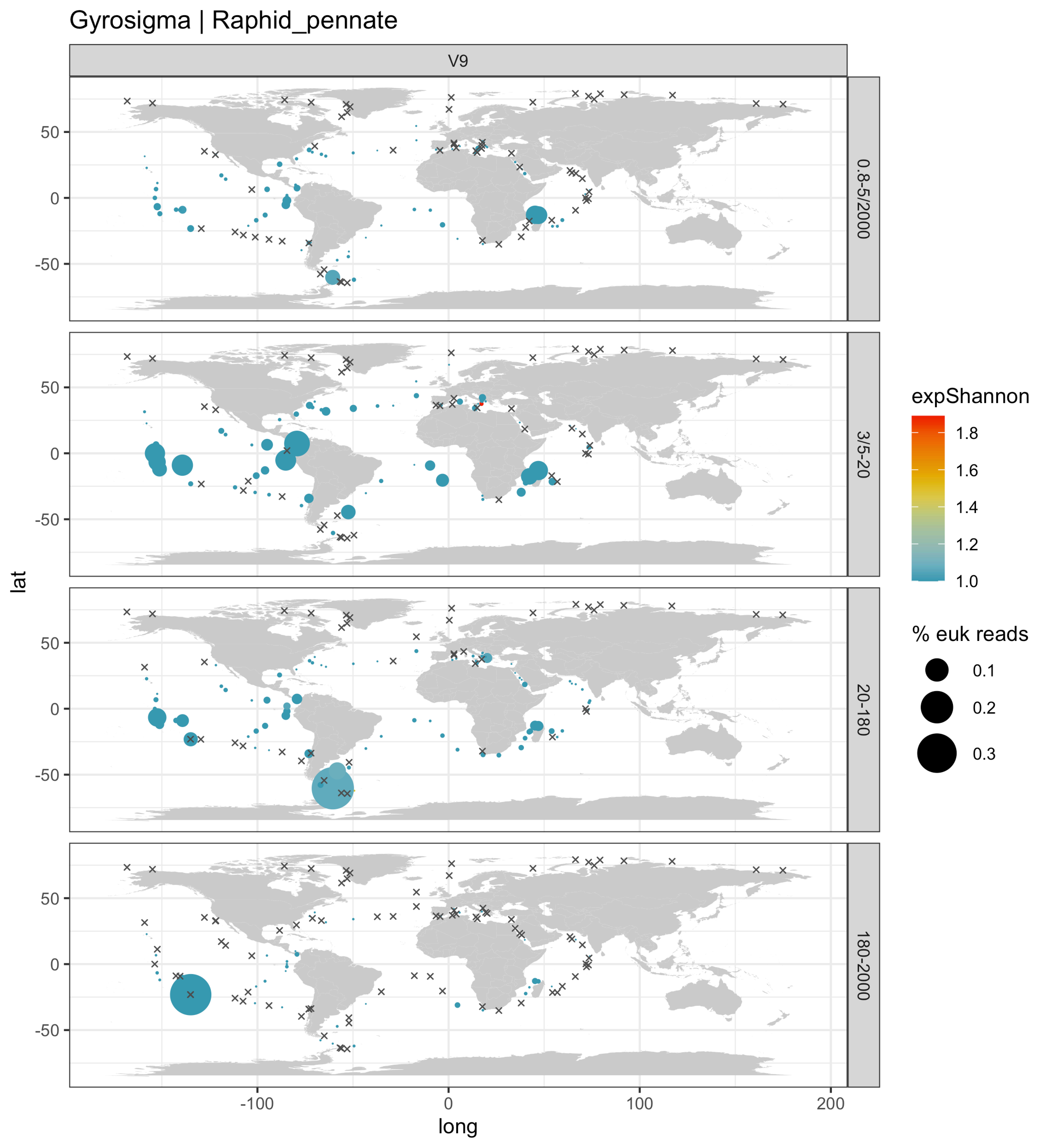

Gyrosigma | Raphid_pennate
V9
0.8-5/2000
3/5-20
20-180
180-2000
lat
long
50
0
-50
-100
0
100
200
expShannon
1.8
1.6
1.4
1.2
1.0
% euk reads
0.1
0.2
0.3

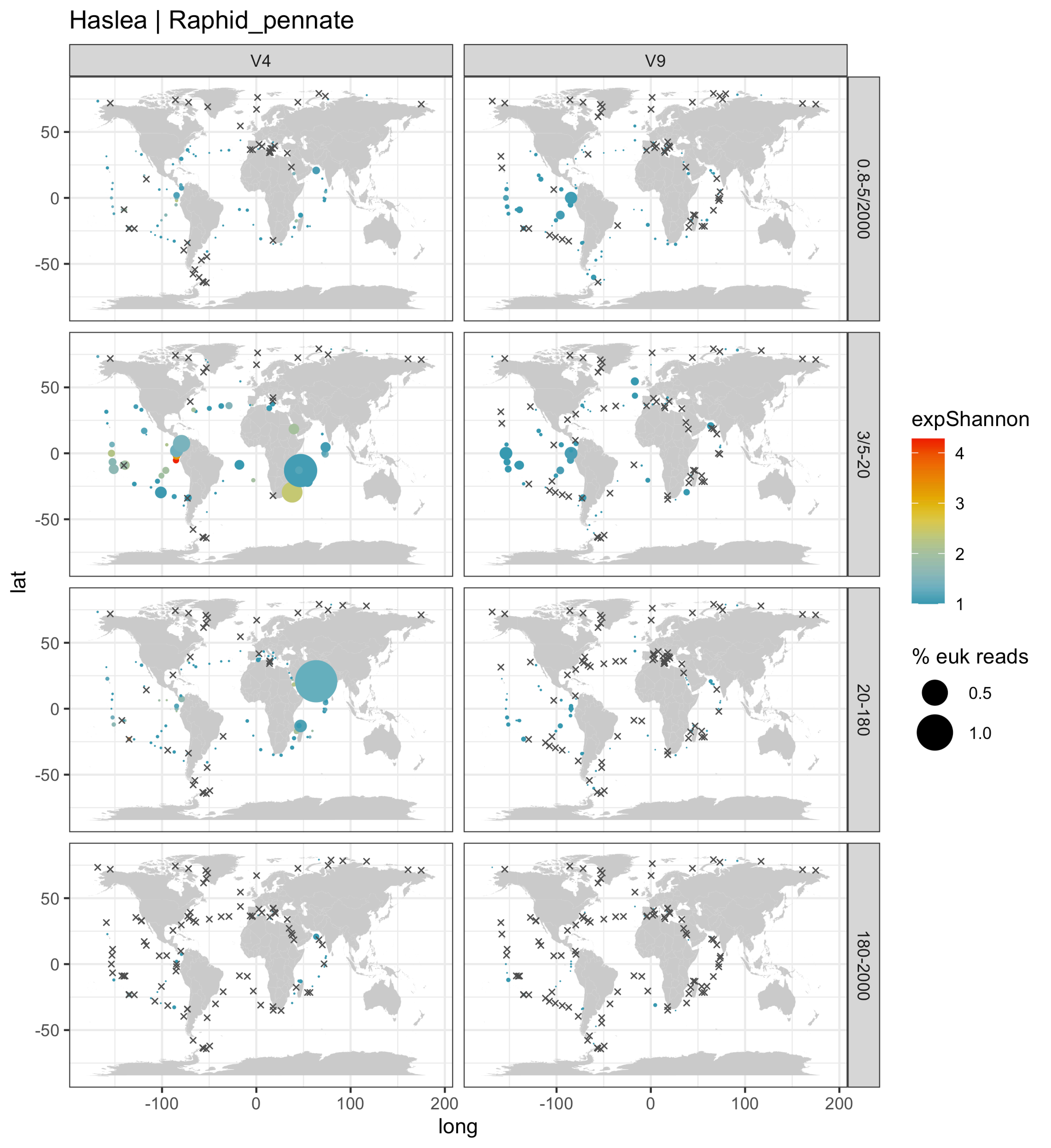

Haslea | Raphid_pennate
V4
V9
0.8-5/2000
3/5-20
20-180
180-2000
expShannon
4
3
2
1
% euk reads
0.5
1.0
lat
long
50
0
-50
-100
0
100
200

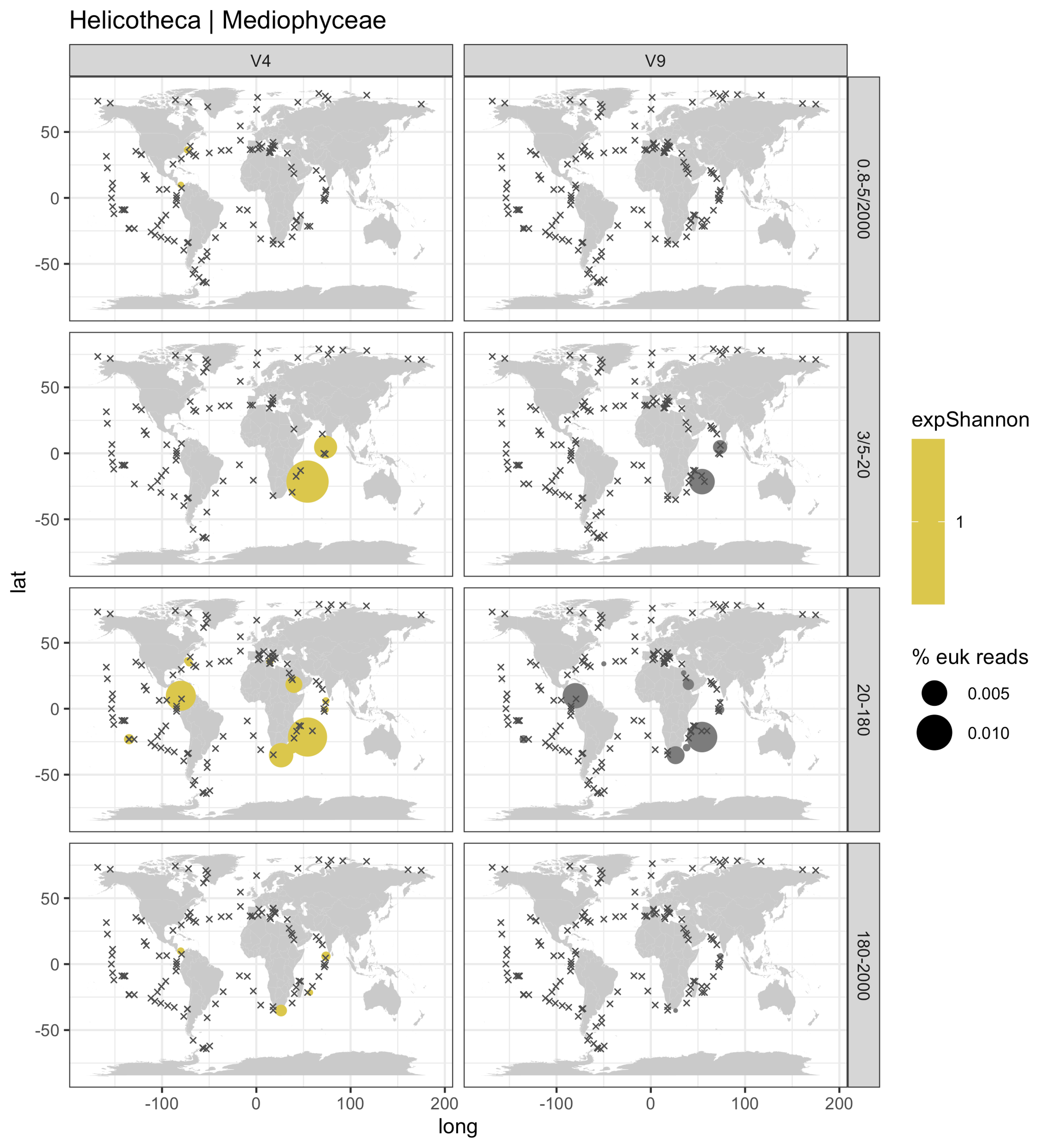

Helicotheca | Mediophyceae
V4
V9
0.8-5/2000
3/5-20
20-180
180-2000
lat
long
50
0
-50
-100
0
100
200
expShannon
1
% euk reads
0.005
0.010

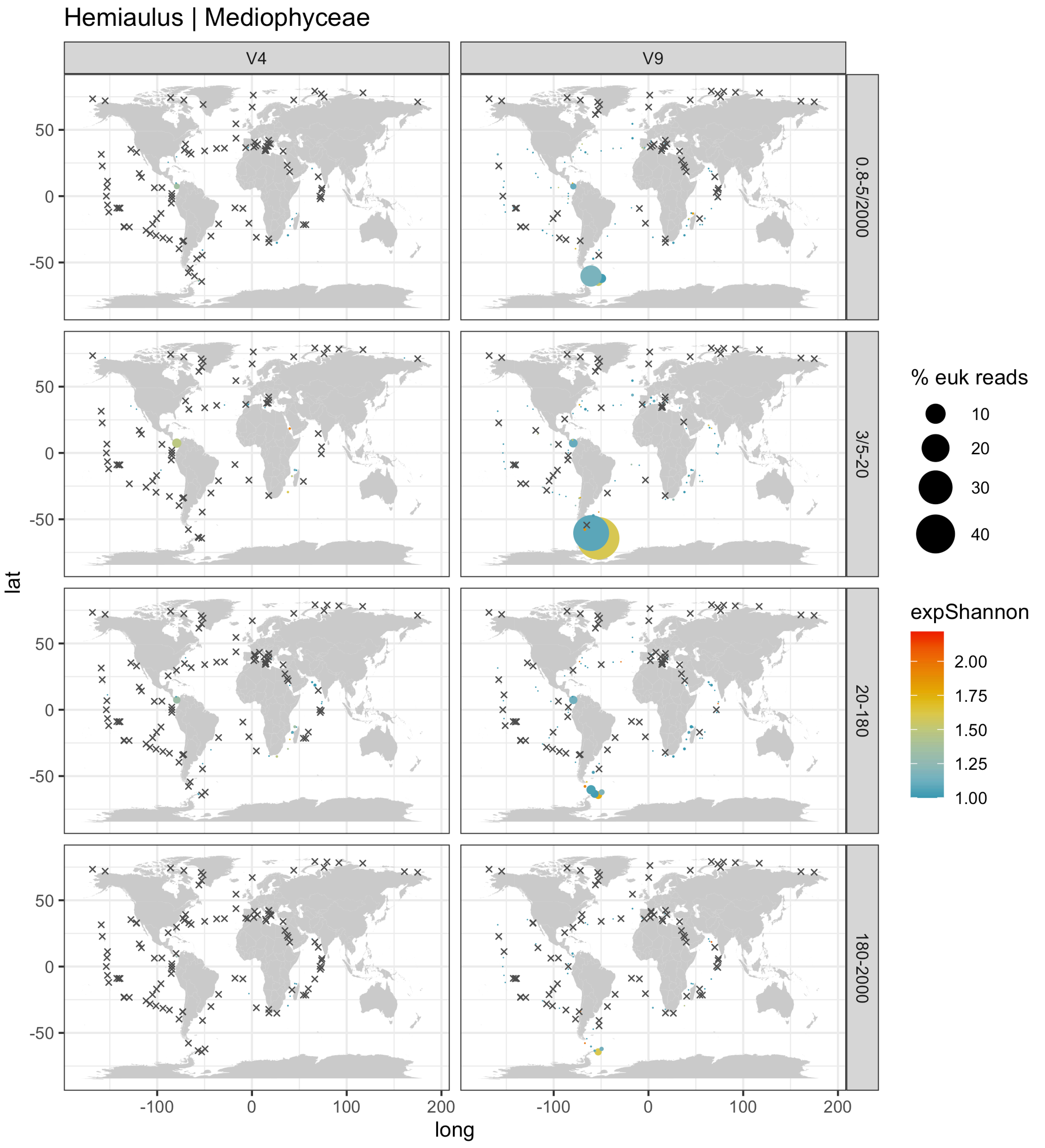
Hemiaulus | Mediophyceae
V4
V9
0.8-5/2000
3/5-20
20-180
180-2000
% euk reads
10
20
30
40
expShannon
2.00
1.75
1.50
1.25
1.00
lat
long
50
0
-50
-100
0
100
200

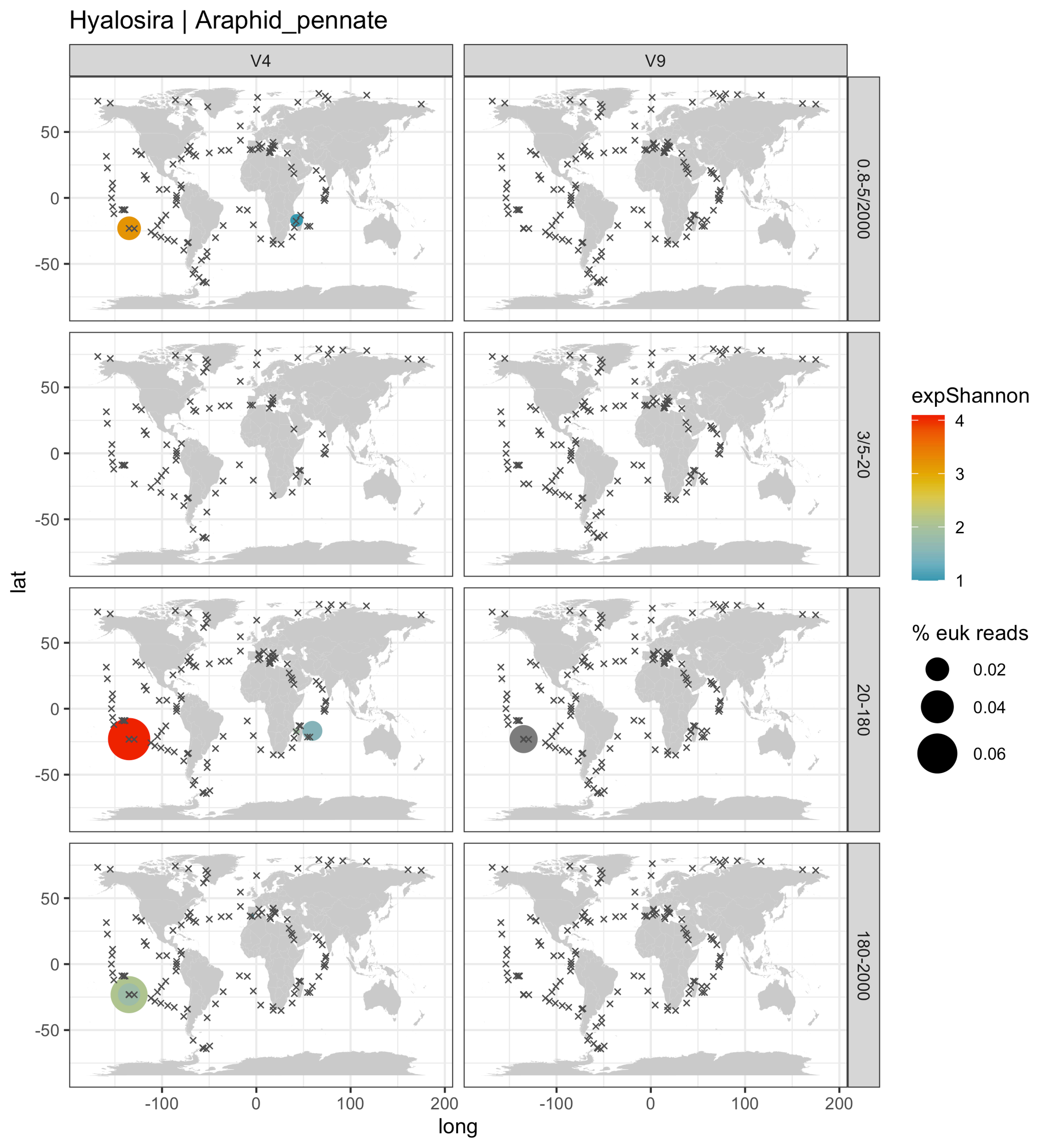

Hyalosira | Araphid_pennate
V4
V9
0.8-5/2000
3/5-20
20-180
180-2000
lat
long
50
0
-50
-100
100
200
expShannon
4
3
2
1
% euk reads
0.02
0.04
0.06

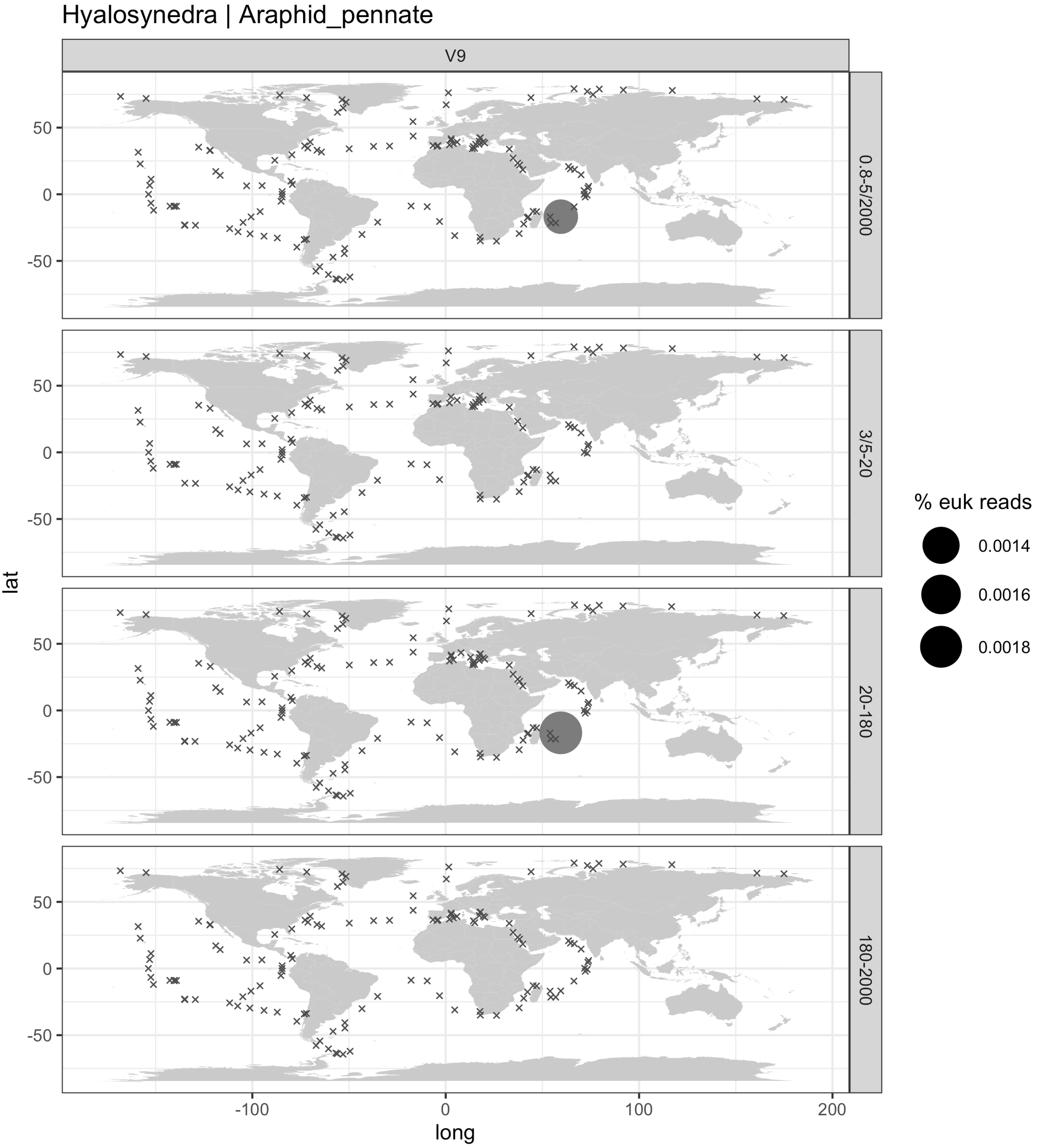

Hyalosynedra | Araphid_pennate
V9
0.8-5/2000
3/5-20
20-180
180-2000
lat
long
50
0
-50
-100
0
100
200
% euk reads
0.0014
0.0016
0.0018

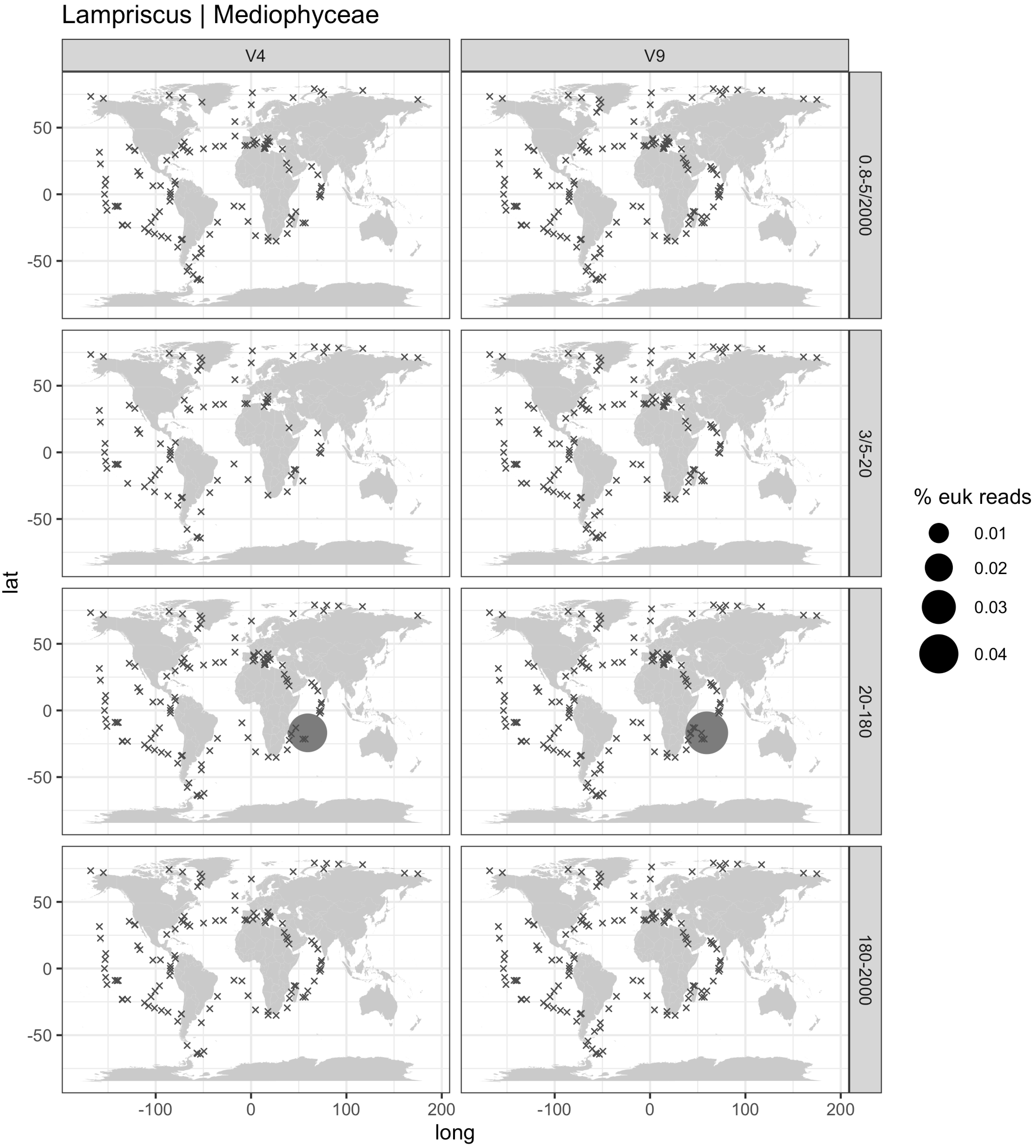

Lampriscus | Mediophyceae
V4
V9
0.8-5/2000
3/5-20
20-180
180-2000
lat
long
50
0
-50
-100
0
100
200
% euk reads
0.01
0.02
0.03
0.04

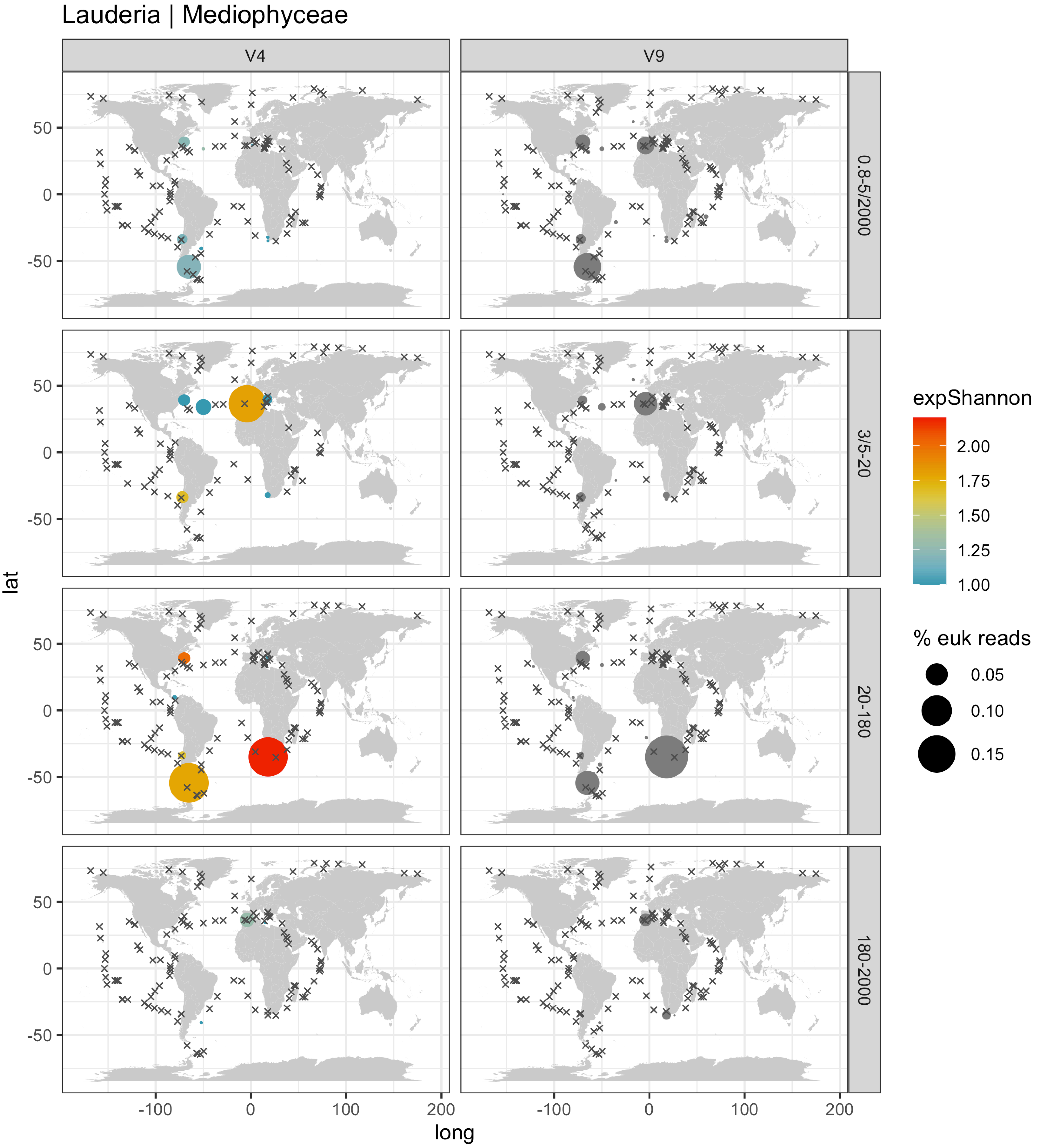

Lauderia | Mediophyceae
V4
V9
0.8-5/2000
3/5-20
20-180
180-2000
lat
long
50
0
-50
-100
0
100
200
expShannon
2.00
1.75
1.50
1.25
1.00
% euk reads
0.05
0.10
0.15

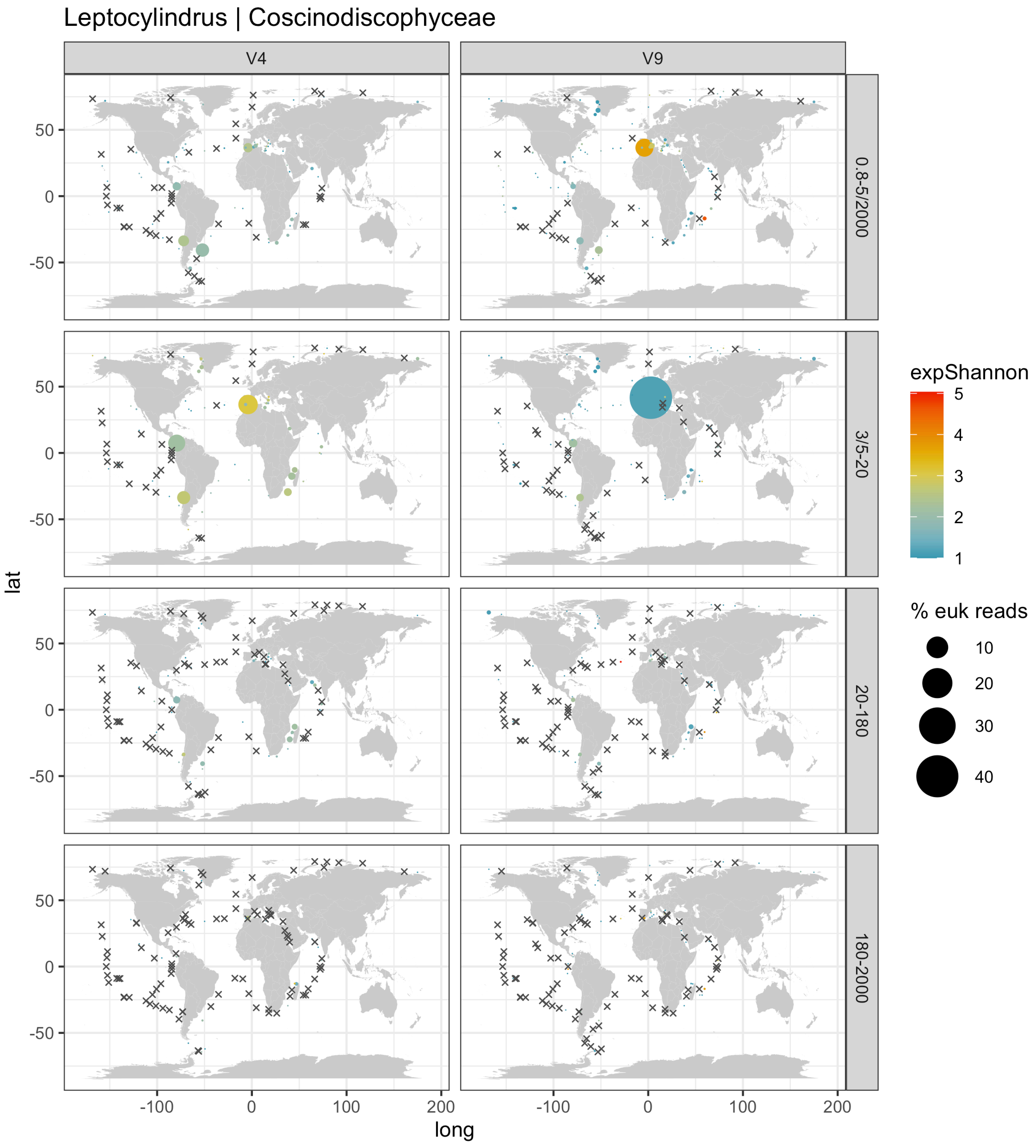

Leptocylindrus | Coscinodiscophyceae
V4
V9
0.8-5/2000
3/5-20
20-180
180-2000
lat
long
50
0
-50
-100
100
200
expShannon
5
4
3
2
1
% euk reads
10
20
30
40

Licmophora | Araphid_pennate

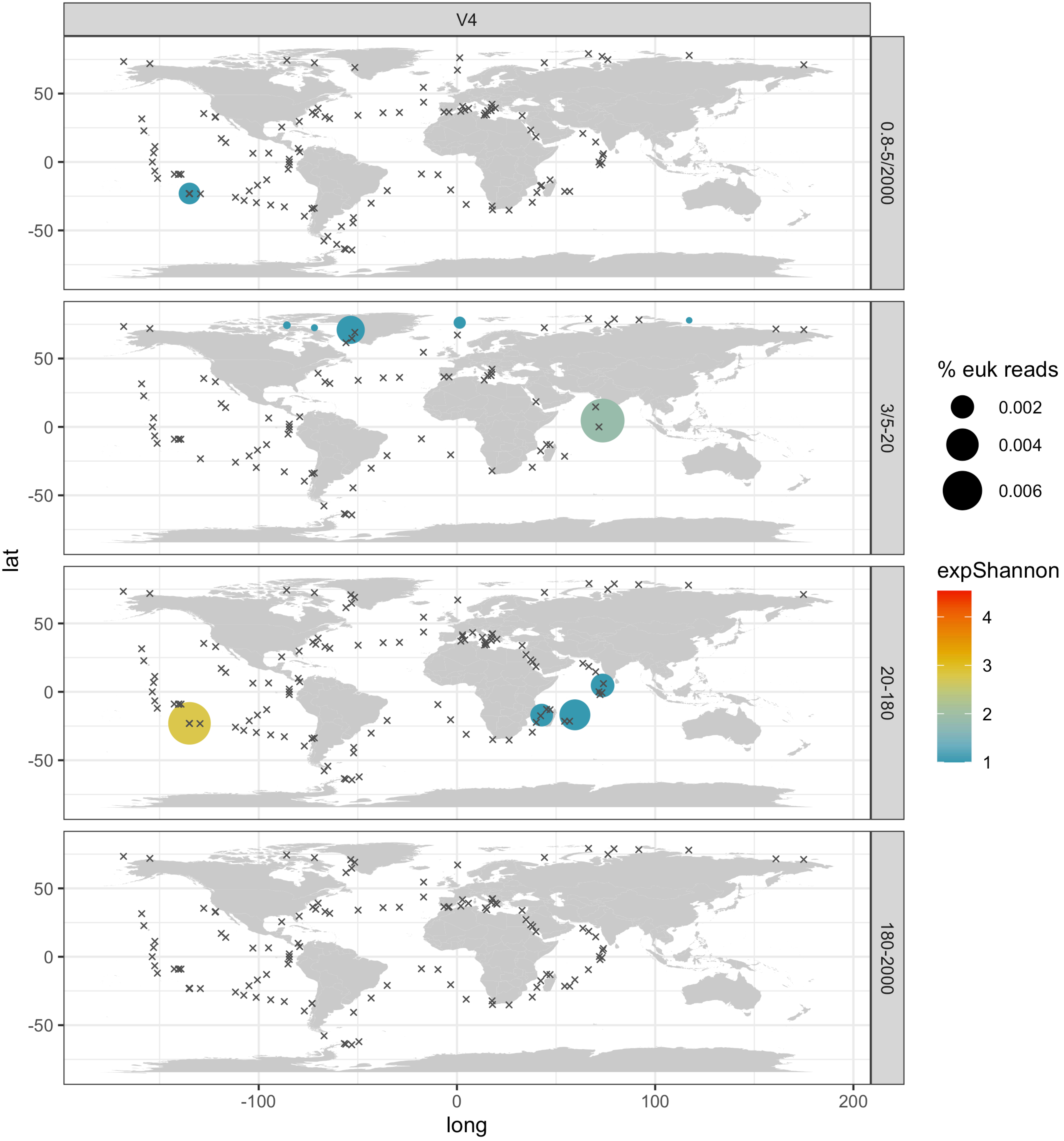

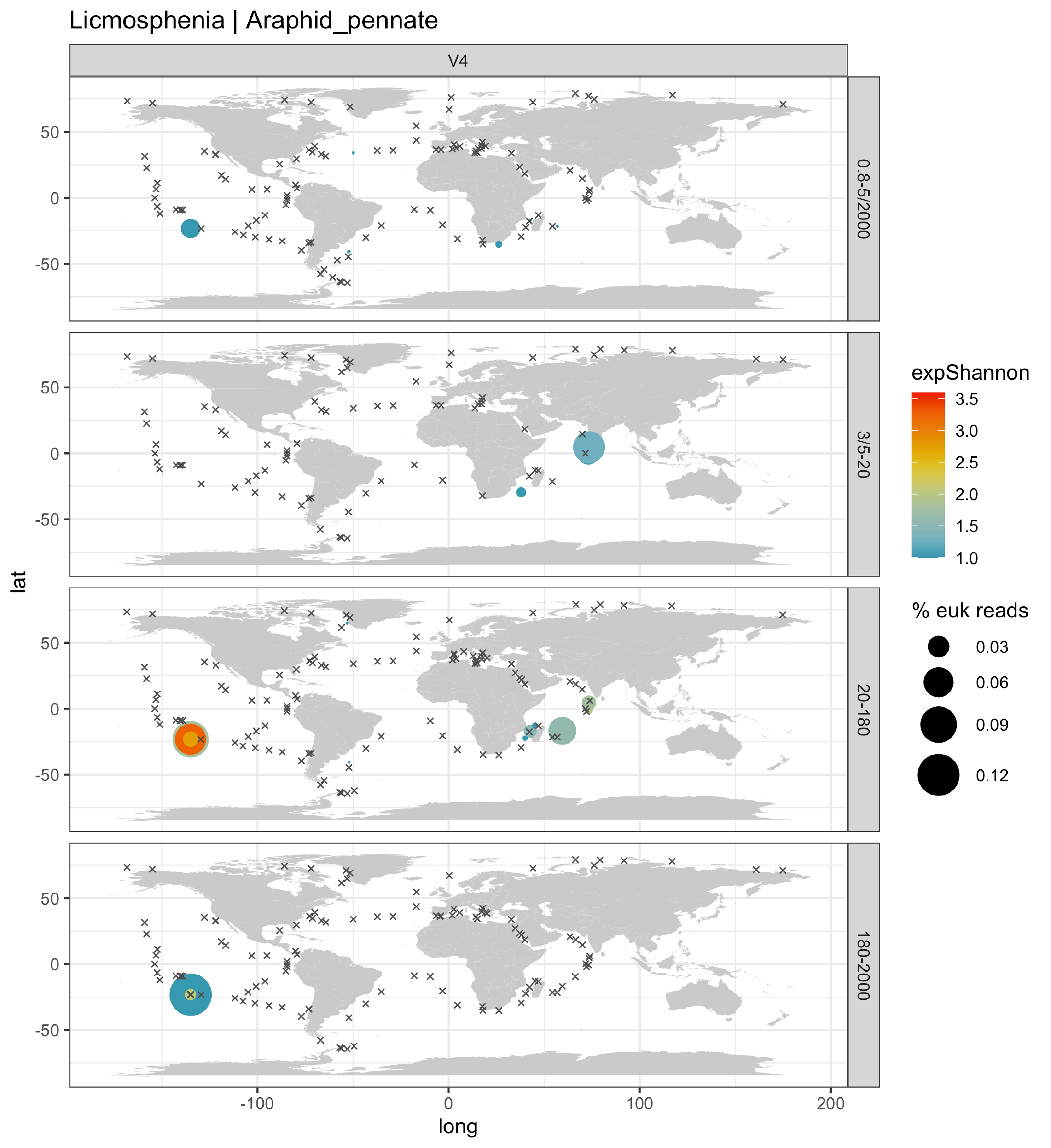

Licmosphenia | Araphid_pennate
V4
0.8-5/2000
3/5-20
20-180
180-2000
lat
long
50
0
-50
-100
0
100
200
expShannon
3.5
3.0
2.5
2.0
1.5
1.0
% euk reads
0.03
0.06
0.09
0.12

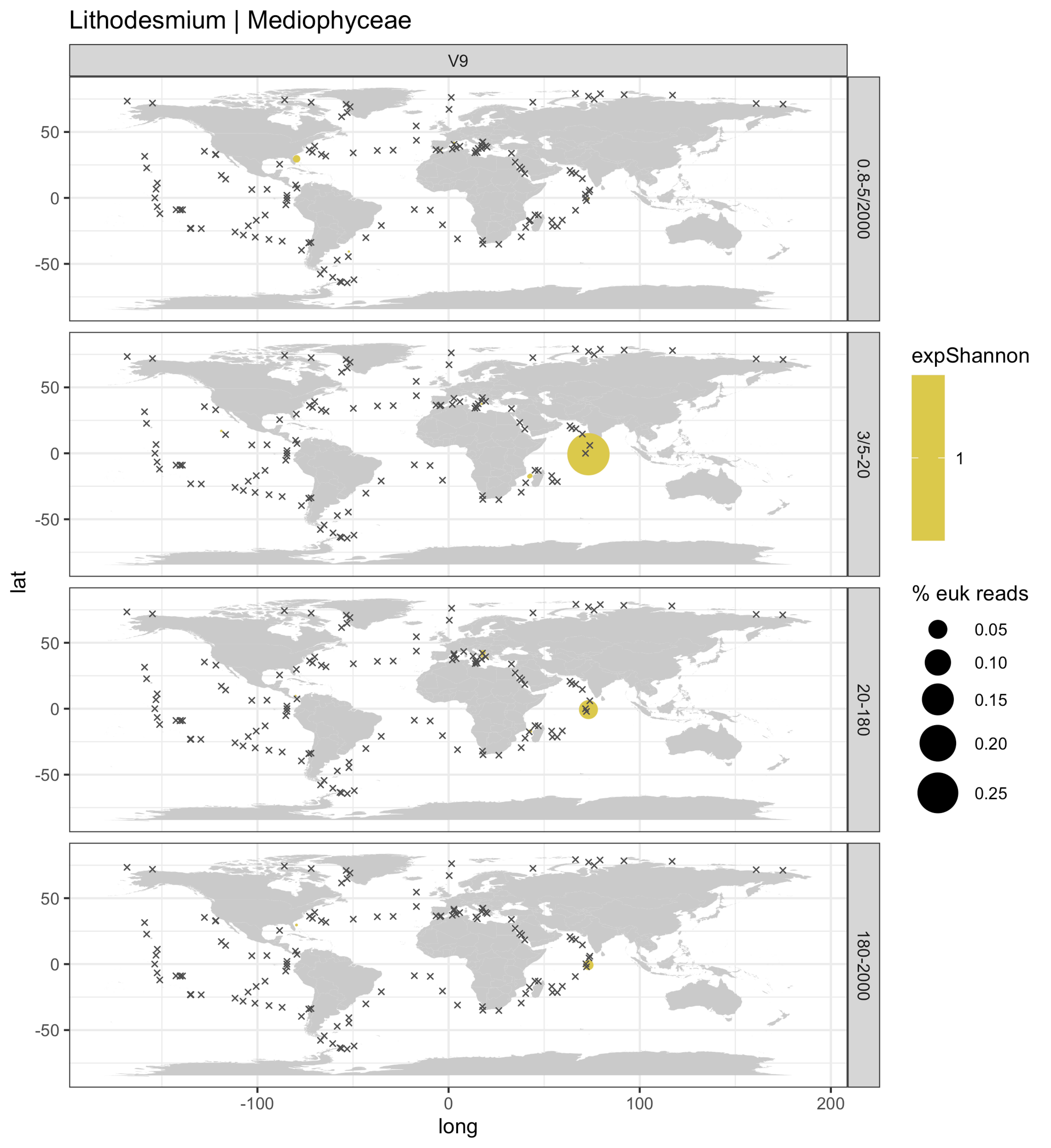

Lithodesmium | Mediophyceae
V9
0.8-5/2000
3/5-20
20-180
180-2000
lat
long
50
0
-50
-100
100
200
expShannon
1
% euk reads
0.05
0.10
0.15
0.20
0.25

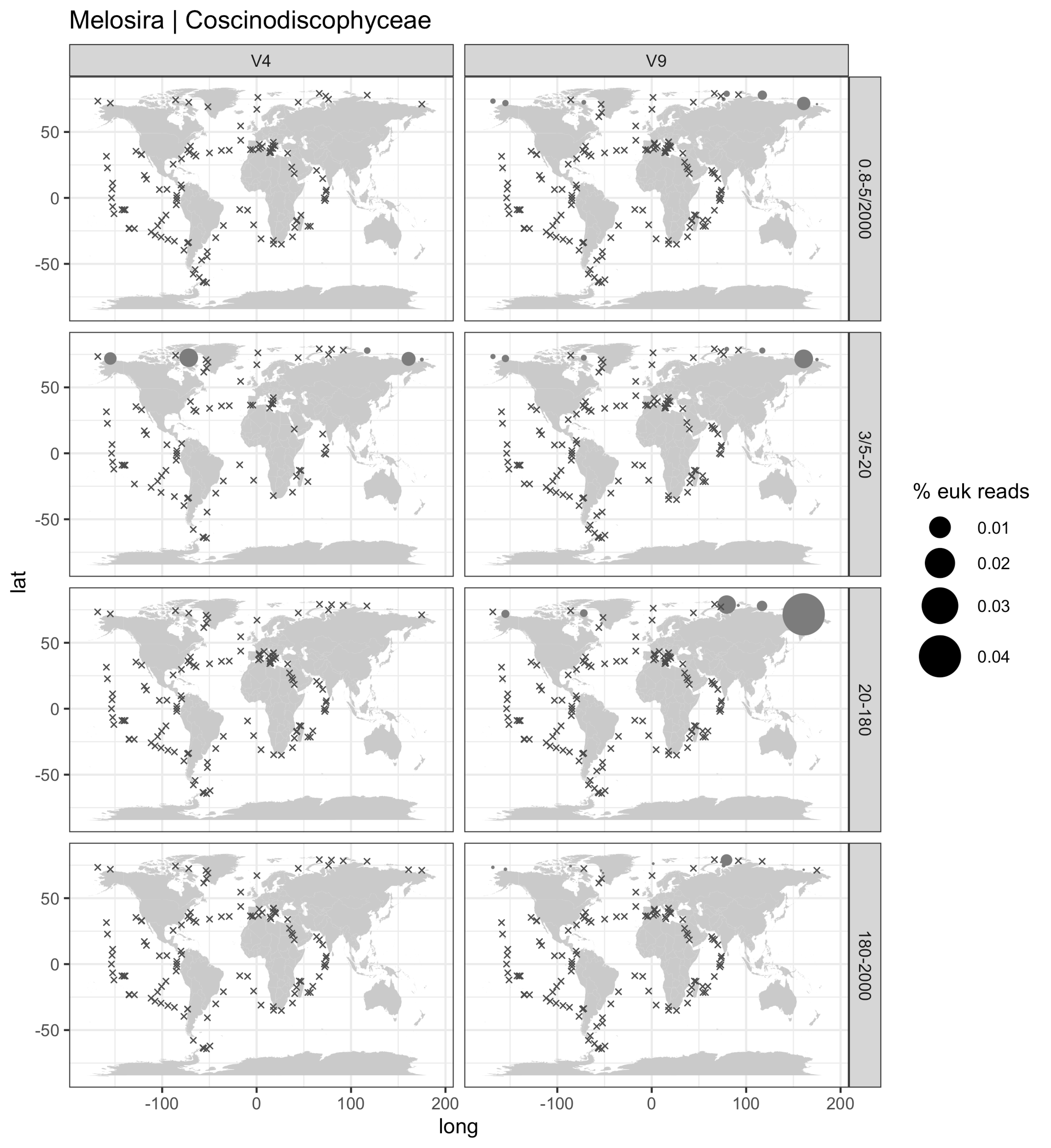

Melosira | Coscinodiscophyceae
V4
V9
0.8-5/2000
3/5-20
20-180
180-2000
lat
long
50
0
-50
-100
100
200
% euk reads
0.01
0.02
0.03
0.04

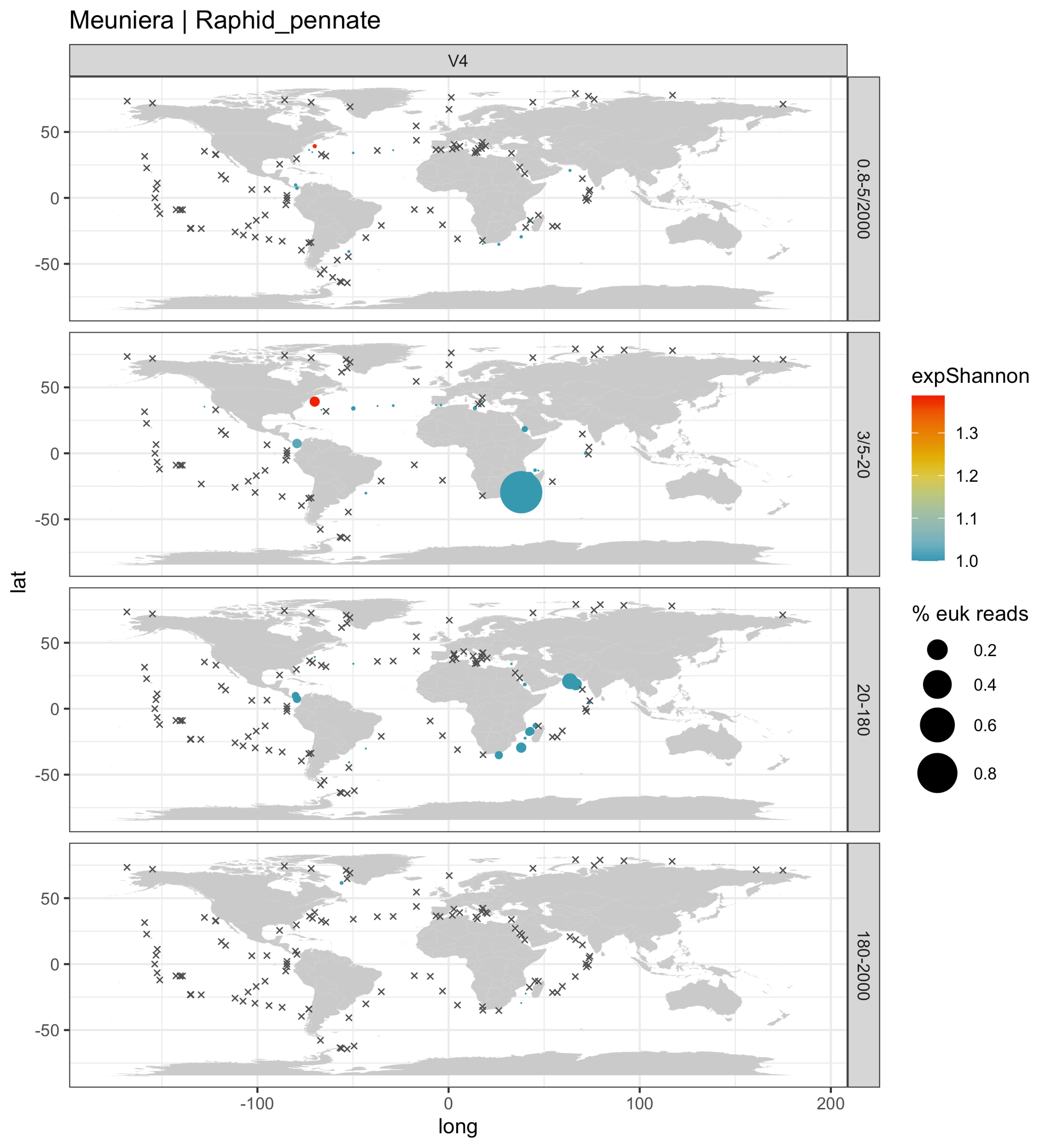

Meuniera | Raphid_pennate
V4
0.8-5/2000
3/5-20
20-180
180-2000
lat
long
50
0
-50
-100
0
100
200
expShannon
1.3
1.2
1.1
1.0
% euk reads
0.2
0.4
0.6
0.8

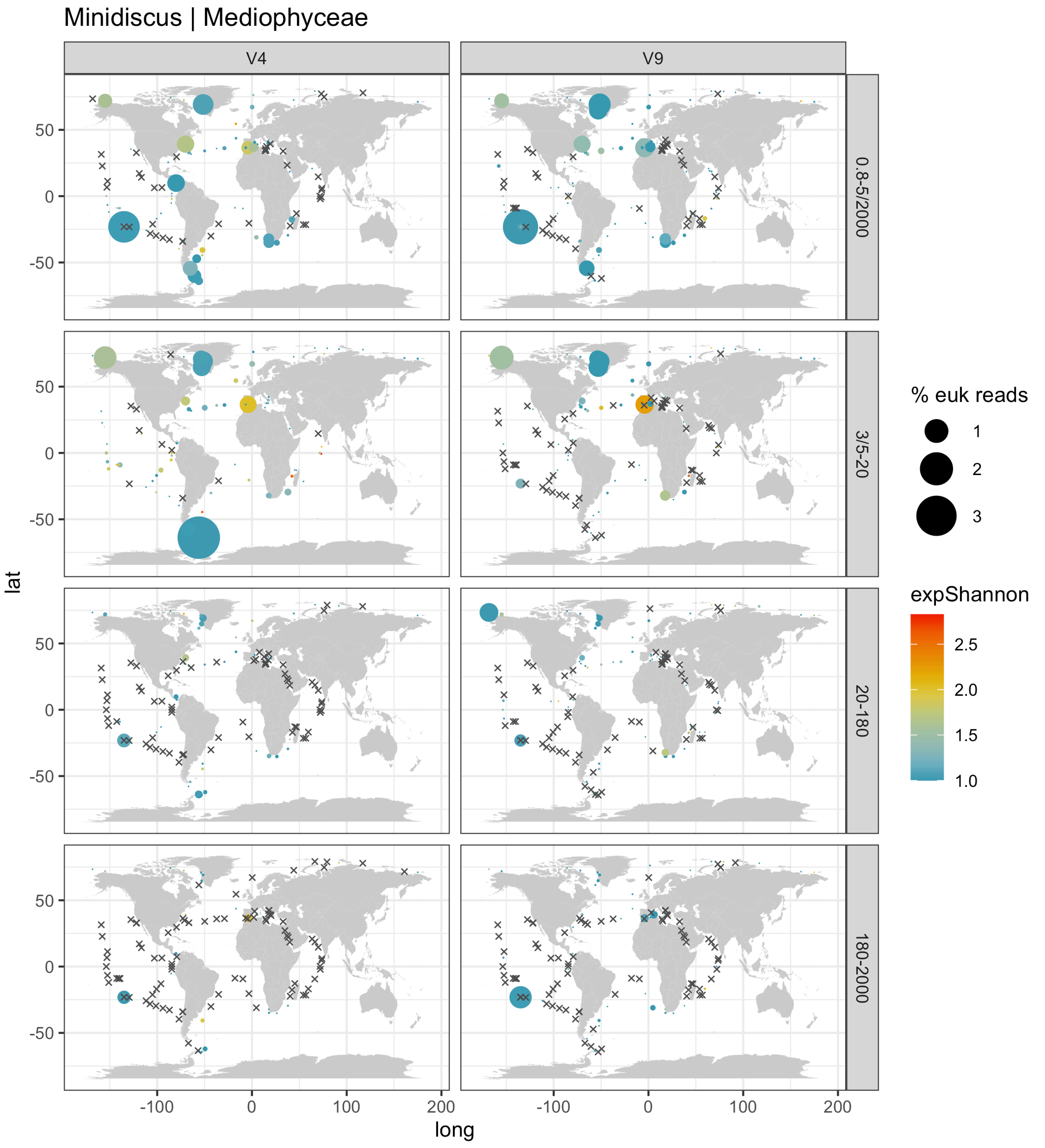

Minidiscus | Mediophyceae
V4
V9
0.8-5/2000
3/5-20
20-180
180-2000
lat
long
50
0
-50
-100
100
200
% euk reads
1
2
3
expShannon
2.5
2.0
1.5
1.0

Minutocellus | Mediophyceae

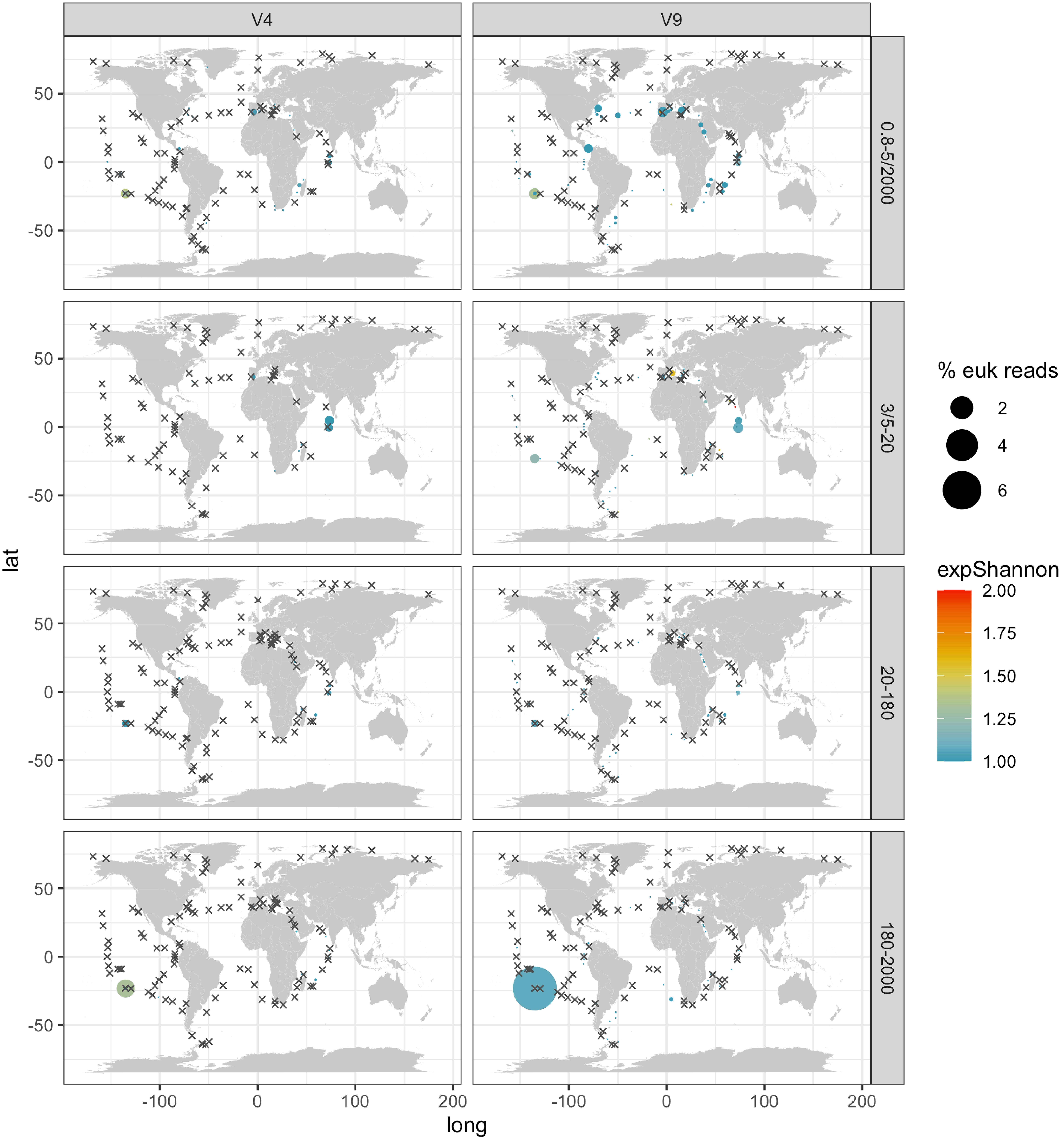

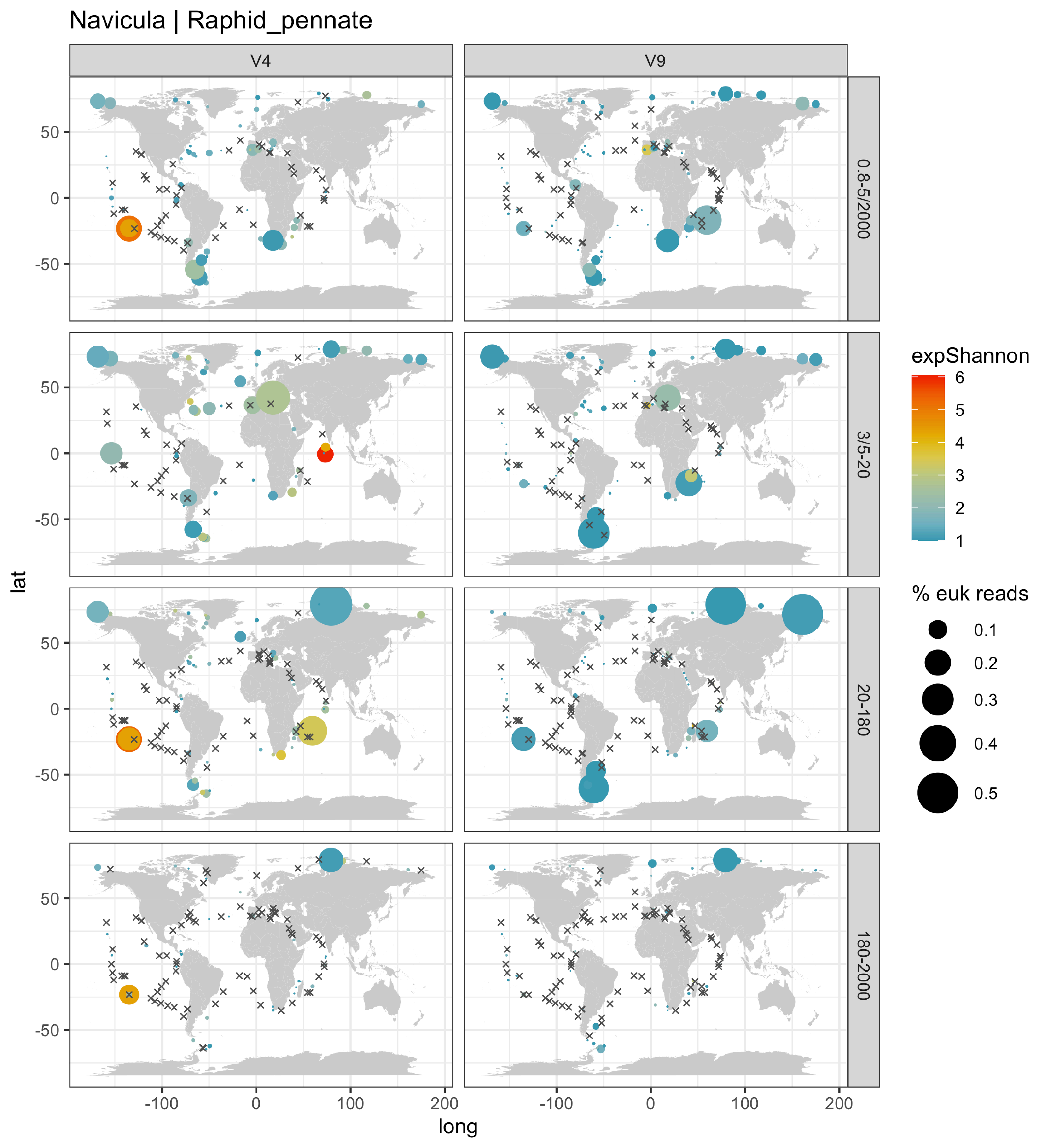

Navicula | Raphid_pennate
V4
V9
0.8-5/2000
3/5-20
20-180
180-2000
expShannon
6
5
4
3
2
1
% euk reads
0.1
0.2
0.3
0.4
0.5
lat
long
50
0
-50
-100
100
200

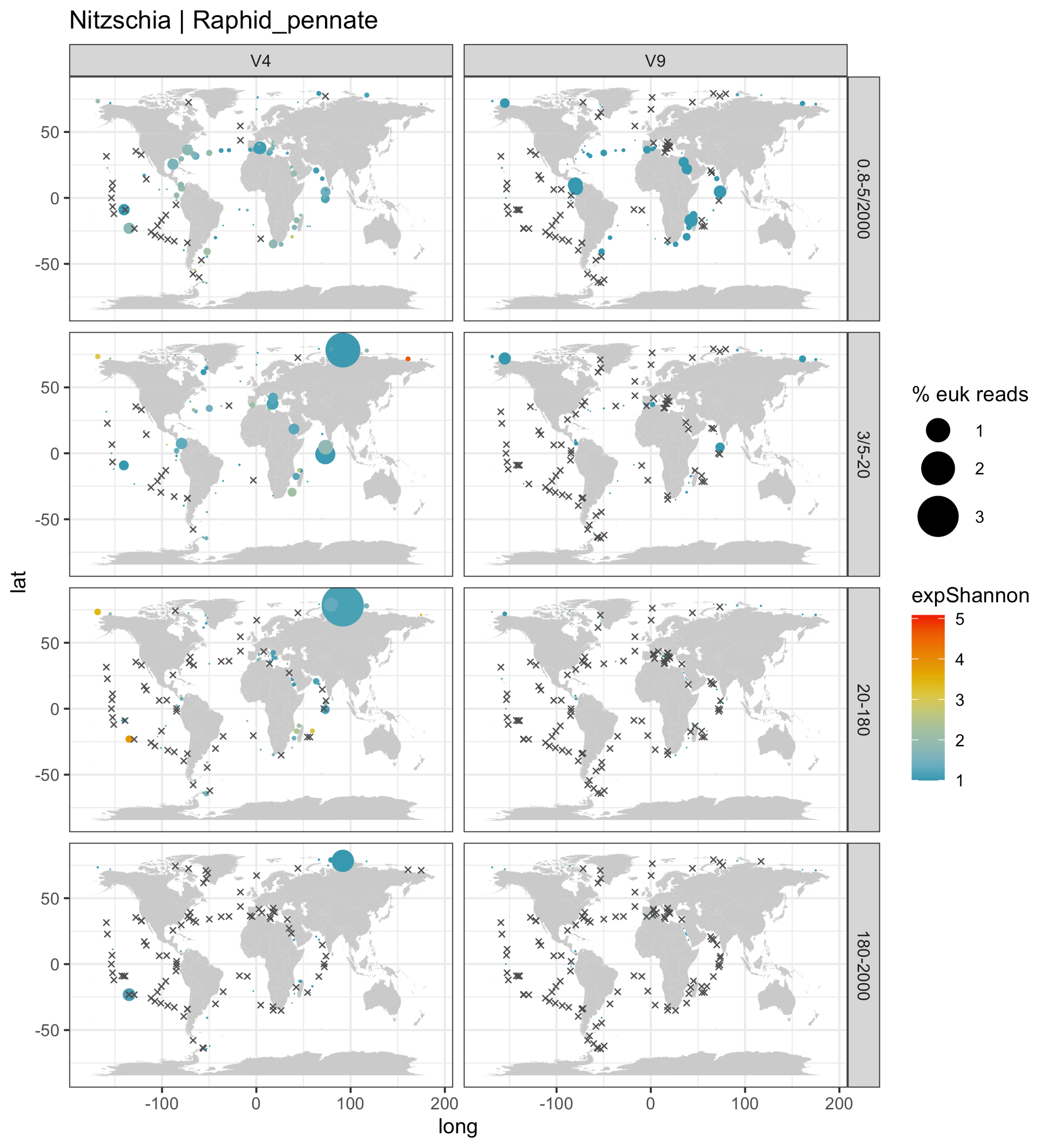

Nitzschia | Raphid_pennate
V4
V9
0.8-5/2000
3/5-20
20-180
180-2000
% euk reads
1
2
3
expShannon
5
4
3
2
1
lat
long
50
0
-50
-100
0
100
200

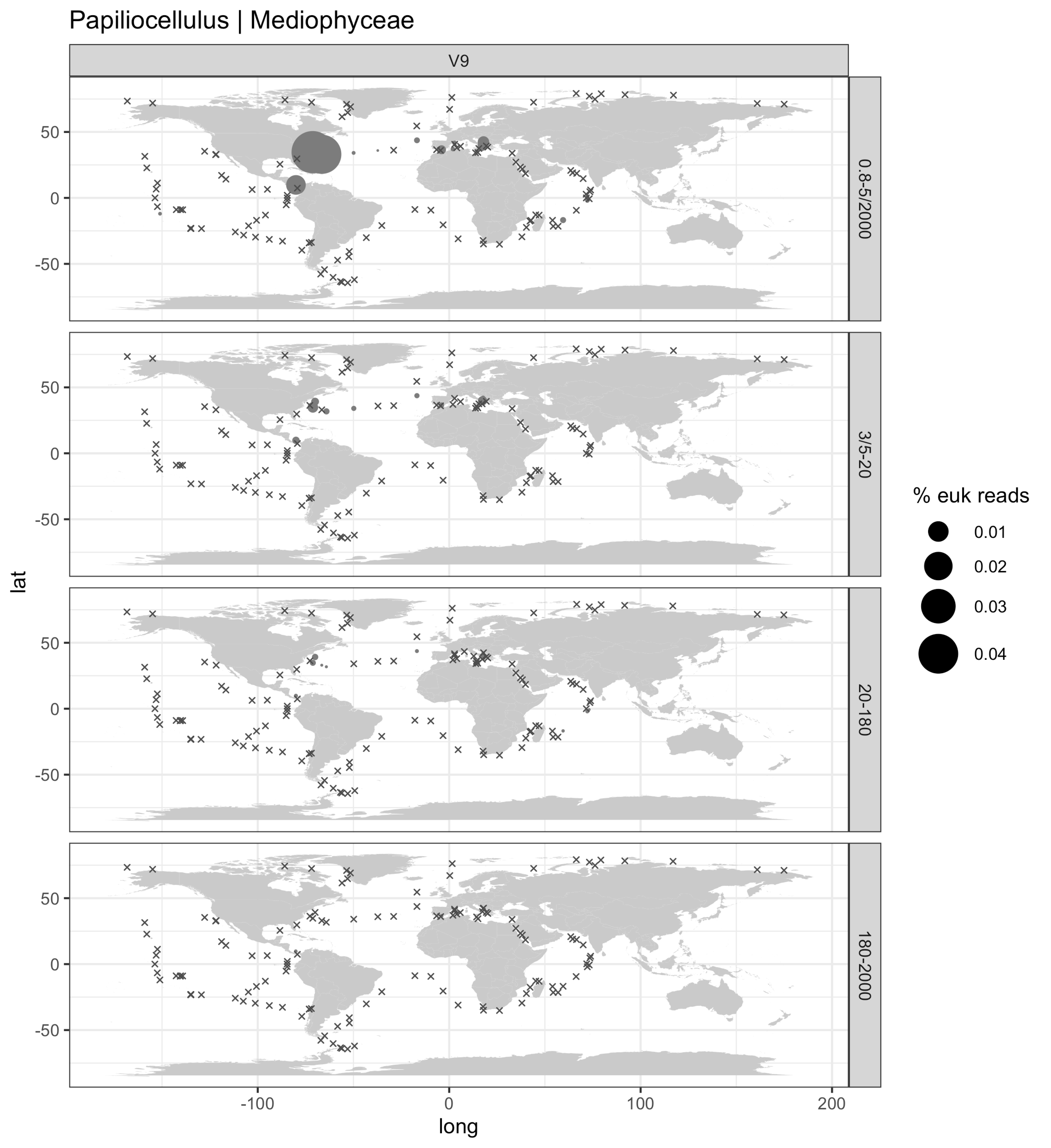
Papiliocellulus | Mediophyceae
V9
0.8-5/2000
3/5-20
20-180
180-2000
lat
long
50
0
-50
-100
100
200
% euk reads
0.01
0.02
0.03
0.04

Paralia | Coscinodiscophyceae

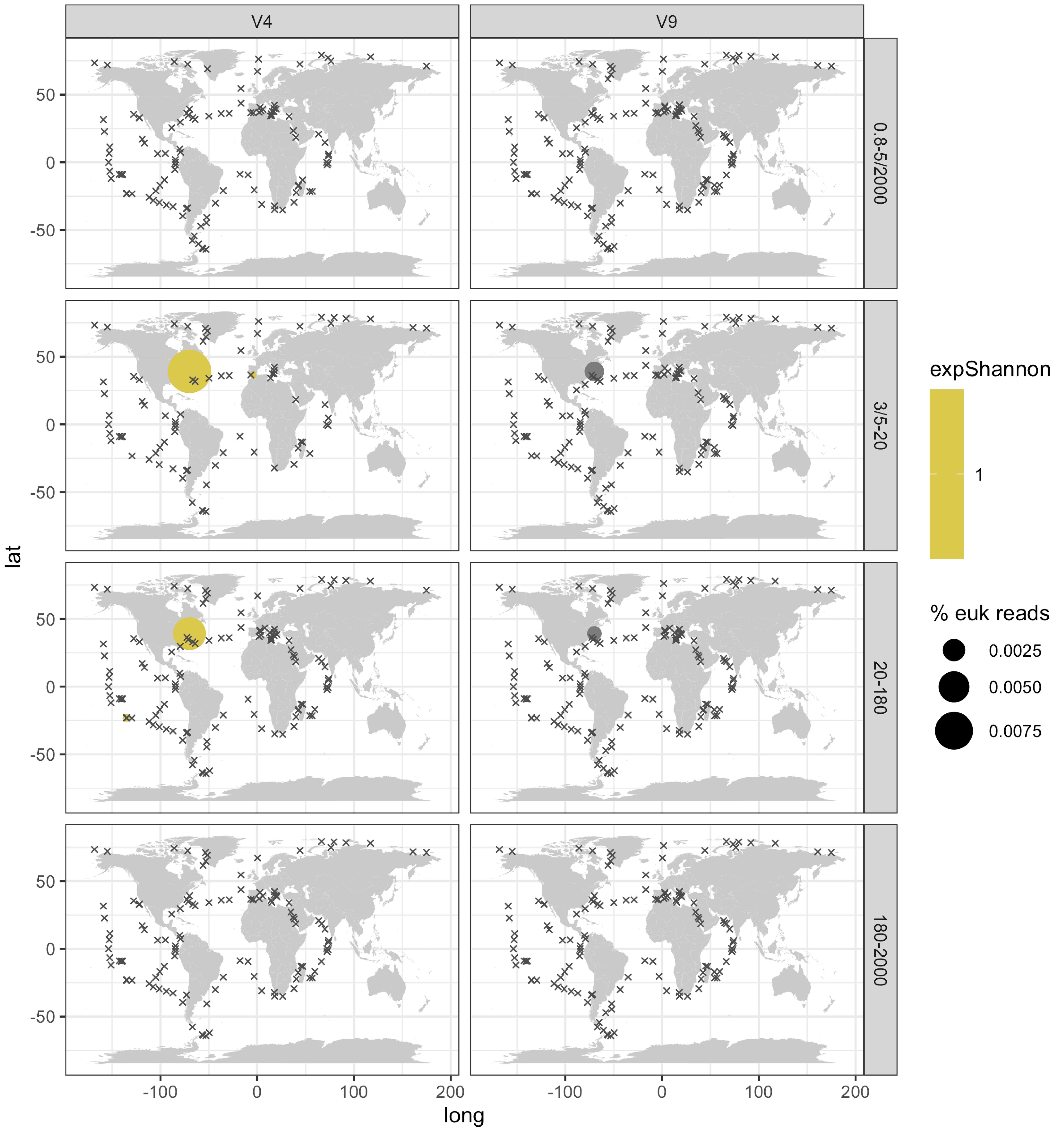

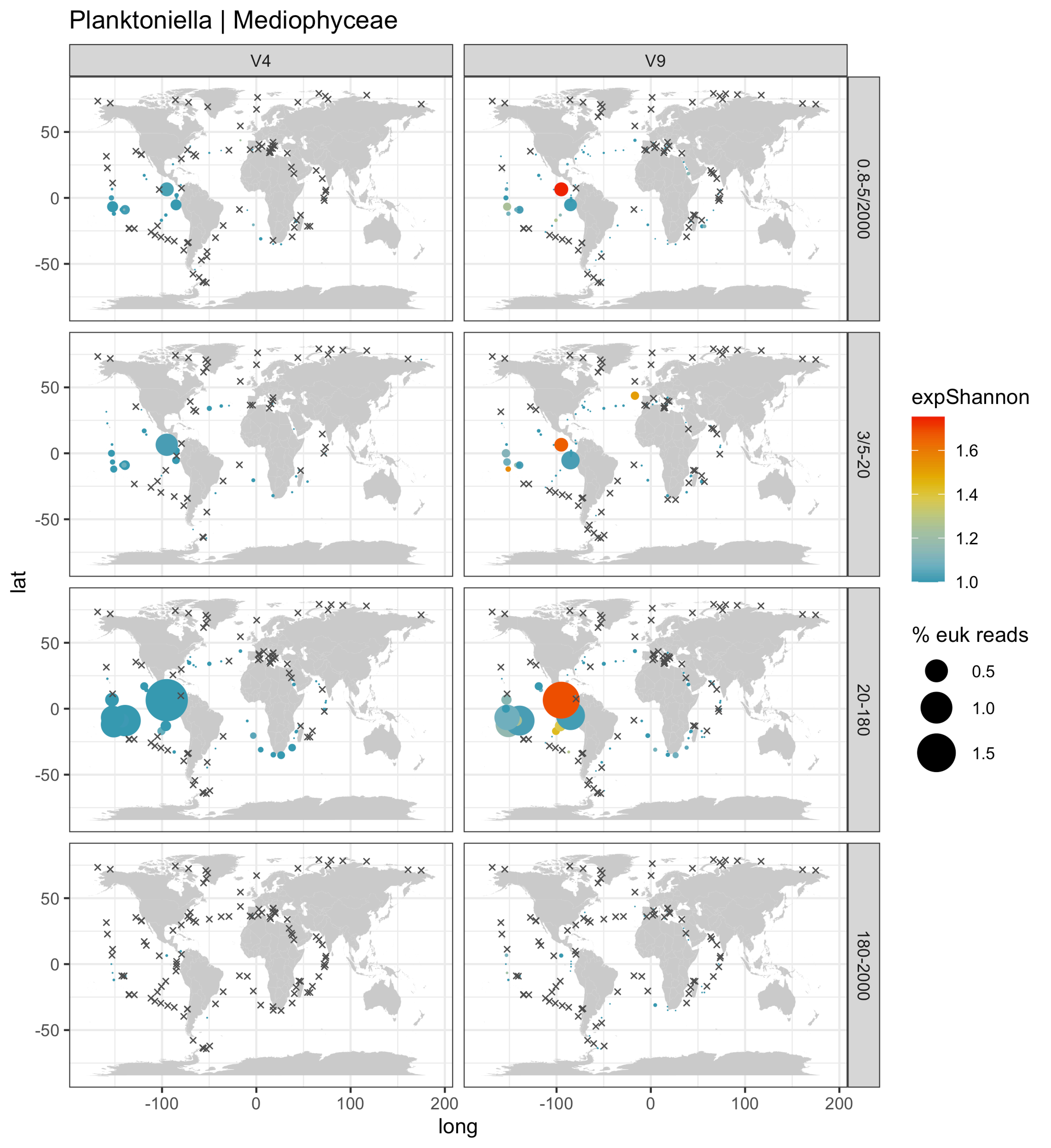
Planktoniella | Mediophyceae
V4
V9
0.8-5/2000
3/5-20
20-180
180-2000
lat
long
50
0
-50
-100
100
200
expShannon
1.6
1.4
1.2
1.0
% euk reads
0.5
1.0
1.5

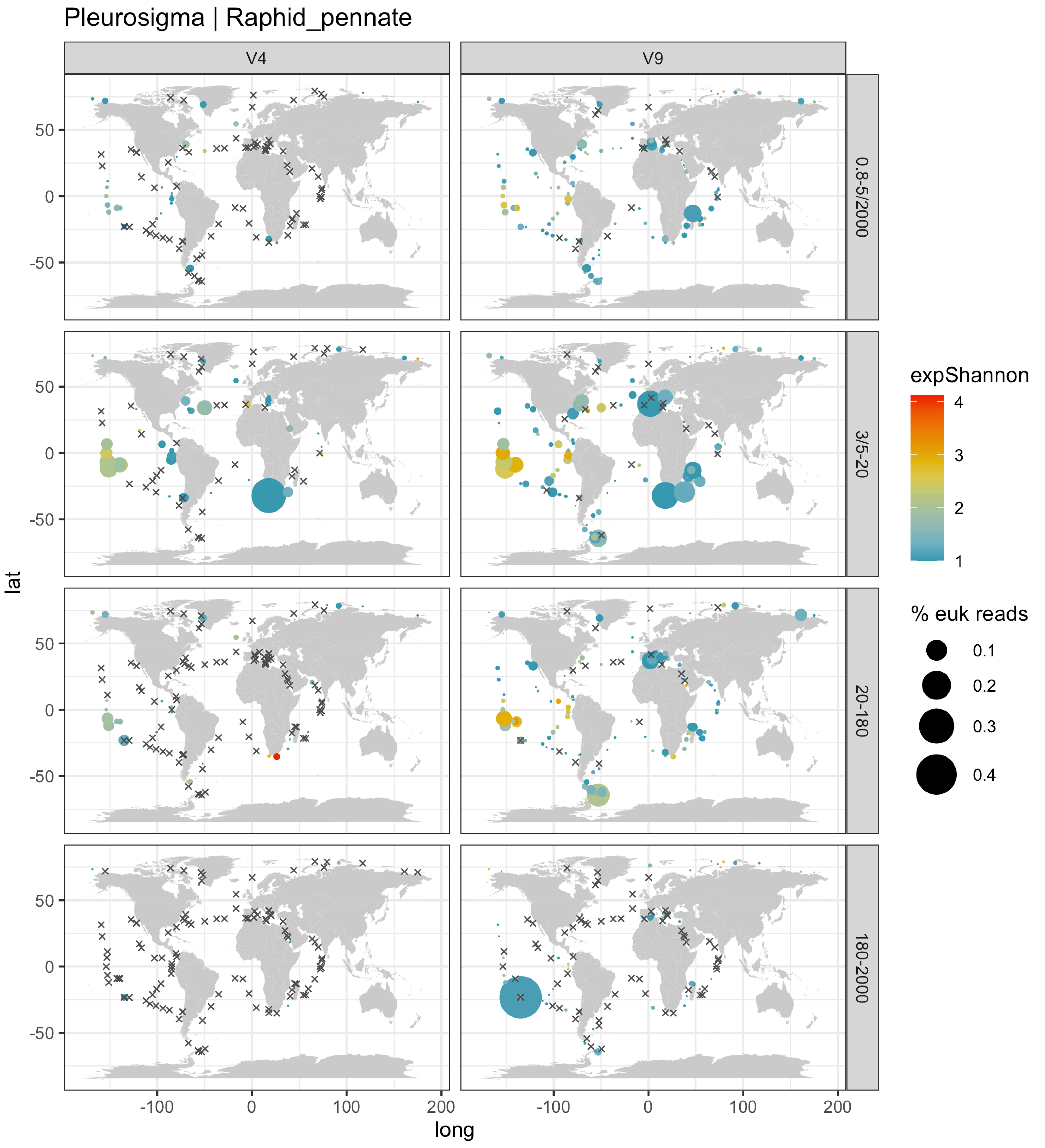

Pleurosigma | Raphid_pennate
V4
V9
0.8-5/2000
3/5-20
20-180
180-2000
expShannon
4
3
2
1
% euk reads
0.1
0.2
0.3
0.4
lat
long
50
0
-50
-100
0
100
200

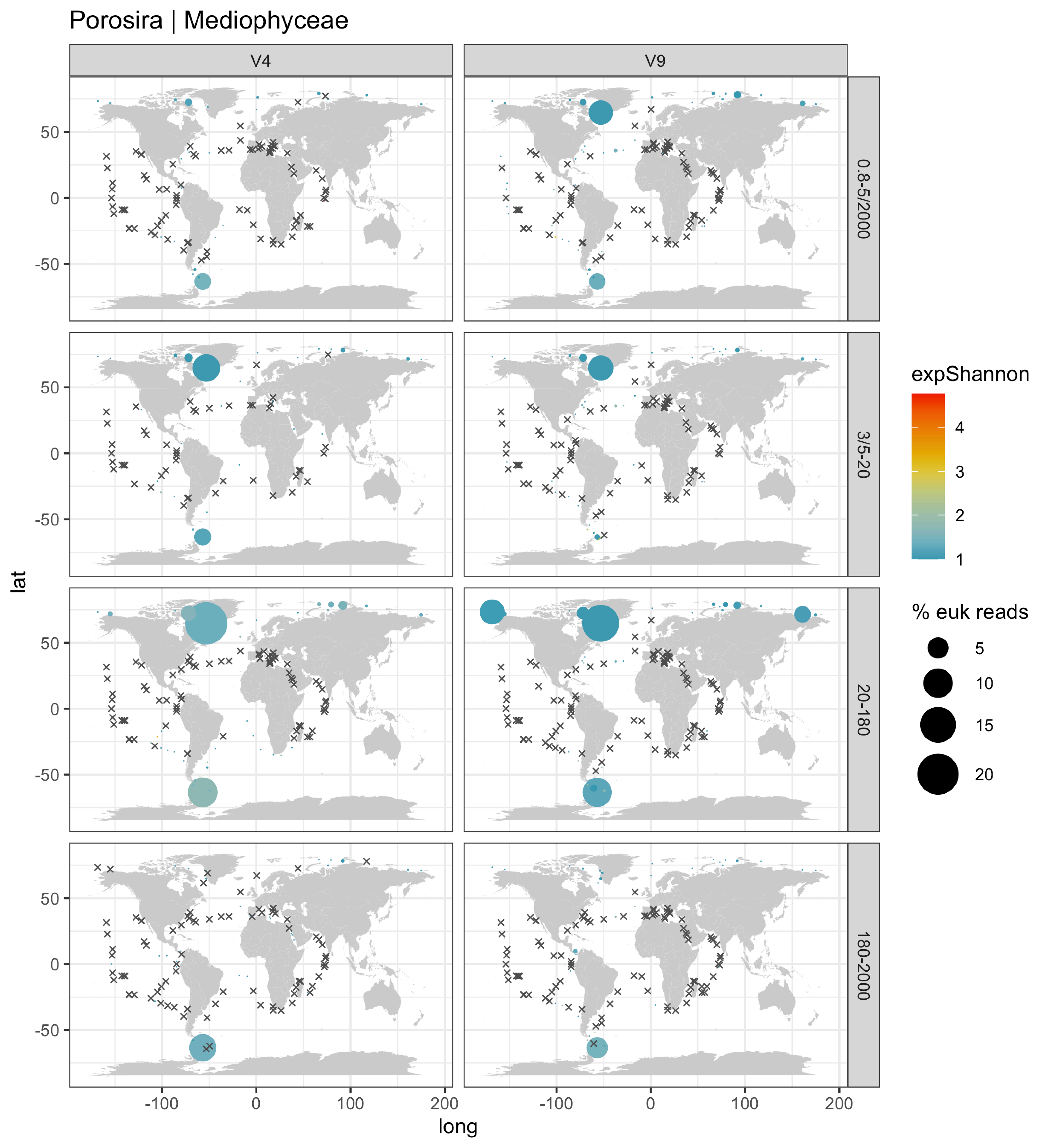

Porosira | Mediophyceae
V4
V9
0.8-5/2000
3/5-20
20-180
180-2000
lat
long
50
0
-50
-100
100
200
expShannon
4
3
2
1
% euk reads
5
10
15
20

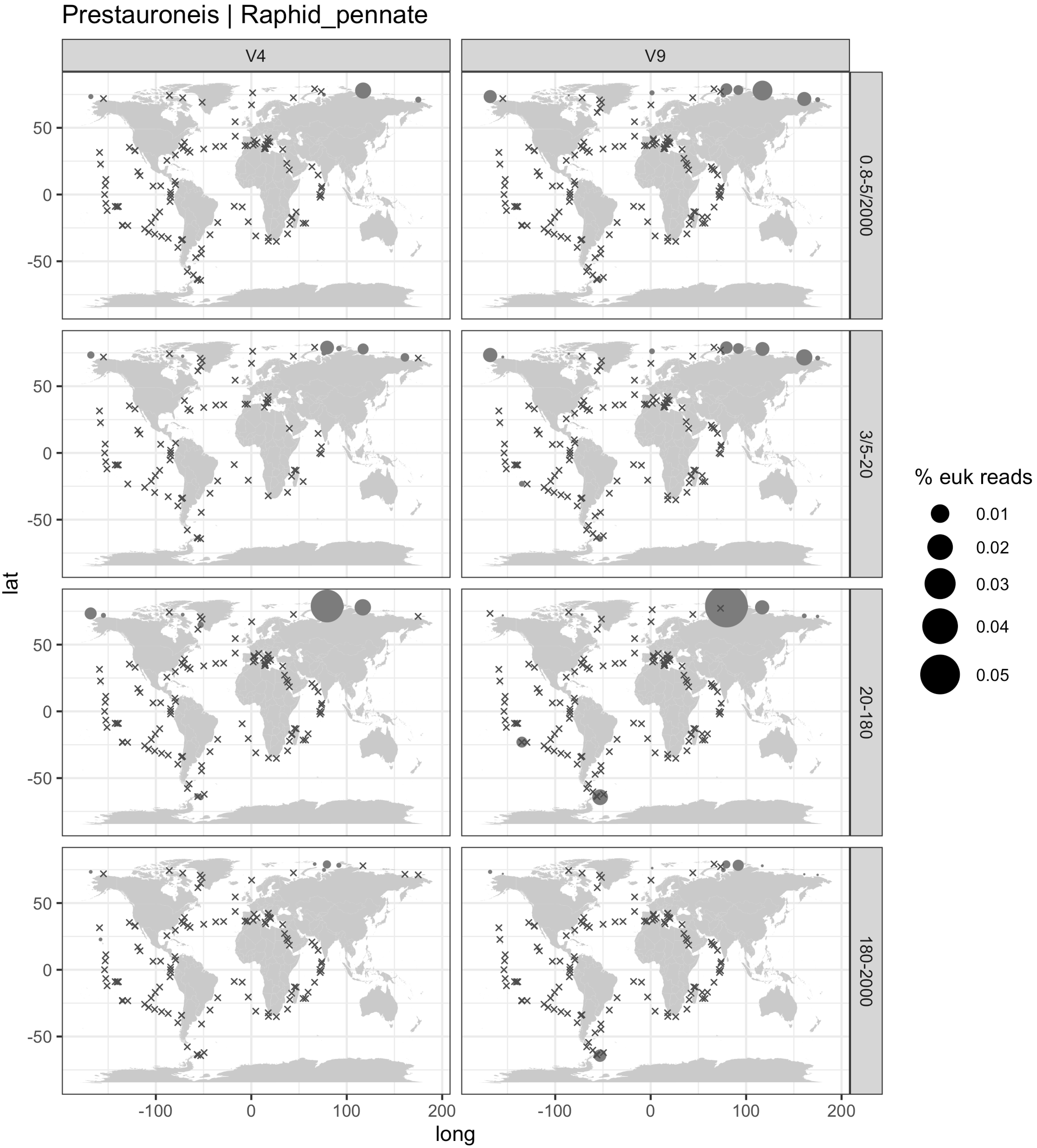

Prestauroneis | Raphid_pennate
V4
V9
0.8-5/2000
3/5-20
20-180
180-2000
lat
long
50
0
-50
-100
100
200
% euk reads
0.01
0.02
0.03
0.04
0.05

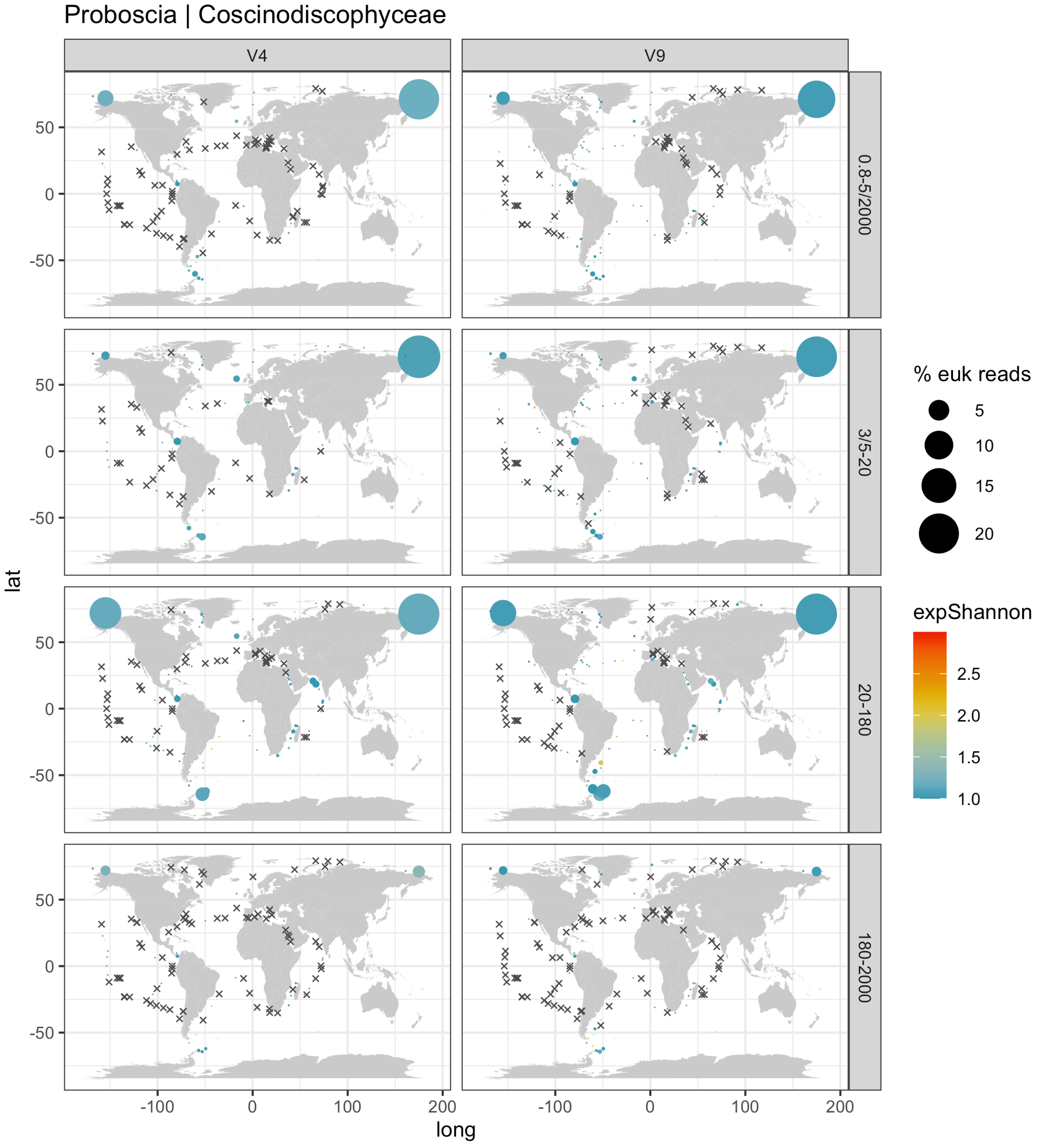

Proboscia | Coscinodiscophyceae
V4
V9
0.8-5/2000
3/5-20
20-180
180-2000
lat
long
50
0
-50
-100
100
200
% euk reads
5
10
15
20
expShannon
2.5
2.0
1.5
1.0

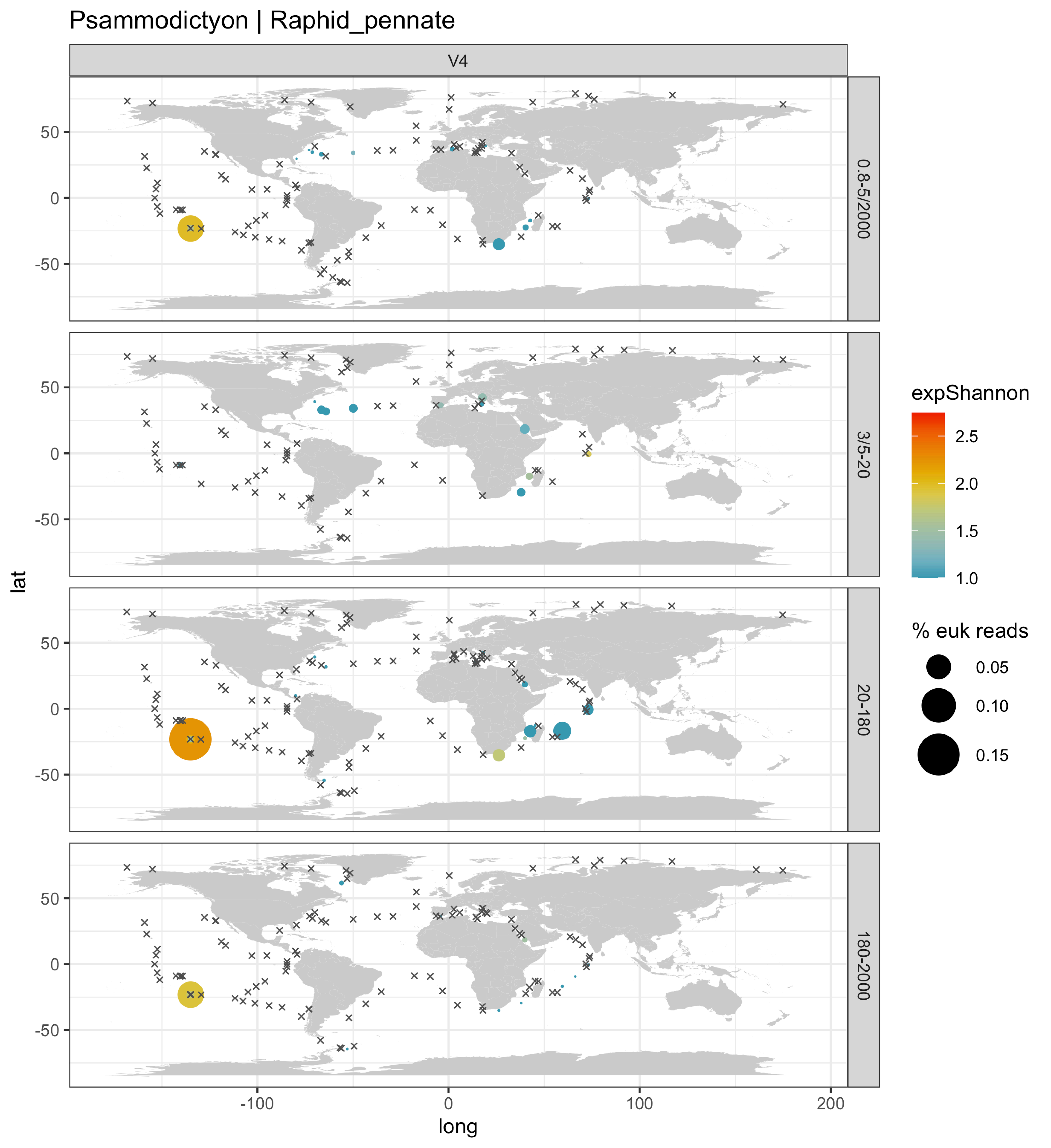
Psammodictyon | Raphid_pennate
V4
0.8-5/2000
3/5-20
20-180
180-2000
lat
long
50
0
-50
-100
100
200
expShannon
2.5
2.0
1.5
1.0
% euk reads
0.05
0.10
0.15

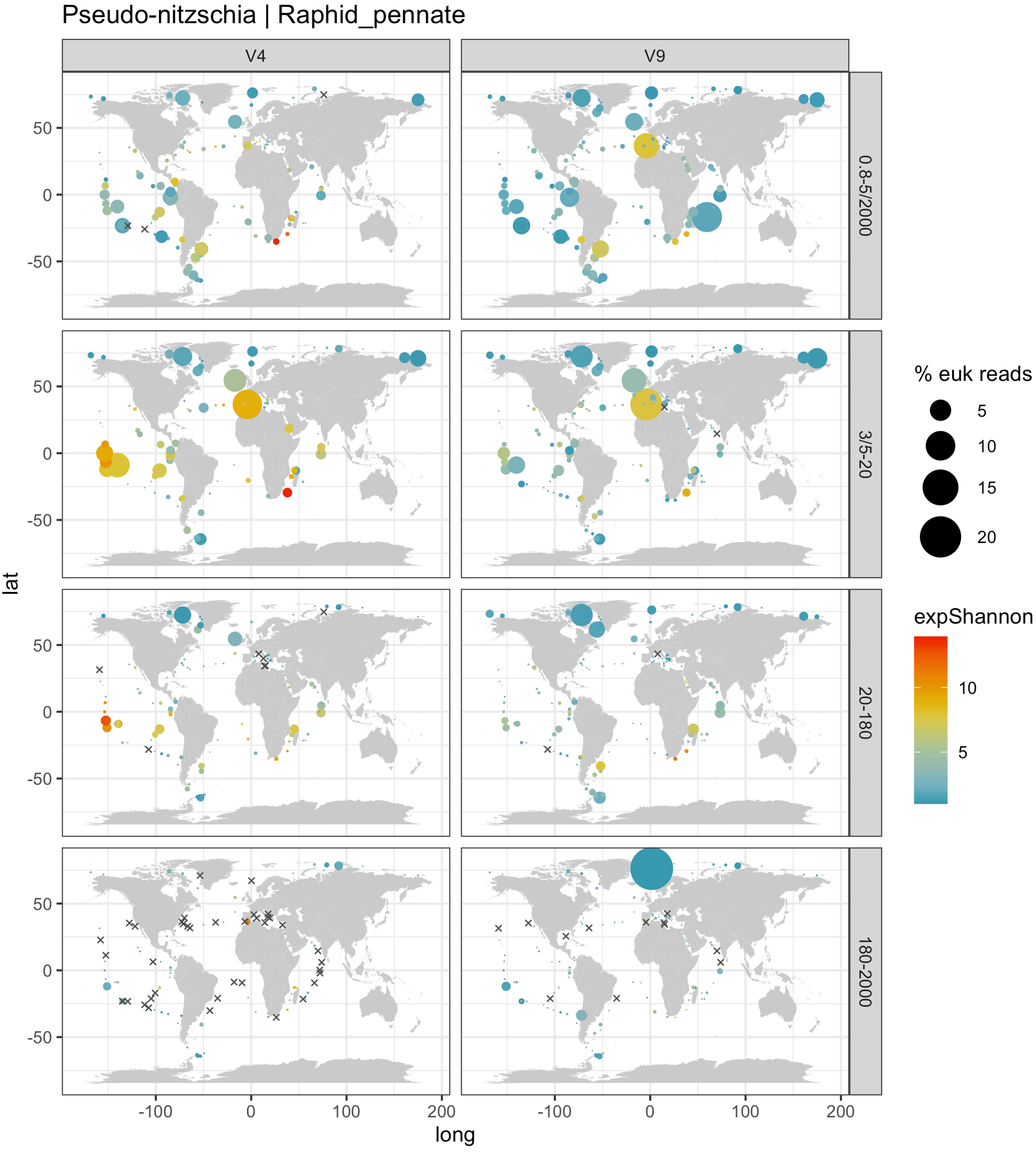

Pseudo-nitzschia | Raphid_pennate
V4
V9
0.8-5/2000
3/5-20
20-180
180-2000
% euk reads
5
10
15
20
expShannon
10
5
lat
50
0
-50
long
-100
0
100
200

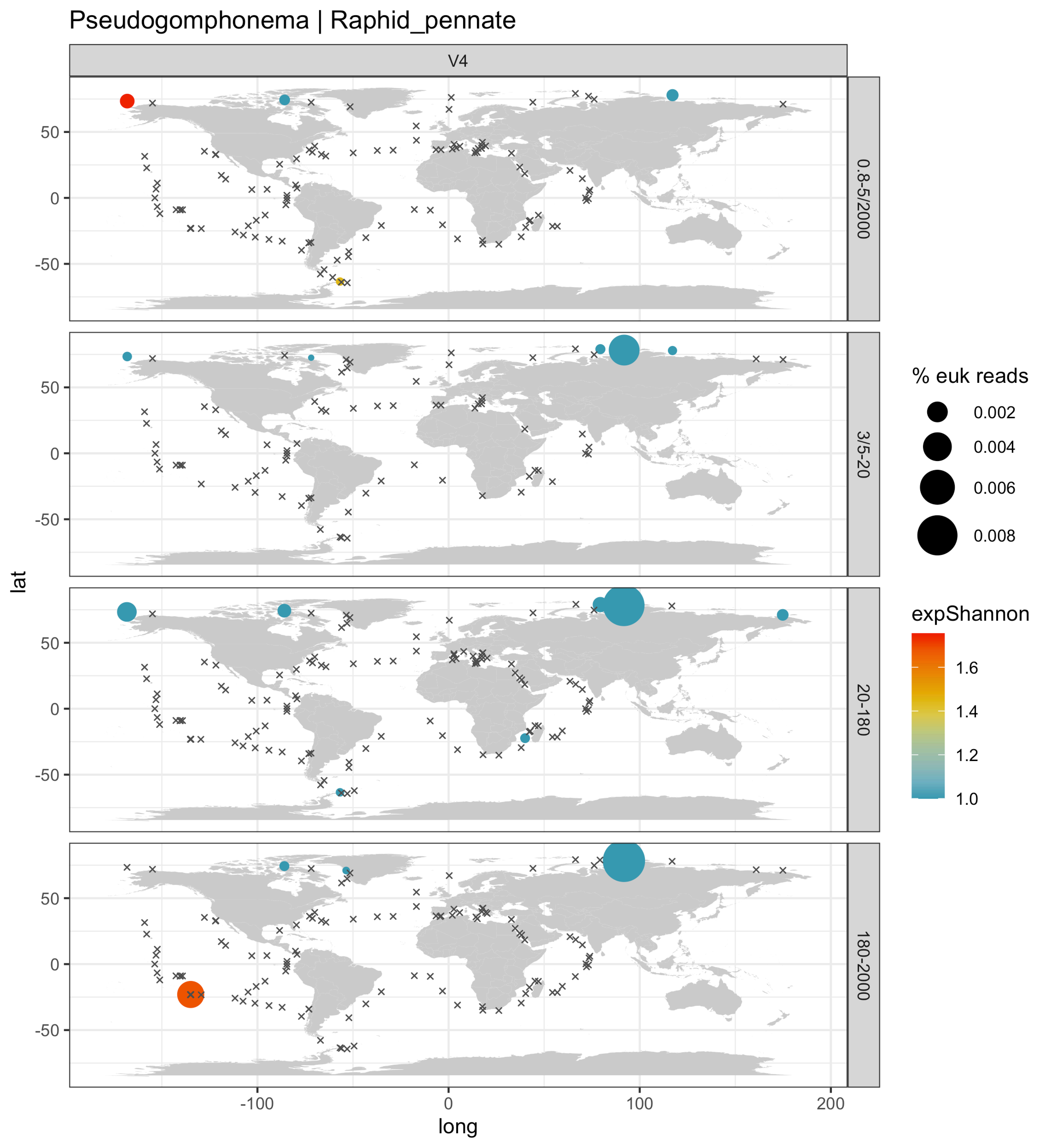

Pseudogomphonema | Raphid_pennate
V4
0.8-5/2000
3/5-20
20-180
180-2000
lat
long
50
0
-50
-100
100
200
% euk reads
0.002
0.004
0.006
0.008
expShannon
1.6
1.4
1.2
1.0

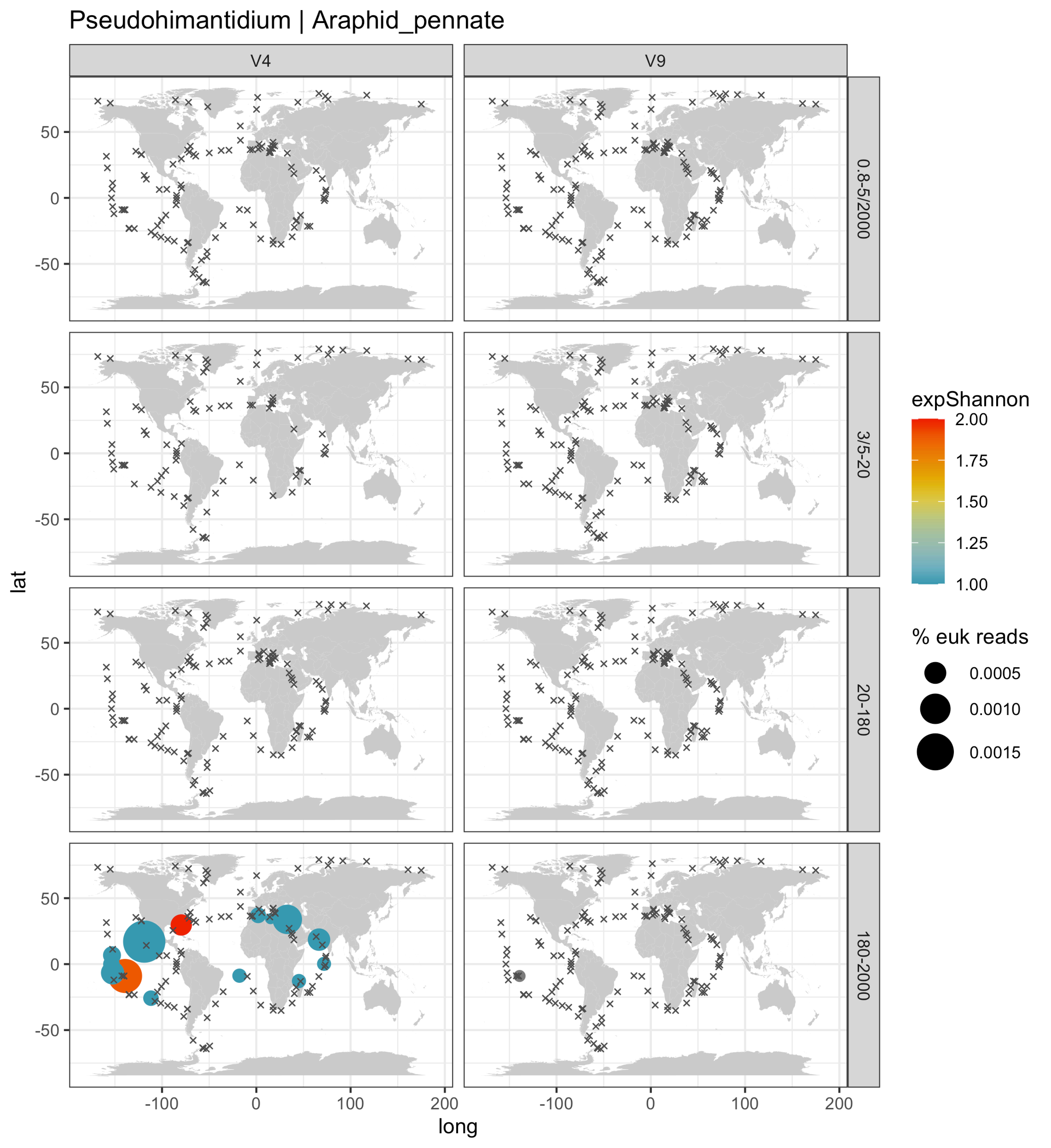

Pseudohimantidium | Araphid_pennate
V4
V9
0.8-5/2000
3/5-20
20-180
180-2000
lat
long
50
0
-50
-100
0
100
200
expShannon
2.00
1.75
1.50
1.25
1.00
% euk reads
0.0005
0.0010
0.0015

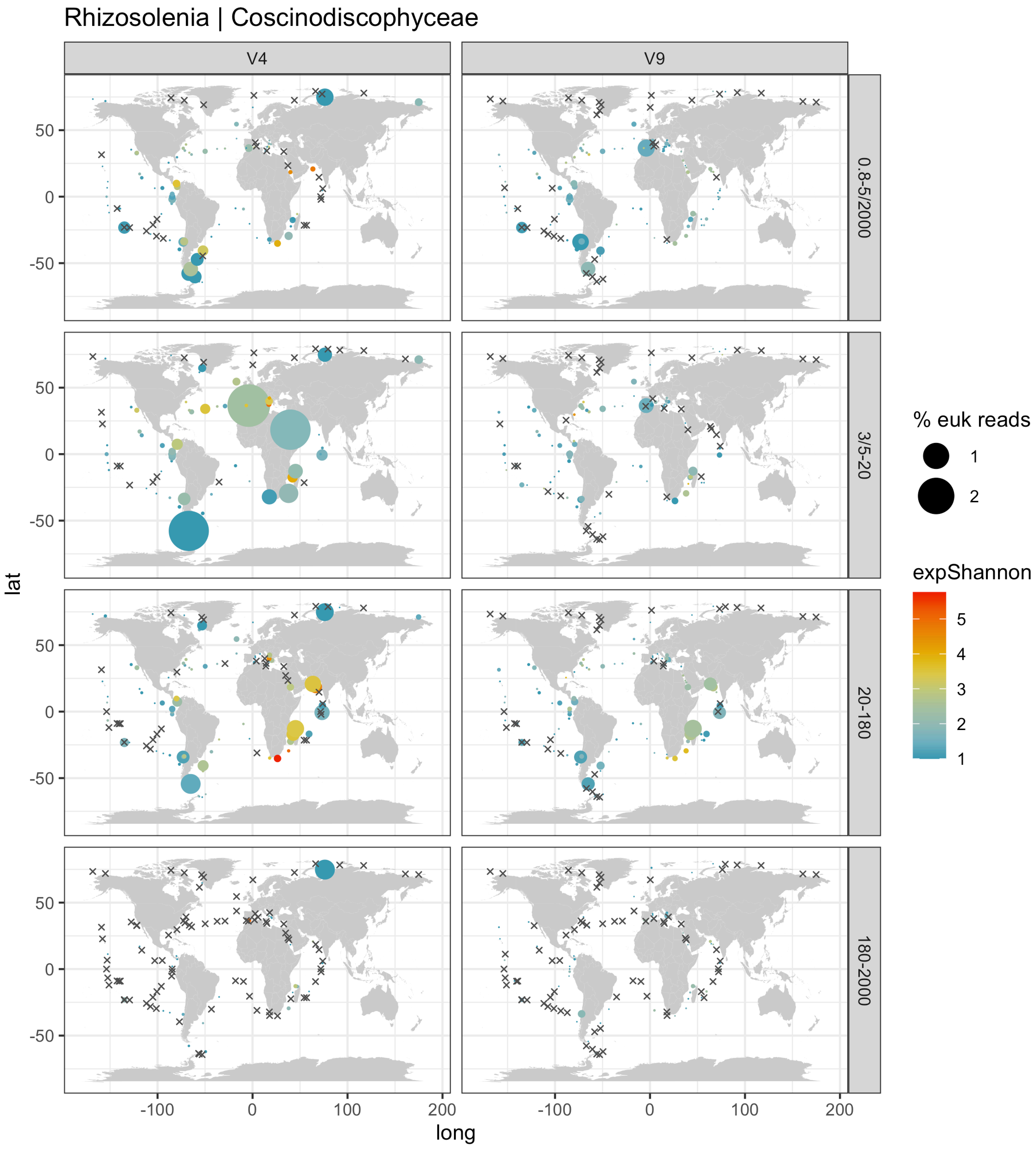

Rhizosolenia | Coscinodiscophyceae
V4
V9
0.8-5/2000
3/5-20
20-180
180-2000
lat
long
50
0
-50
-100
0
100
200
% euk reads
1
2
expShannon
5
4
3
2
1

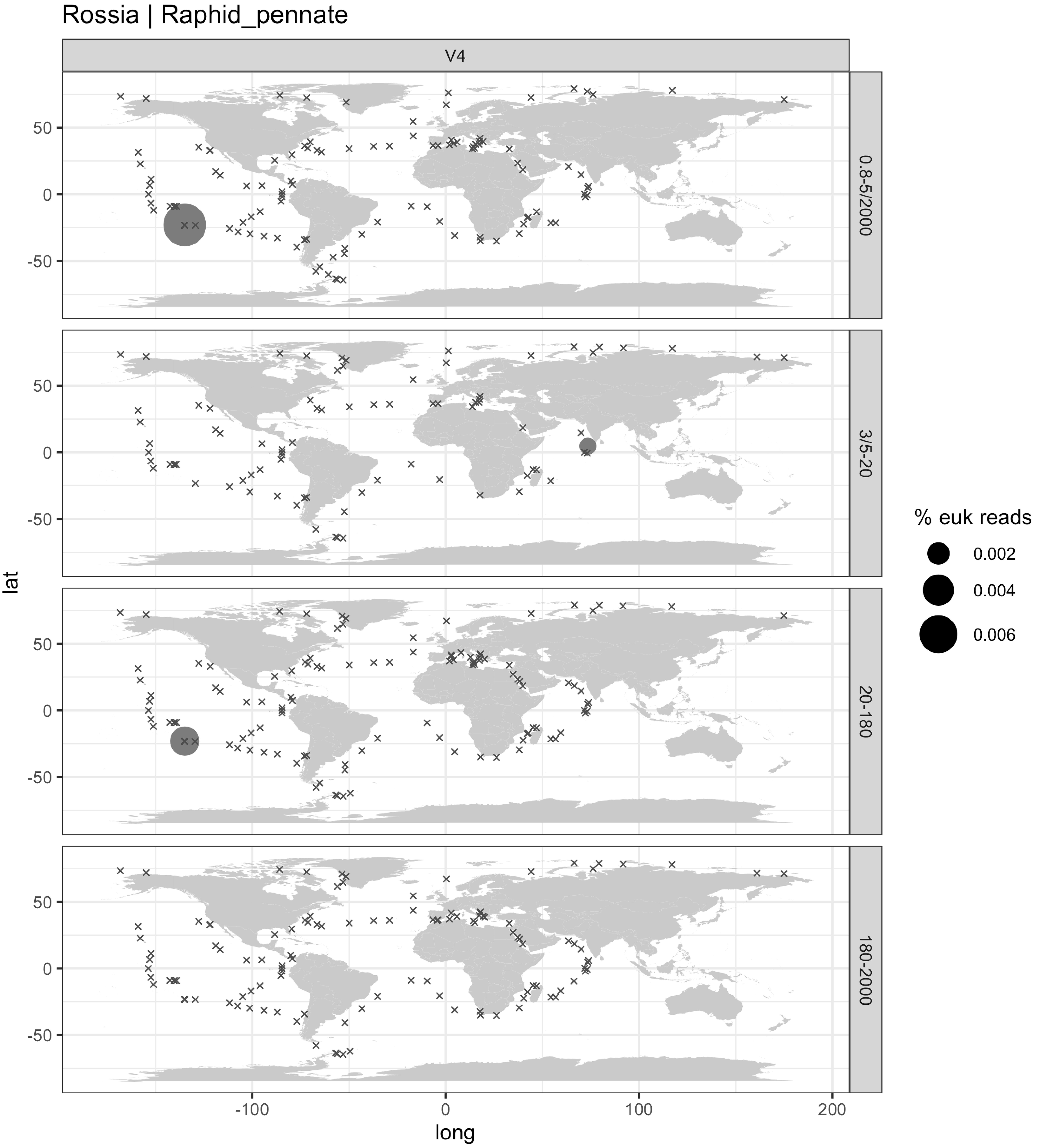

Rossia | Raphid_pennate
V4
0.8-5/2000
3/5-20
20-180
180-2000
lat
long
50
0
-50
-100
0
100
200
% euk reads
0.002
0.004
0.006

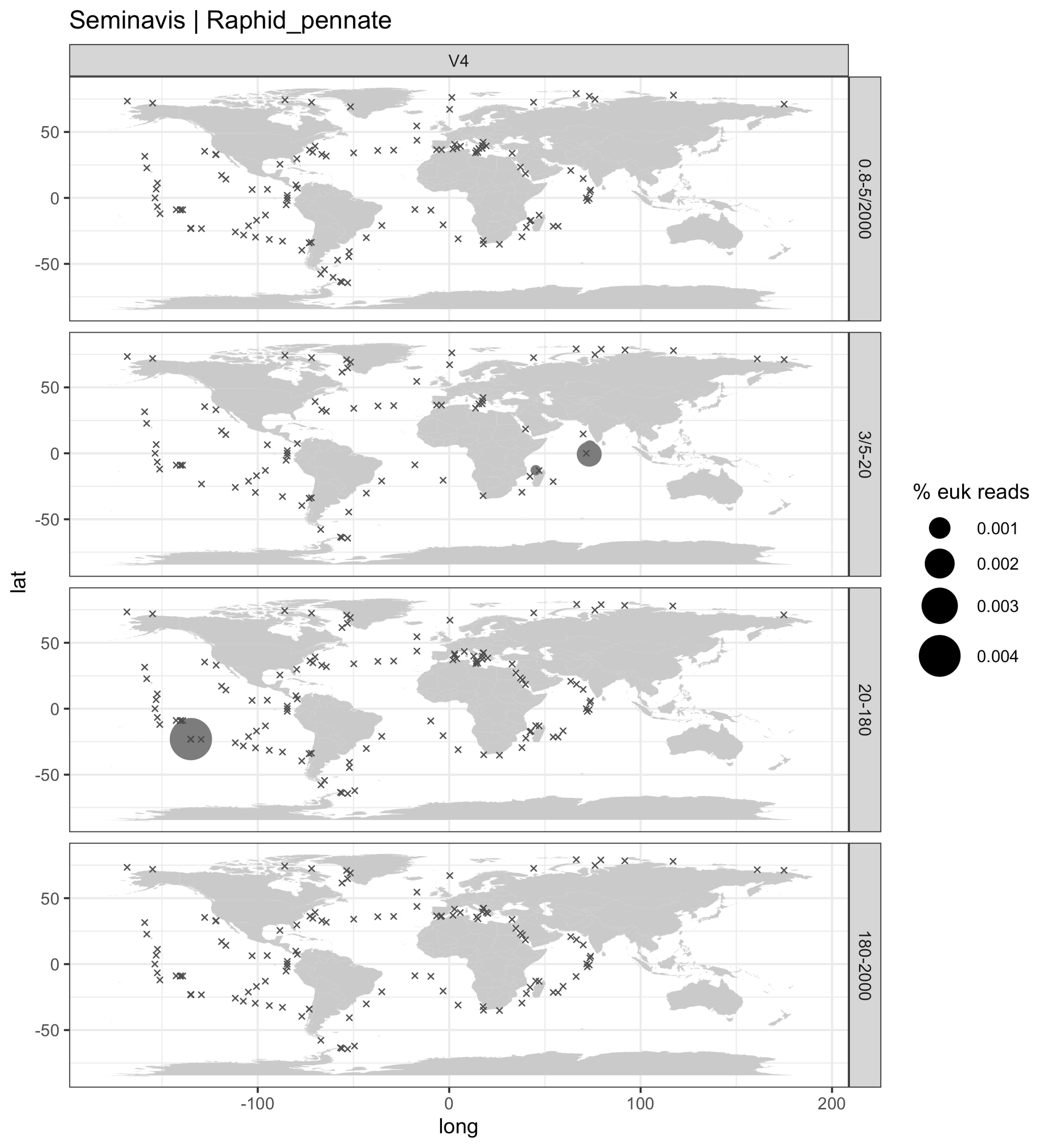
Seminavis | Raphid_pennate
V4
0.8-5/2000
3/5-20
20-180
180-2000
lat
long
50
0
-50
-100
0
100
200
% euk reads
0.001
0.002
0.003
0.004

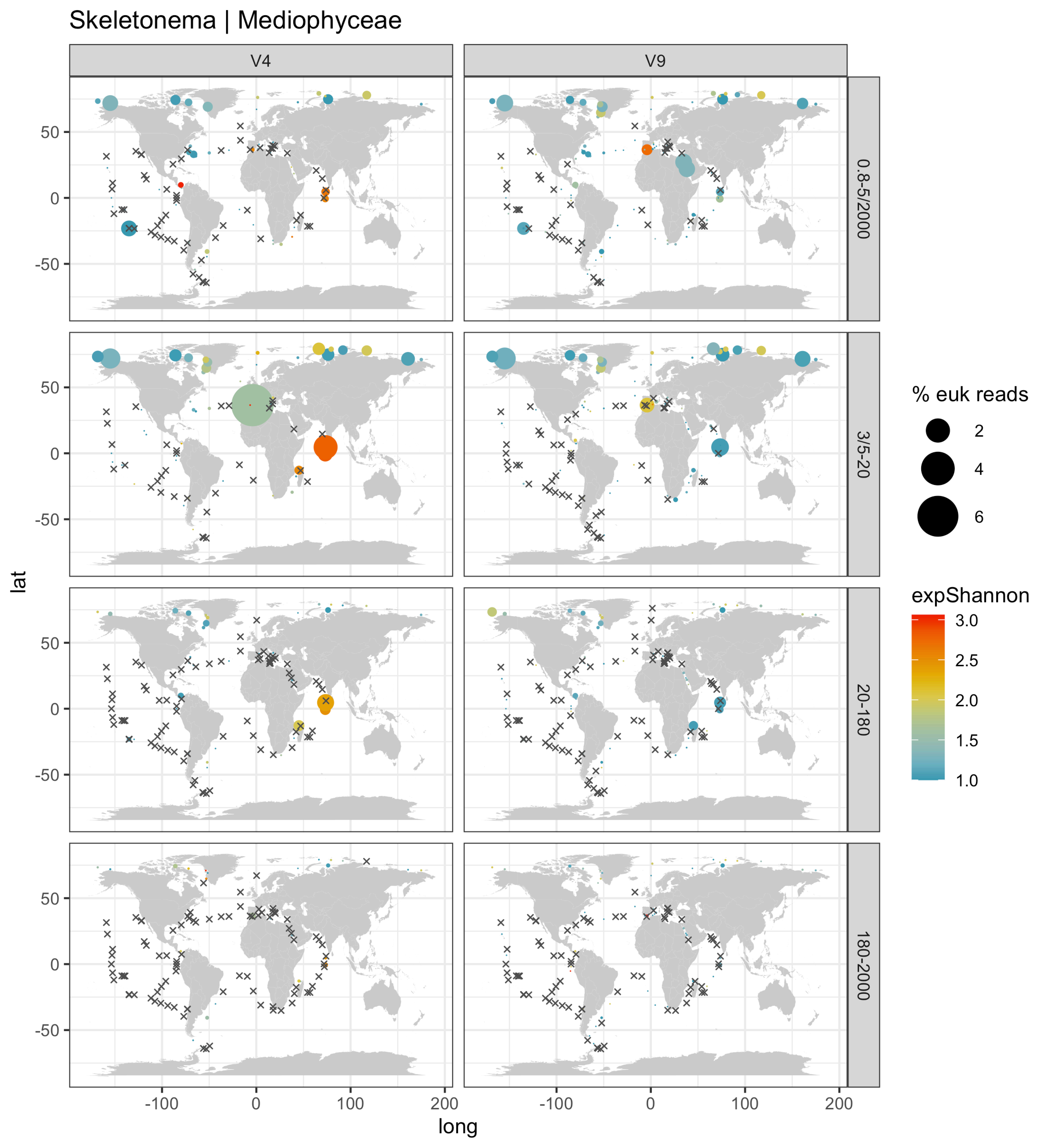

Skeletonema | Mediophyceae
V4
V9
0.8-5/2000
3/5-20
20-180
180-2000
% euk reads
2
4
6
expShannon
3.0
2.5
2.0
1.5
1.0
lat
long
50
0
-50
-100
0
100
200

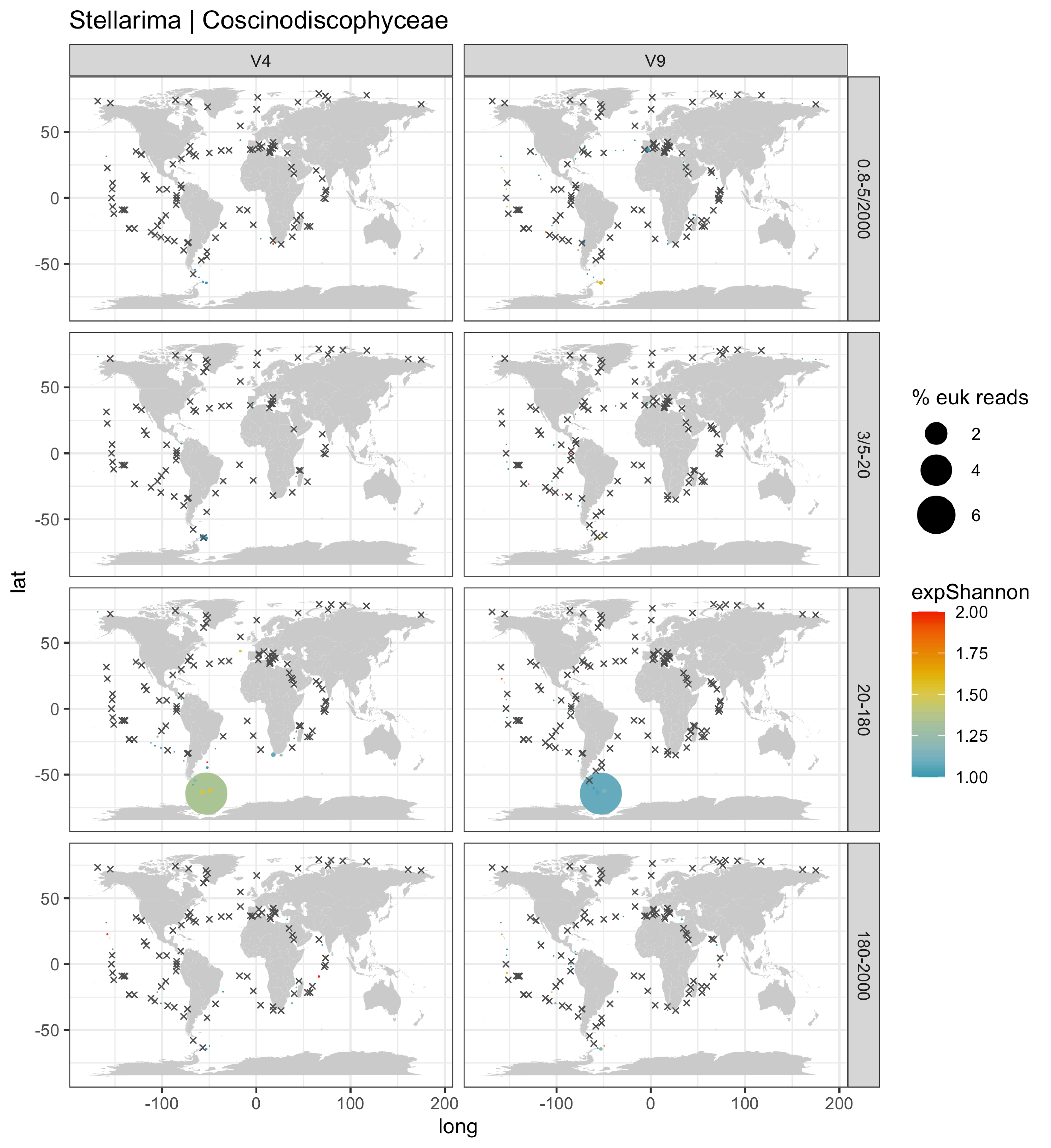
Stellarima | Coscinodiscophyceae
V4
V9
0.8-5/2000
3/5-20
20-180
180-2000
lat
long
% euk reads
2
4
6
expShannon
2.00
1.75
1.50
1.25
1.00
50
0
-50
-100
0
100
200

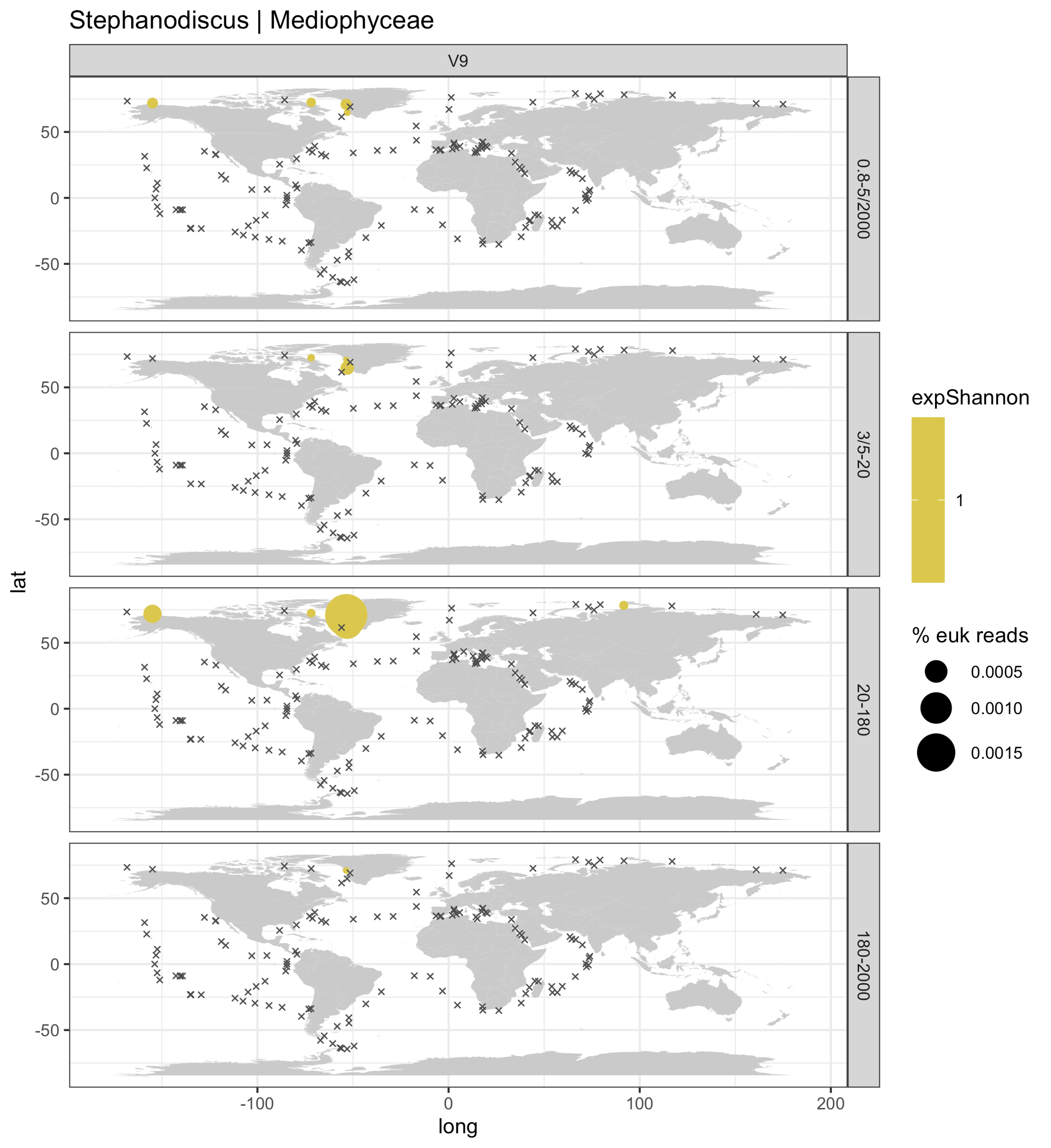
Stephanodiscus | Mediophyceae
V9
0.8-5/2000
3/5-20
20-180
180-2000
lat
long
50
0
-50
-100
100
200
expShannon
1
% euk reads
0.0005
0.0010
0.0015

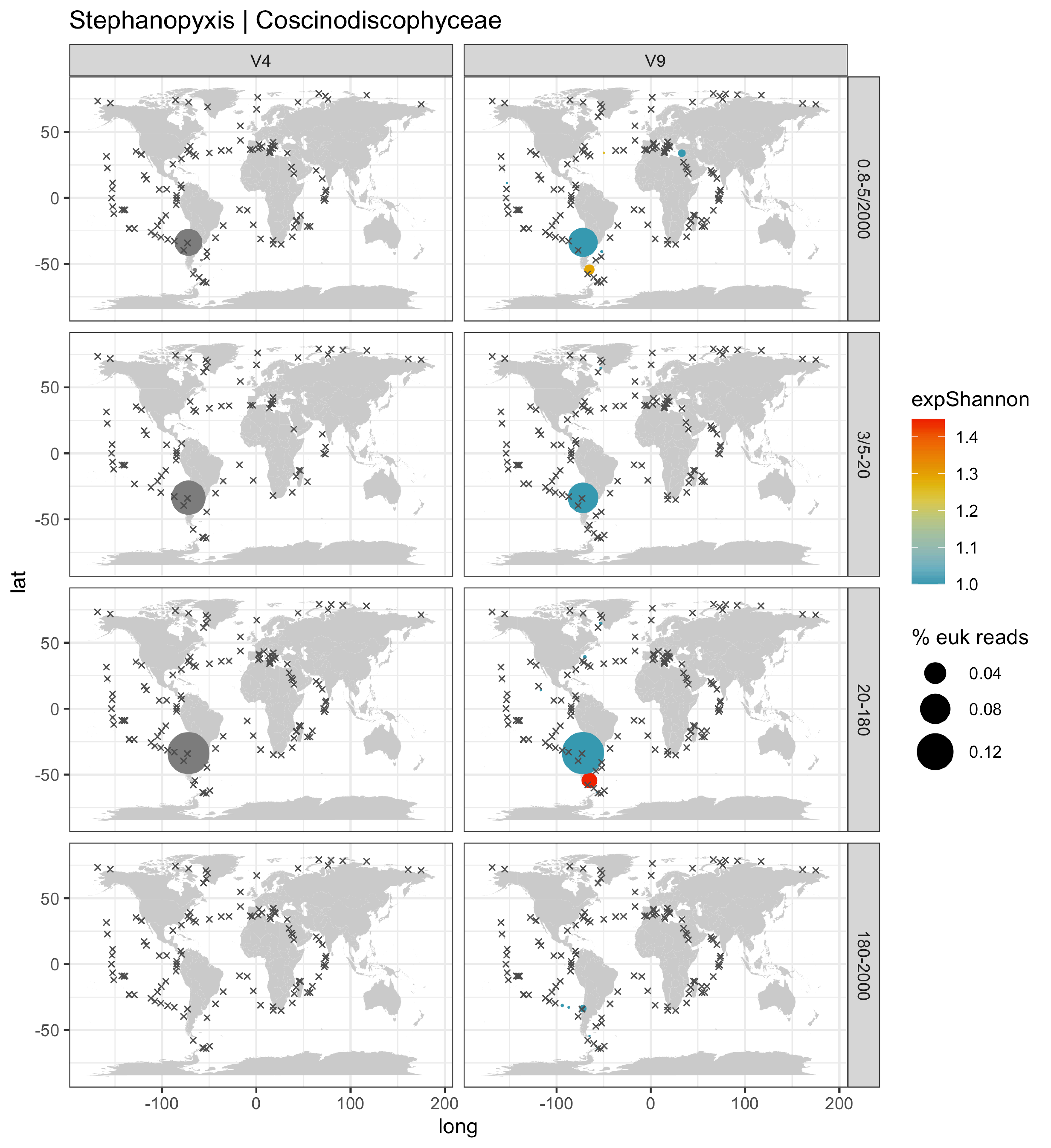

Stephanopyxis | Coscinodiscophyceae
V4
V9
0.8-5/2000
3/5-20
20-180
180-2000
lat
long
50
0
-50
-100
0
100
200
expShannon
1.4
1.3
1.2
1.1
1.0
% euk reads
0.04
0.08
0.12

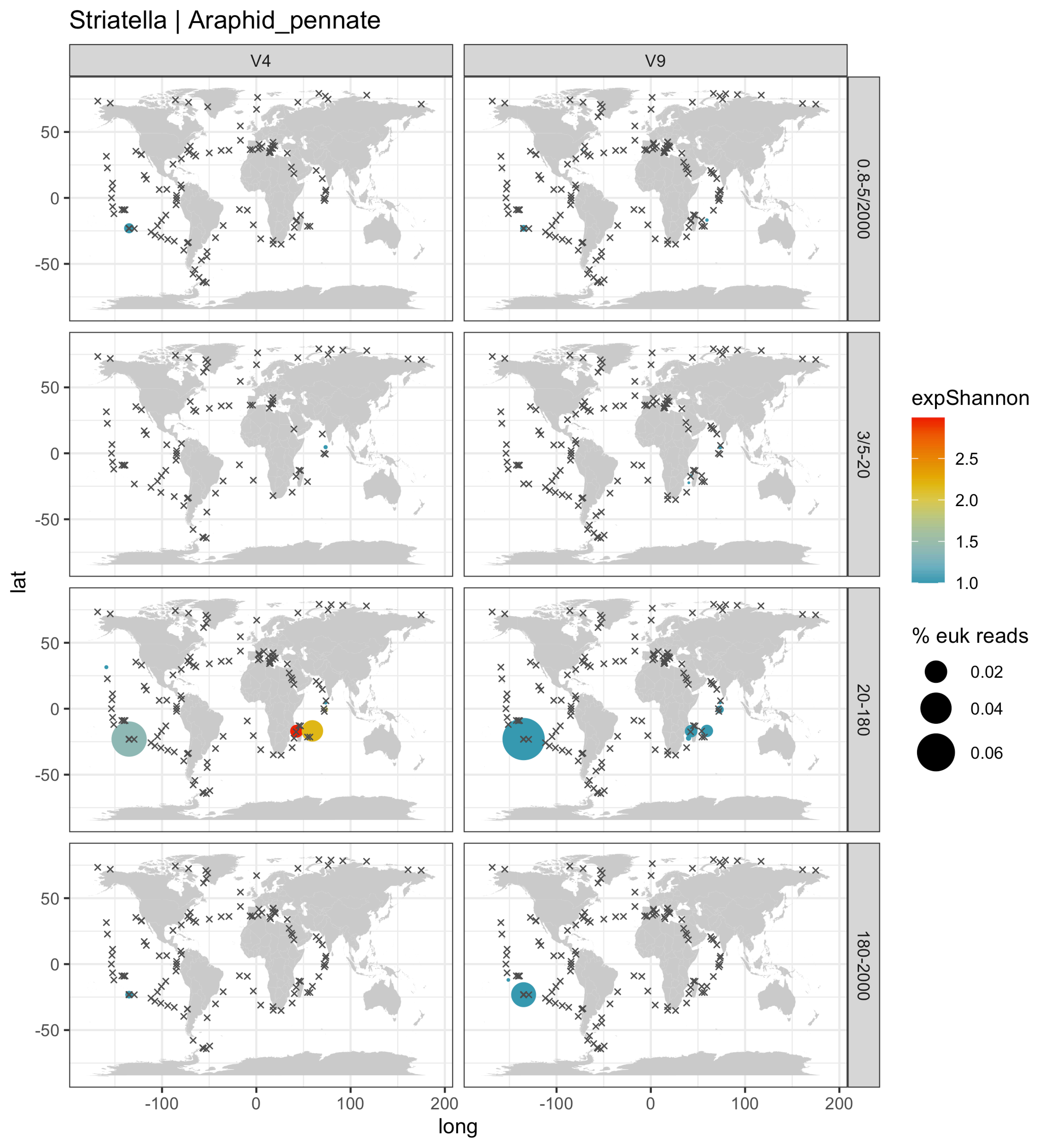

Striatella | Araphid_pennate
V4
V9
0.8-5/2000
3/5-20
20-180
180-2000
lat
long
-100
0
100
200
50
0
-50
expShannon
2.5
2.0
1.5
1.0
% euk reads
0.02
0.04
0.06

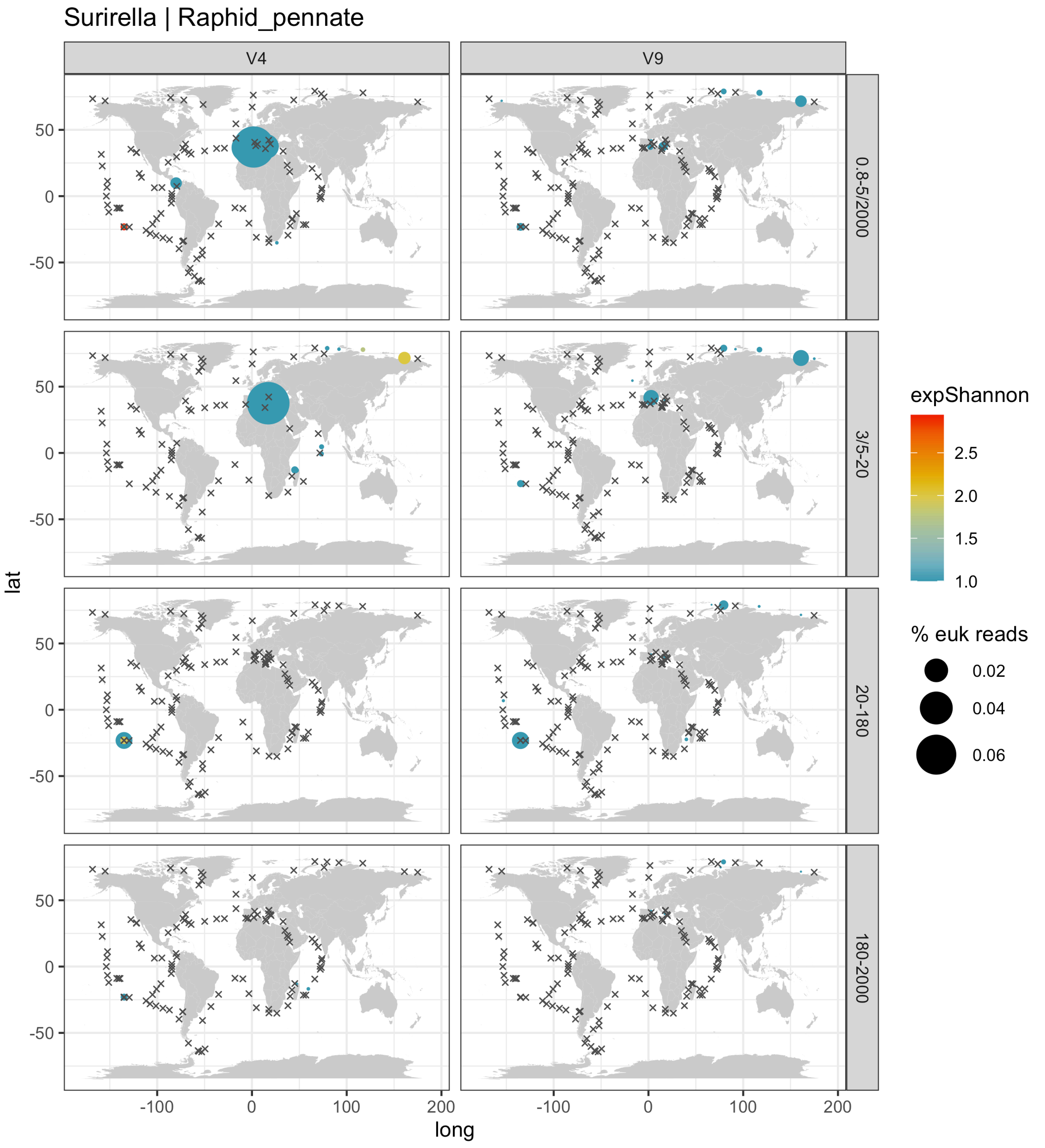

Surirella | Raphid_pennate
V4
V9
0.8-5/2000
3/5-20
20-180
180-2000
lat
long
50
0
-50
-100
0
100
200
expShannon
2.5
2.0
1.5
1.0
% euk reads
0.02
0.04
0.06

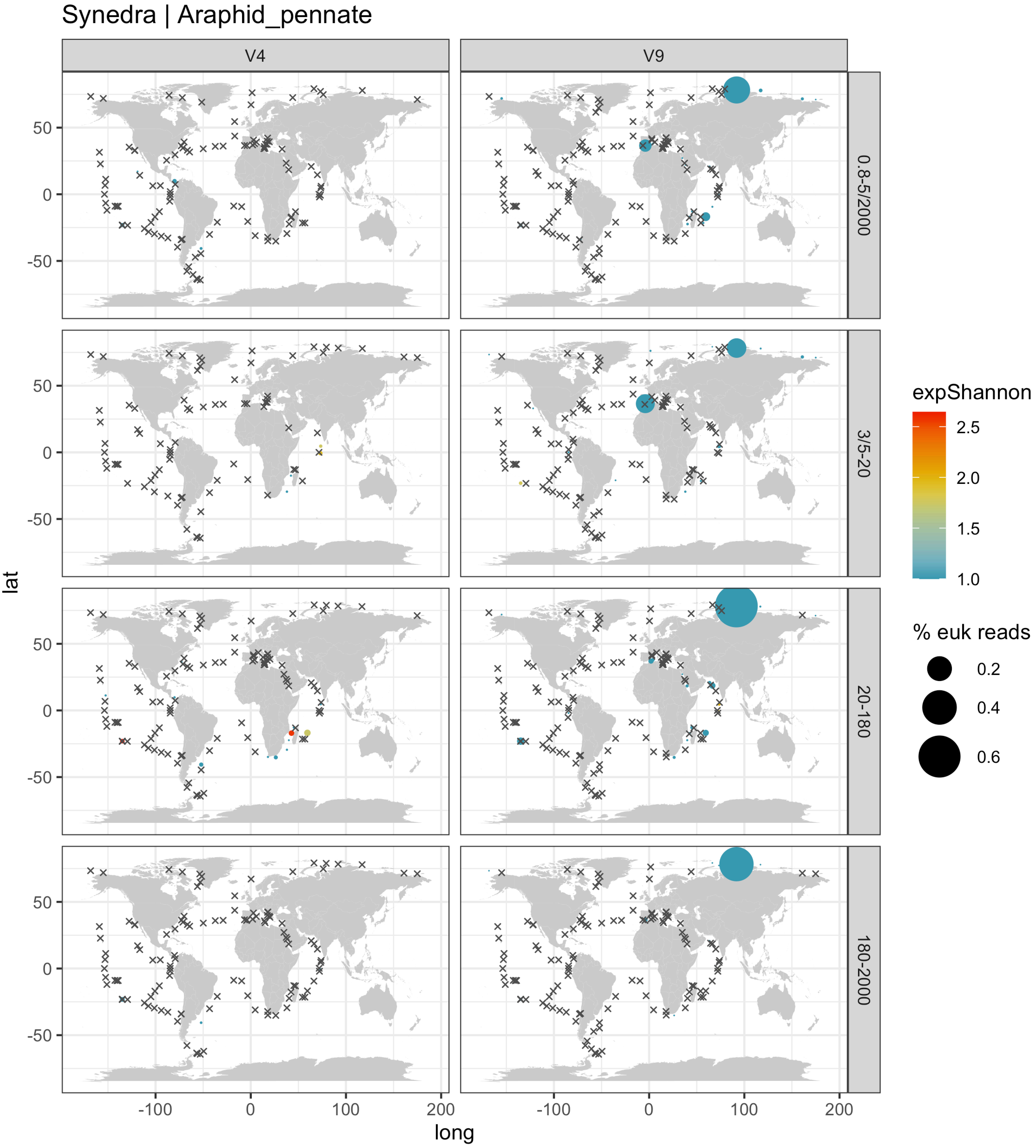

Synedra | Araphid_pennate
V4
V9
0.8-5/2000
3/5-20
20-180
180-2000
lat
long
expShannon
2.5
2.0
1.5
1.0
% euk reads
0.2
0.4
0.6
50
0
-50
-100
0
100
200

### Talaroneis | Araphid_pennate

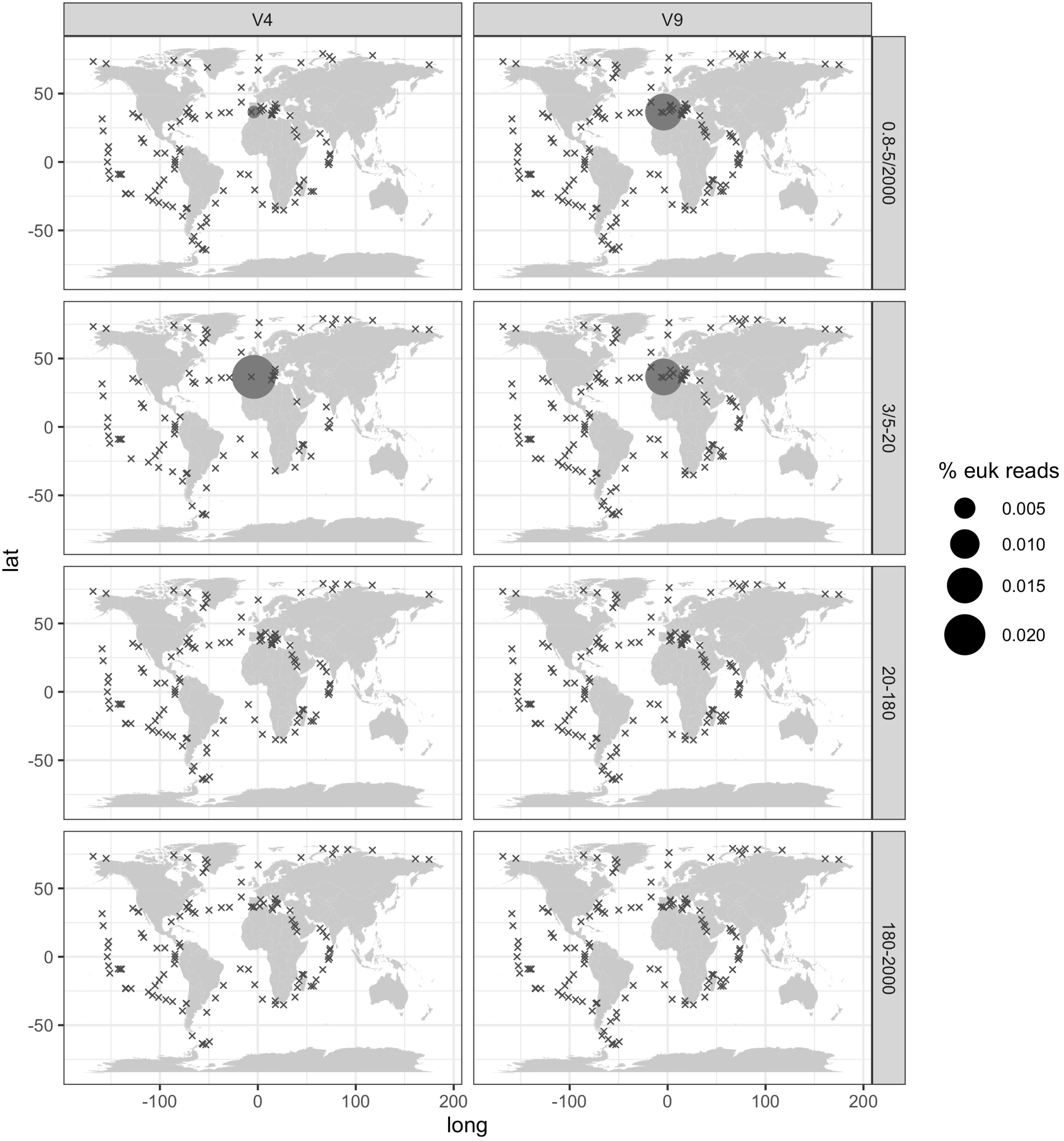

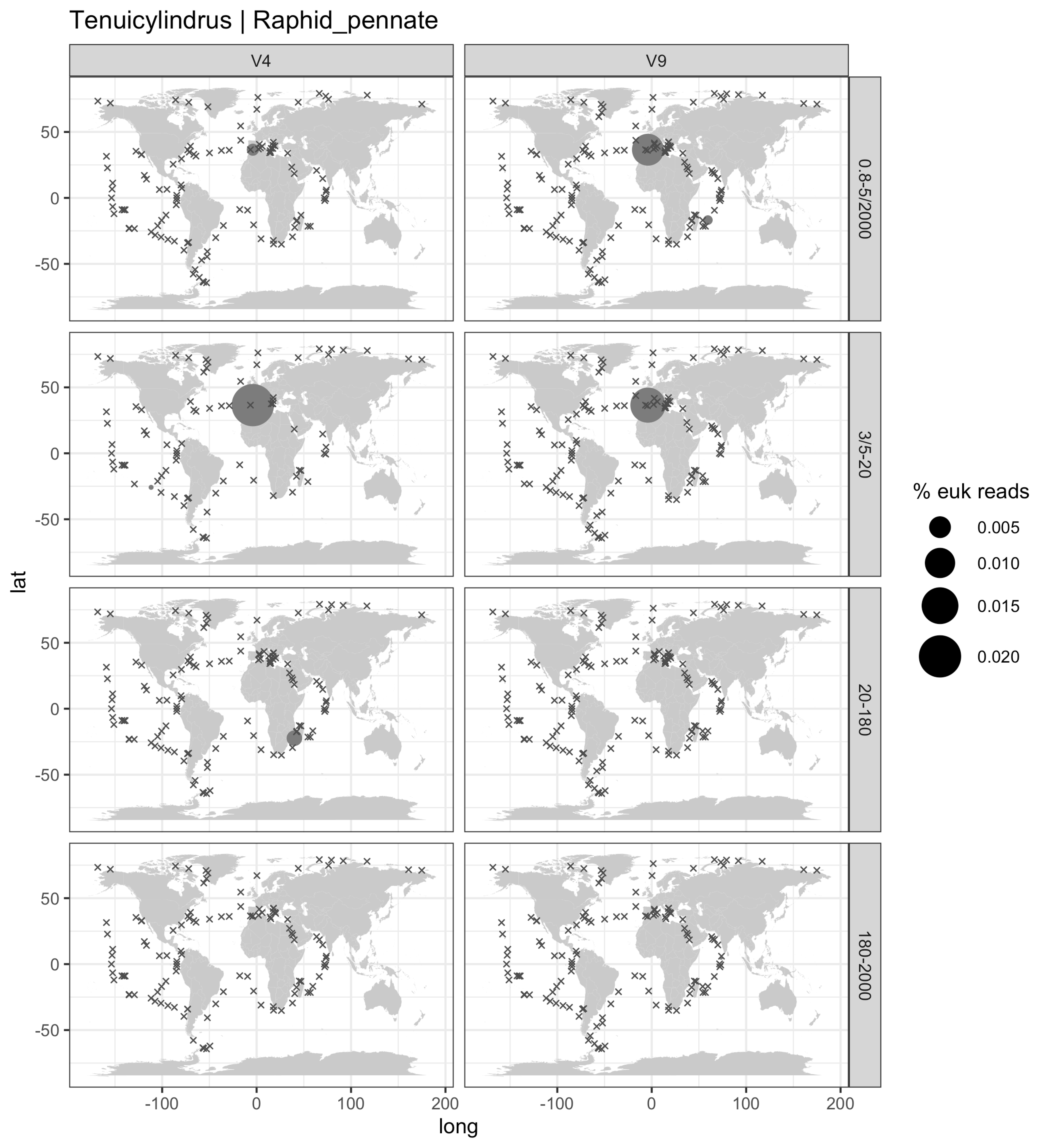

Tenuicylindrus | Raphid_pennate
V4
V9
0.8-5/2000
3/5-20
20-180
180-2000
lat
long
50
0
-50
-100
100
200
% euk reads
0.005
0.010
0.015
0.020

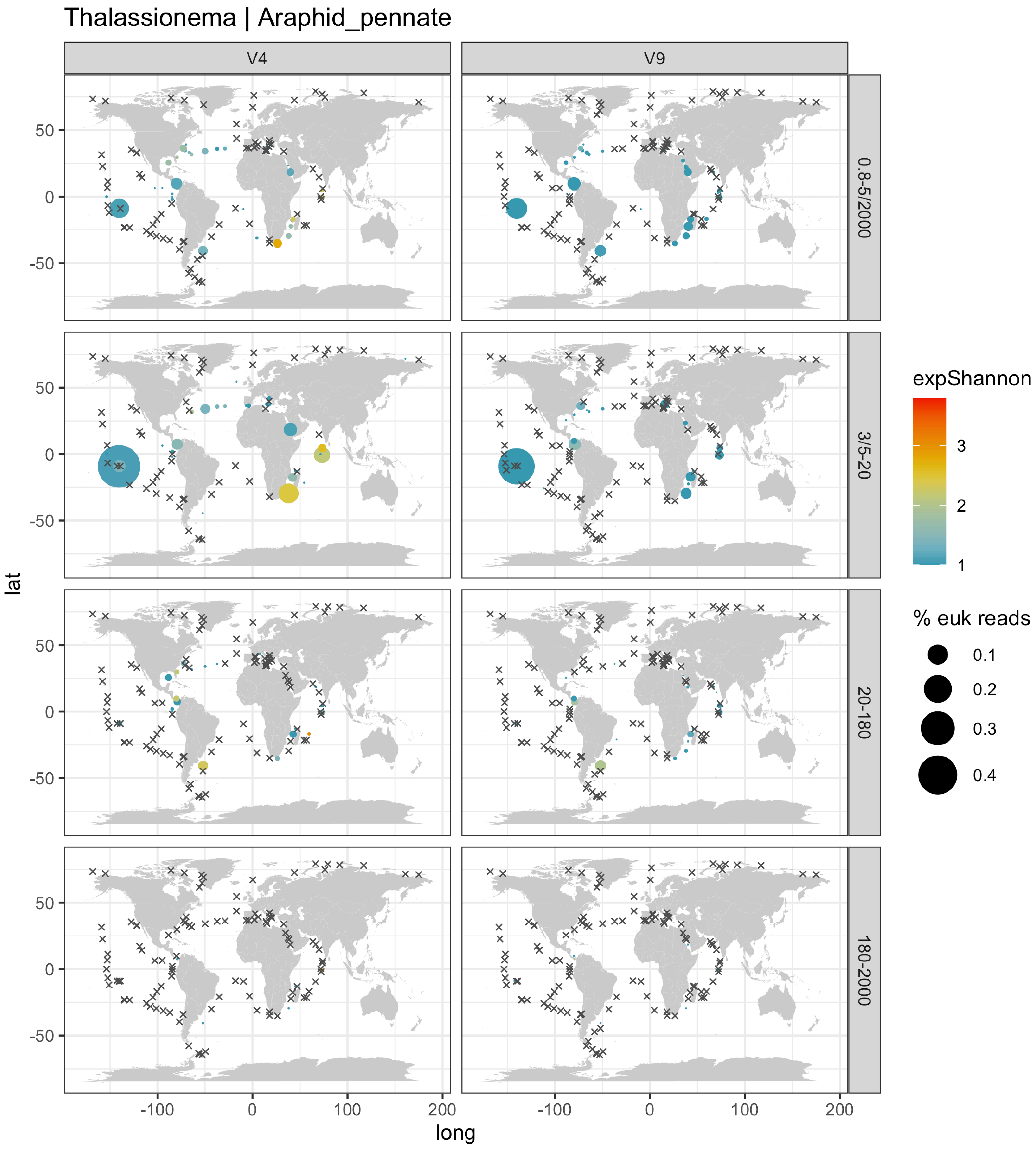

Thalassionema | Araphid_pennate
V4
V9
0.8-5/2000
3/5-20
20-180
180-2000
lat
long
50
0
-50
-100
100
200
expShannon
3
2
1
% euk reads
0.1
0.2
0.3
0.4

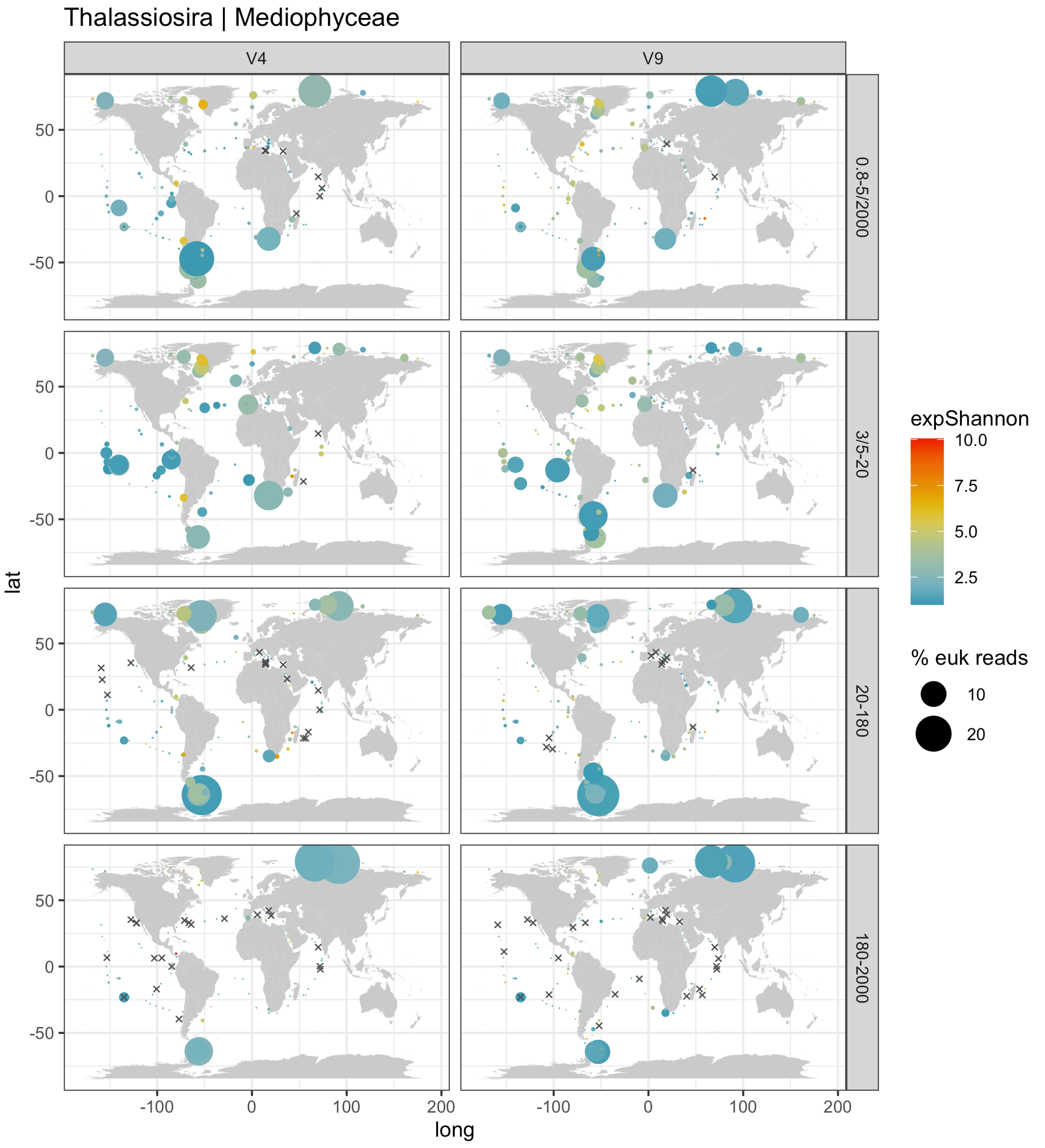

Thalassiosira | Mediophyceae
V4
V9
0.8-5/2000
3/5-20
20-180
180-2000
lat
long
50
0
-50
-100
0
100
200
expShannon
10.0
7.5
5.0
2.5
% euk reads
10
20

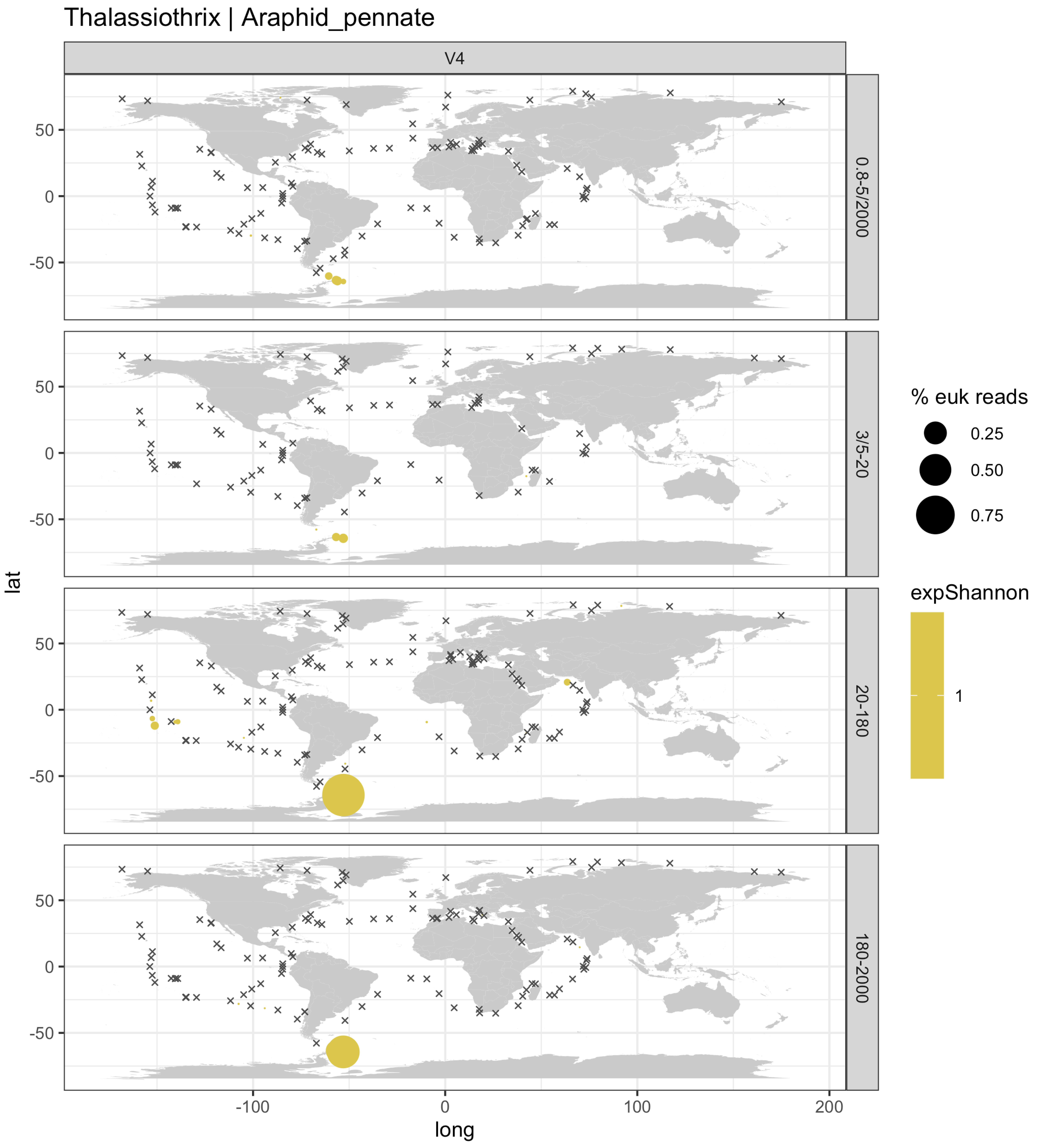
Thalassiothrix | Araphid_pennate
V4
0.8-5/2000
3/5-20
20-180
180-2000
lat
long
-100
0
100
200
50
0
-50
% euk reads
0.25
0.50
0.75
expShannon
1

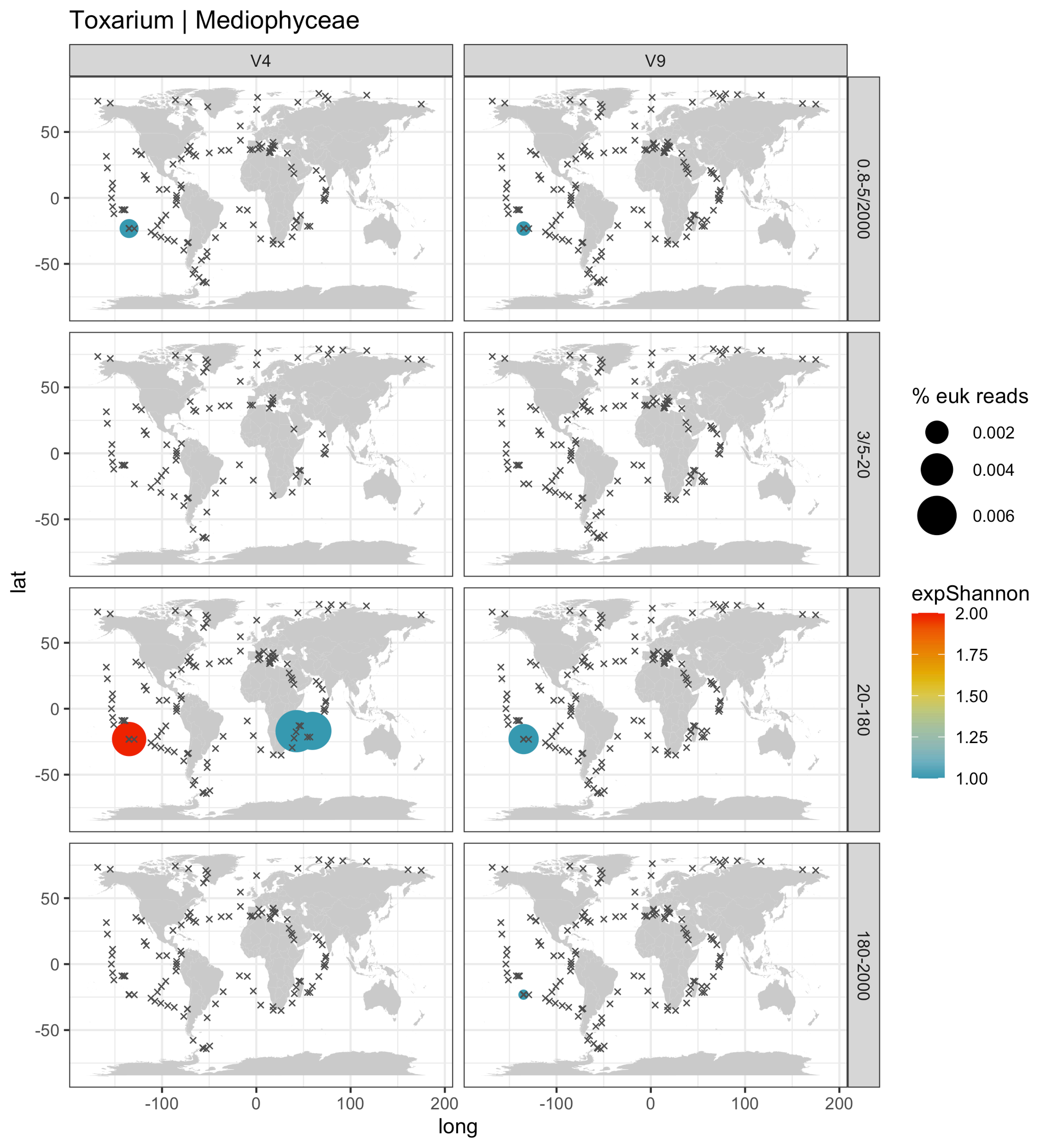

Toxarium | Mediophyceae
V4
V9
0.8-5/2000
3/5-20
20-180
180-2000
50
0
-50
-100
0
100
200
lat
long
% euk reads
0.002
0.004
0.006
expShannon
2.00
1.75
1.50
1.25
1.00

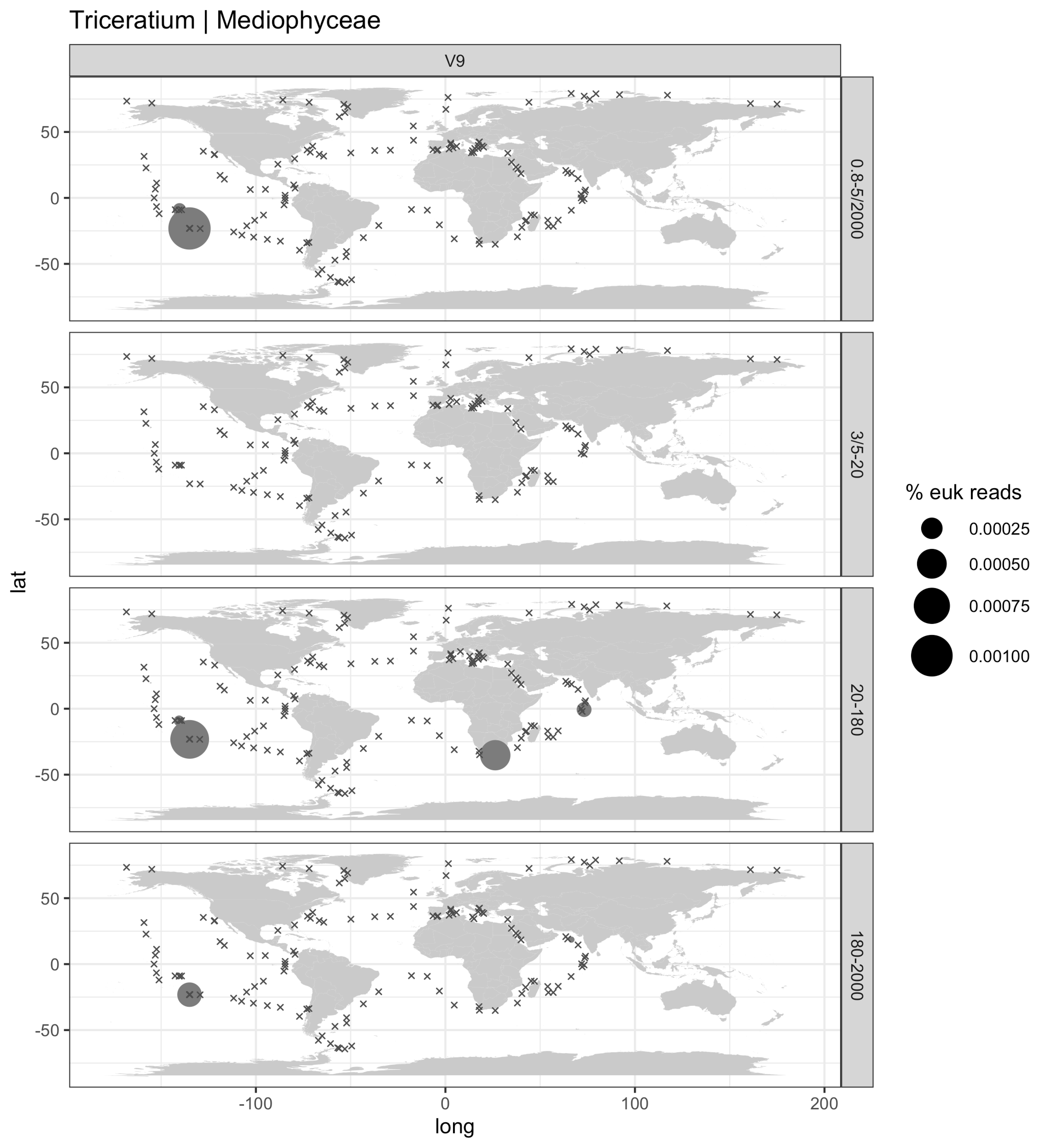

Triceratium | Mediophyceae
V9
0.8-5/2000
3/5-20
20-180
180-2000
lat
long
50
0
-50
-100
0
100
200
% euk reads
0.00025
0.00050
0.00075
0.00100

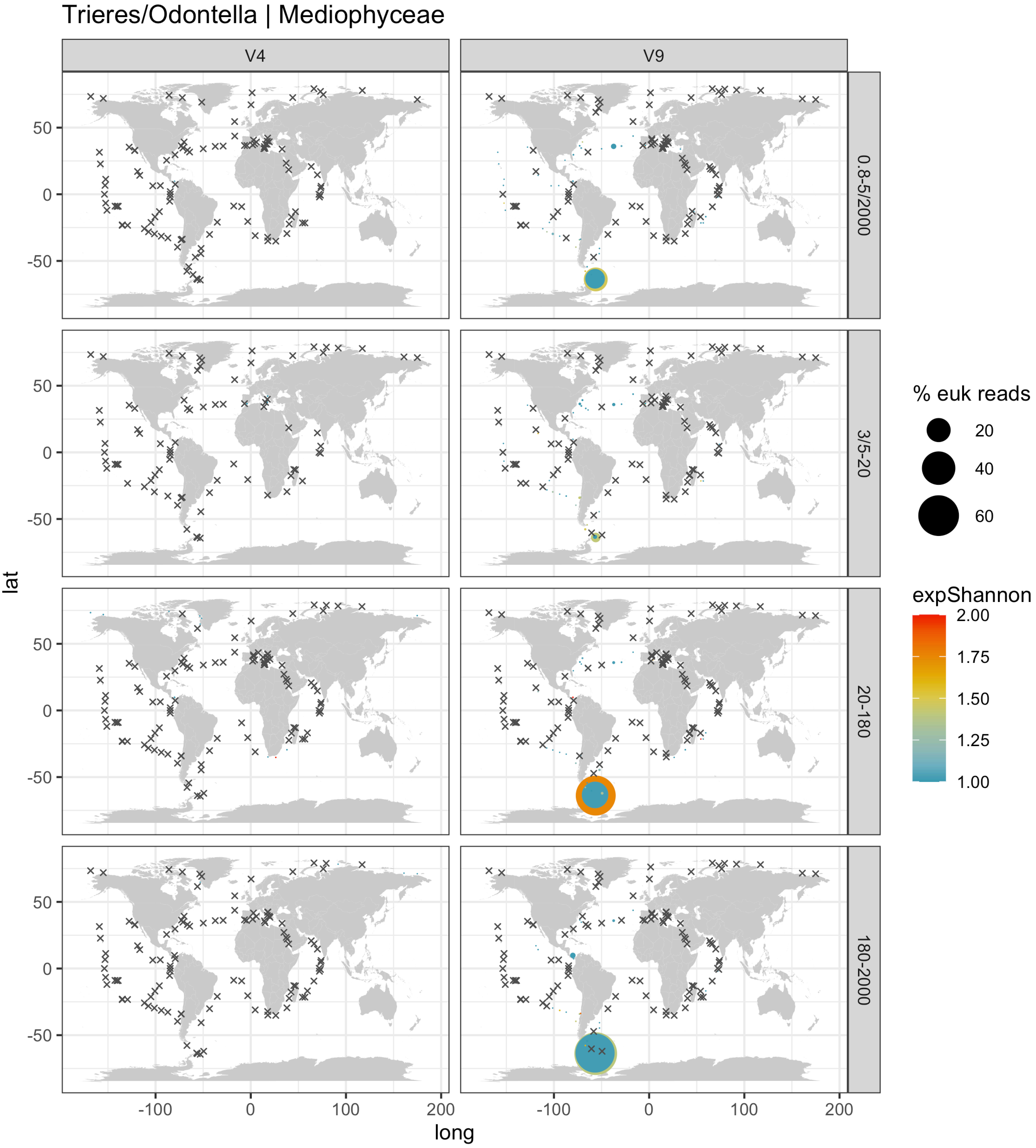
Trieres/Odontella | Mediophyceae
V4
V9
0.8-5/2000
3/5-20
20-180
180-2000
% euk reads
20
40
60
expShannon
2.00
1.75
1.50
1.25
1.00
lat
50
0
-50
long
-100
0
100
200
